# Progressive plasticity during colorectal cancer metastasis

**DOI:** 10.1101/2023.08.18.553925

**Authors:** AR Moorman, F Cambuli, EK Benitez, Q Jiang, Y Xie, A Mahmoud, M Lumish, S Hartner, S Balkaran, J Bermeo, S Asawa, C Firat, A Saxena, A Luthra, V Sgambati, K Luckett, F Wu, Y Li, Z Yi, I Masilionis, K Soares, E Pappou, R Yaeger, P Kingham, W Jarnagin, P Paty, MR Weiser, L Mazutis, M D’Angelica, J Shia, J Garcia-Aguilar, T Nawy, TJ Hollmann, R Chaligné, F Sanchez-Vega, R Sharma, D Pe’er, K Ganesh

## Abstract

Metastasis is the principal cause of cancer death, yet we lack an understanding of metastatic cell states, their relationship to primary tumor states, and the mechanisms by which they transition. In a cohort of biospecimen trios from same-patient normal colon, primary and metastatic colorectal cancer, we show that while primary tumors largely adopt LGR5^+^ intestinal stem-like states, metastases display progressive plasticity. Loss of intestinal cell states is accompanied by reprogramming into a highly conserved fetal progenitor state, followed by non-canonical differentiation into divergent squamous and neuroendocrine-like states, which is exacerbated by chemotherapy and associated with poor patient survival. Using matched patient-derived organoids, we demonstrate that metastatic cancer cells exhibit greater cell-autonomous multilineage differentiation potential in response to microenvironment cues than their intestinal lineage-restricted primary tumor counterparts. We identify PROX1 as a stabilizer of intestinal lineage in the fetal progenitor state, whose downregulation licenses non-canonical reprogramming.

## Introduction

As cancers progress, they become increasingly aggressive; metastatic tumors are less responsive to first-line therapies than primary tumors, they acquire resistance to successive therapies, and eventually lead to death^1–3^. Mutations are largely conserved between primary tumors and metastases resected from the same patients, suggesting that non-genetic phenotypic plasticity plays a major role in cancer progression and therapy resistance^4–7^. Since metastases are rarely resected surgically and epithelial cells are technically challenging to profile, most single-cell genomic studies of human cancer have either focused on primary tumors or immune cells from different tumor sites, while epithelial cell-state trajectories during tumor progression remain unresolved. Colorectal cancer (CRC), the second leading cause of cancer death worldwide, offers a unique opportunity to study cancer cell transitions during metastasis. Synchronous colorectal resection and metastasectomy, the simultaneous surgical resection of primary tumor, adjoining normal tissue and metastatic tumor (frequently from the liver), is a standard of care, providing access to patient-matched biospecimen trios^8^.

Colorectal cancer originates from Lgr5^+^ intestinal stem cells (ISCs) at the base of intestinal crypts^9^. During homeostasis, ISCs give rise to transit amplifying progenitors, which differentiate into distinct intestinal cell types^10^ Upon the loss of Lgr5^+^ ISCs by injury, multiple cell types can dedifferentiate into ISCs and subsequently regenerate intestinal crypts, demonstrating tremendous plasticity within the intestinal lineage^11^. ISCs that acquire oncogenic mutations serve as cancer stem cells that initiate canonical Wnt-driven intestinal adenomas and adenocarcinomas^12,13^. However, cancer cells that exist at the invasion front, enter the circulation, seed metastases, or regenerate tumors after therapy downregulate *LGR5* expression, and enter an L1CAM^+^ metastasis-initiating cell state resembling injury-associated regenerative progenitors^14^. We previously showed that entry into the L1CAM^+^ state is triggered by the disruption of adherens junctions between epithelial cells.

The phenotypic states of distant macrometastases that grow out from L1CAM+ metastasis-initiating cells remain unresolved. In mouse organoid transplantation models, Lgr5^-^ metastasis-initiating cells have been shown to re-enter Lgr5^+^ ISC states to regenerate liver metastasis^15–17^. However, the role of *LGR5*-dependent ISC-like states in metastasis is less clear in patients, where metastasis occurs over months or years and selects for cancer cells that must overcome numerous evolutionary stressors. Expression of *LGR5* and other WNT target genes is downregulated in liver metastasis^18^, while patient-derived CRC organoids transplanted into the mouse liver display variable *LGR5* expression and dependency following ablation of *LGR5*^+^ cells^19–21^. Since metastatic disease is the target of most cancer drugs, resolving the phenotypic states in metastatic CRC, their emergence from the primary tumor, plasticity and dependencies, is of paramount clinical importance.

To study metastatic progression in patients, we prospectively collected matched trios of normal colon, primary colon cancer and metastatic tissue from a cohort of 31 patients undergoing synchronous hemicolectomy and metastasectomy, including both treatment-naïve patients and those who received pre-operative chemotherapy. Using single-cell RNA-seq (scRNA-seq) of fresh sorted cells, multiplexed immunofluorescence, and epithelial organoid generation from matched trios, we investigated the evolution of cell states during metastatic progression, assessed the extent to which cell states and transitions are shared across patients or associated with therapy, and examined the roles of cell-autonomous plasticity and microenvironmental selection in progression.

We find that CRC progression involves three distinct, ordered cell-state transitions: (i) from differentiated intestinal lineages in normal colon to an LGR5^+^ ISC-like state in the primary tumor, (ii) developmental reprogramming to a conserved fetal progenitor-like state associated with epithelial injury during the transition to metastasis, and (iii) expression of gene programs associated with distinct non-intestinal lineages such as squamous, neuroendocrine and osteoblast, during outgrowth at the metastatic site. Organoids derived from a subset of profiled trios reveal that metastatic cells possess greater cell-intrinsic plasticity *in vitro* relative to primary tumor cells from the same patient, enabling dynamic adaptation to distinct microenvironments in the colon and liver *in vivo*. We find that the transcriptional repressor *PROX1* is coordinately induced with the fetal progenitor state across multiple patients, and functions to repress expression of non-intestinal lineage genes. Loss of PROX1-dependent lineage restriction during tumor progression licenses differentiation into non- canonical lineages. Together, our data support a two-stage model of metastatic plasticity; metastasis selects for highly plastic cell states that can be induced to differentiate along diverse trajectories by cues from the tumor microenvironment.

## Main

We prospectively collected fresh surgical tissue from primary CRC, adjacent normal colon and metastasis from 31 patients undergoing synchronous resection of colorectal tumor and metastasectomy at Memorial Sloan Kettering between February 2019 and September 2021 (**Fig. 1a–c** and **Extended Data Fig. 1** and **Supplementary Data 1**). Metastatic tissue was collected from the liver (29 samples), peritoneum, lung, and chest wall. Nine patients received no treatment prior to surgery, allowing us to observe the evolution to metastasis without chemotherapy as a confounder, while 22 patients received 5-fluorouracil-based chemotherapy prior to surgery. All patients had microsatellite stable, mismatch repair proficient (MSS/pMMR) metastatic colorectal cancer (see **Fig. 1b**,**c**, **Extended Data Fig. 1** and **Supplementary Data 1** for clinical, pathological and tumor genomic characteristics of each patient). For each tissue site, we performed scRNA- seq (83 samples), derived organoids from the single-cell suspension used for scRNA-seq (29 samples) and carried out multiplex immunofluorescence when tissue was available (72 samples). All patients were tracked longitudinally and additional surgical tissue was collected at a later time point from 6 patients who underwent a second, metachronous metastasectomy.

**Figure 1.**
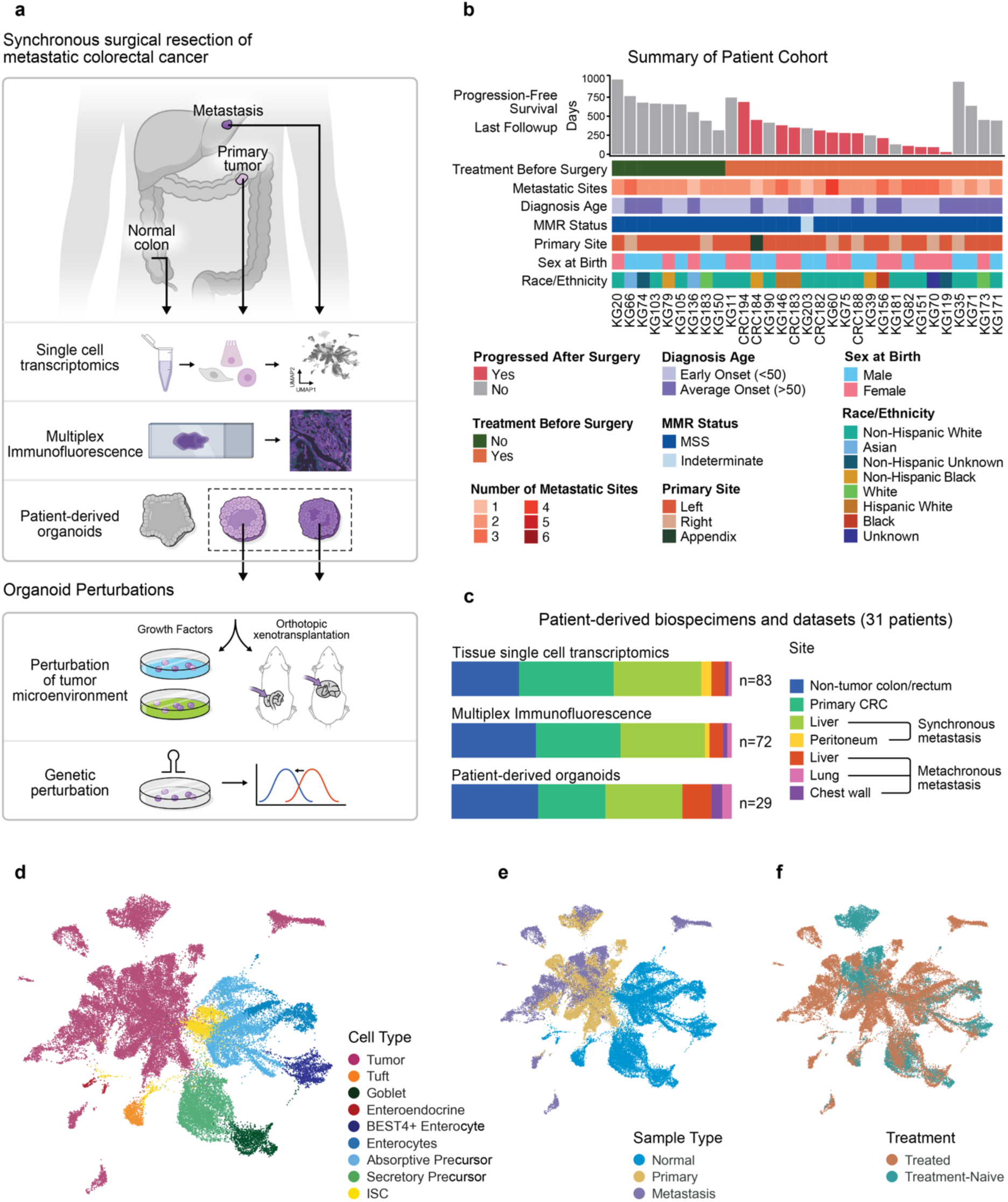
A patient-derived biospecimen platform to dissect transitions to the metastatic state. **a**, Study design. Matched trios of normal colon, primary colorectal cancer and metastasis were collected from 31 patients undergoing synchronous resection of the primary colorectal tumor and metastasectomy. Tissue was processed fresh for single-cell transcriptomics and organoid generation, and formalin-fixed and paraffin- embedded for multiplexed immunofluorescence assays. Patient-derived organoids were used for functional studies *in vitro* or via orthotopic xenotransplantation into the cecum or liver. **b**, Key clinical characteristics of each patient. **c**, Summary of biospecimens and datasets collected. **d**, Uniform manifold approximation and projection (UMAP) of all epithelial cell scRNA-seq profiles collected from non-tumor colon (20,817 cells), CRC primary tumor (14,402 cells), and CRC metastatic tumor (12,218 cells) samples, colored by cell type. ISC, Intestinal stem cell **e**, UMAP of epithelial cells, colored by tissue origin. **f**, UMAP of epithelial cells, colored by pre-surgical treatment status (see **Supplementary Data 1** for patient treatment details).

### A corrupted intestinal stem cell program is the strongest axis of variation in untreated colorectal cancer

We carried out scRNA-seq to identify shared mechanisms and bottlenecks in CRC progression across our patient cohort. Given the known sensitivity of epithelial cells to dissociation and their underrepresentation in single-cell CRC datasets, we optimized single-cell processing to maximize epithelial recovery, generating 47,437 high-quality epithelial cell profiles after initial data processing (Methods). We annotated epithelial cells using PhenoGraph^22^ to cluster cells and InferCNV^23^ to distinguish cancer from normal diploid cell copy number profiles, identifying intestinal stem cells (ISCs), differentiated secretory and absorptive cells, and tuft cells among other expected intestinal cell types (**Fig. 1d** and **Extended Data Fig. 2**). Moreover, cells from chemotherapy-treated and untreated normal colons from multiple patients cluster together, while primary and metastatic tumor cells are separate from normal cells (**Fig. 1e,f**). Primary tumor cells cluster more closely to one another and to normal epithelial cells, while metastases tend to occupy divergent and distant phenotypes (**Fig. 1e**).

The enrichment of epithelial cells in our atlas enabled us to explore transitions from normal colon to primary tumor, and from primary to metastasis. We first focused on samples from the 9 untreated patients whose tumor cells underwent transitions without the confounding influence of therapy (**Fig. 2a**). Using principal component analysis (PCA), we identified the strongest axis of variation in the data, and found that genes varying along the first (strongest) PC are significantly enriched for previously reported ISC signatures (FDR *q* < 0.04; **Supplementary Table 1f**)^24–26^. This agrees with observations that primary tumors, particularly those corresponding to consensus molecular subtype (CMS) 2, are enriched in *LGR5-*expressing cells^25,27,28^. In mice, the *Lgr5*+ ISC state is important for CRC initiation and macrometastatic outgrowth^13,16^ but is downregulated in metastasis-initiating cells^14,15,17^. We therefore sought to delineate the role of ISCs in cancer transitions represented in our unique trios of untreated patients with metastatic colorectal cancer. We first generated a *de novo* ISC signature based on genes that are differentially expressed in ISCs compared to differentiated intestinal cells in our untreated normal colon data (**Fig. 2b,c** and **Supplementary Table 1e**). Primary and metastatic tumors express high levels of the ISC signature relative to differentiated intestinal cell states, suggesting strong but incomplete selection for the ISC program in cancer (**Fig. 2d**). Genes associated with DNA synthesis and cell cycle progression are upregulated in normal ISCs relative to cancer cells (*RFC4*, *PRIM1*, *MCM2*) (**Extended Data** Fig 3a). On the other hand, many WNT-signaling associated genes are expressed at even higher levels in cancer cells than ISCs (*LGR5*, *EPHB2*, *ASCL2*, *TCF7*). In addition, cancer cells exhibit high expression of embryonic developmental genes that are lowly expressed in normal adult ISCs (*BMP7*, *SOX4*, *CYP2W1*). The reactivation of genes from more primitive stem cell programs suggests that cancer cells may inhabit a more dedifferentiated or embryonic state than ISCs.

**Figure 2.**
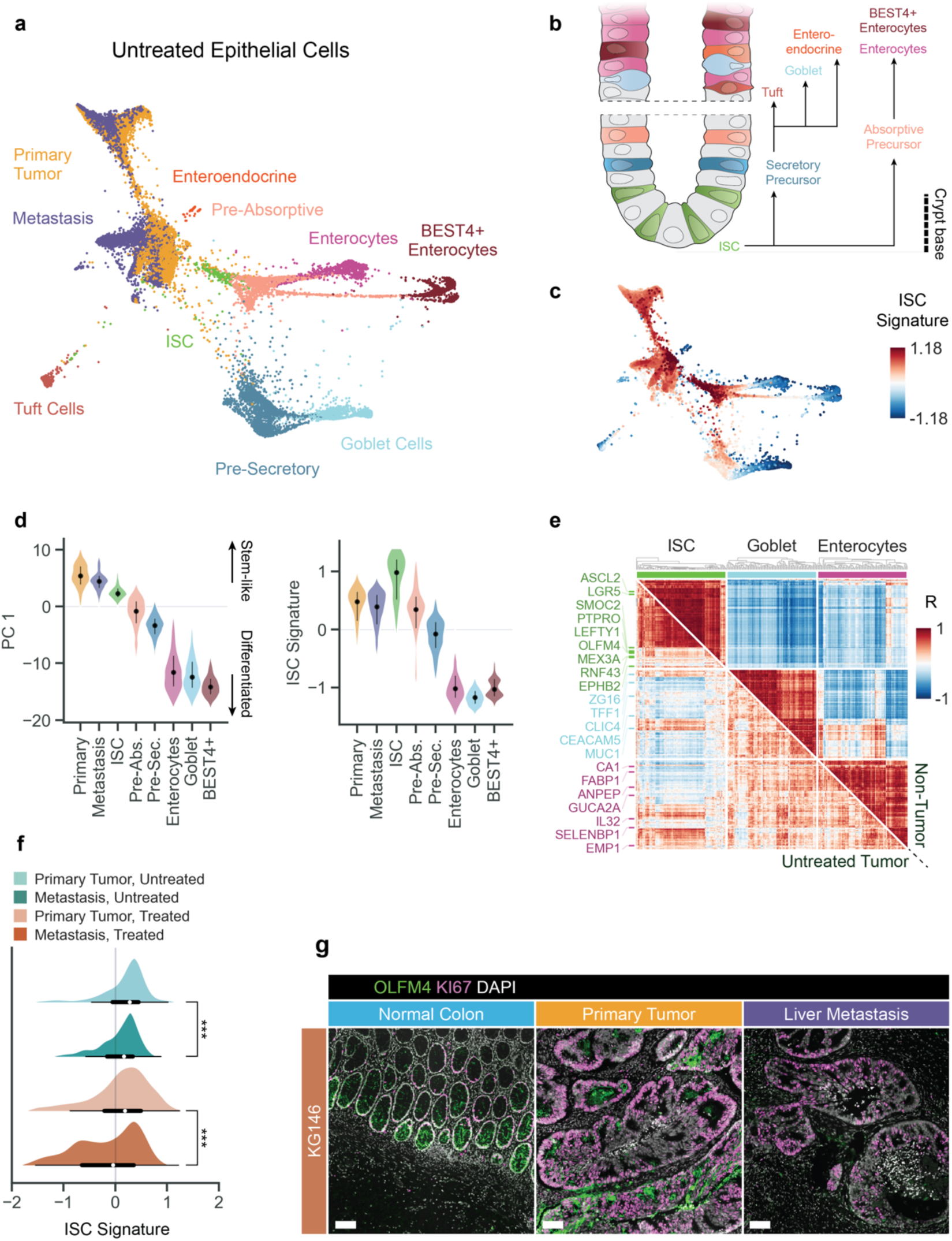
CRC primary and metastatic tumor cells display aberrant ISC and mixed lineage gene programs. **a**, Force-directed layout (FDL) of all treatment-naive epithelial cells (13,935 cells), colored by tumor type (primary, metastasis) for tumor cells or cell type for all other cells. **b**, Major cell types of the colon crypt. **c**, FDL of treatment-naive epithelial cells as in **a**, colored by ISC gene signature score (Methods). Scale indicates average, *z*-normalized gene expression (Methods). **d**, Distributions of first principal component (PC1) values (left) or relative ISC signature expression for all treatment-naive epithelial cell types, ordered by median PC1 value and colored as in **b**. Gene set enrichment analysis (GSEA) performed on genes ranked by their PC1 loadings was used to determine the more ‘differentiated’ versus ‘stem-like’ directions (Methods; see **Supplementary Table 1f** for all gene sets and enrichment scores). Dots, median values; lines, interquartile range. **e**, Gene-gene correlations in cells of the normal intestine (upper-diagonal matrix), and untreated tumors (lower-diagonal) showing expression of both ISC and mixed lineage markers in treatment-naive tumor cells. Genes (some highlighted) consist of the top 100 DEGs in ISCs, enterocytes, and goblet cells compared to all other non-tumor epithelial cells. Color values represent average gene-gene correlations calculated for each sample (Methods). **f**, Histograms of ISC gene signature scores for all tumor cells, indicating median (dots) and interquartile range (black bars). ***, p < 0.001; two-sided Wilcoxon rank-sum test. **g**, Vectra multiplex immunofluorescence from matched trios of normal colon, primary tumor and liver metastasis, showing OLFM4 (ISC marker), Ki67 and DAPI in representative samples from patient KG146 (scale bar = 100 μm).

To characterize cancer cells beyond the ISC signature, we searched for genes that are differentially expressed between cancer cells and normal ISCs (**Extended Data Fig. 3b**). Despite most closely resembling ISCs, we found that the untreated cancer cells also express gene programs associated with epithelial mesenchymal transition (EMT) and stress responses (unfolded protein response (UPR), nutrient stress, MTORC1) (**Extended Data Fig. 3b**). Normal ISCs, secretory goblet cells and absorptive enterocytes display strict cell-type-specific gene expression as expected (**Fig. 2e** and **Extended Data Fig. 3c**-e). In contrast, cancer cells co-express mixed intestinal cell-type programs for absorptive and secretory lineages with ISC-specific genes in the same cells (**Fig. 2e** and **Extended Data Fig. 3c**–e). This considerable lineage promiscuity indicates dysregulation of physiological intestinal hierarchies and gain of tumor-specific programs in colorectal cancer^29^, consistent with observations in lung adenocarcinoma, leukemia and others^22,30^. Also, in contrast to the ISC dependency described in mouse CRC metastases^16^, we find lower ISC program expression in metastases than primary tumors—a difference that is particularly notable in tumors that were resected following chemotherapy (**Fig. 2f,g**). Together, our patient data suggest that CRC tumors are enriched in ISC-like programs with primitive developmental and mixed lineage features, while metastatic progression appears to select against ISC programs.

### Non-canonical gene modules are enriched in metastasis

To delineate the transcriptional programs that define the transition from primary tumor to metastasis, we expanded our analysis to the entire diverse CRC cohort (**Supplementary Data 1**). Contrary to healthy intestine, cancer cells display substantial intra- and inter-patient phenotypic heterogeneity (**Fig. 1d,e**), thus, we aimed to find gene modules that are shared across patients and also capture variation within patients. Our assumption is that cells execute important biological functions by coordinating gene expression, which manifests as gene covariation in our data^31,32^. Moreover, to account for heterogeneity and pleiotropy, we wished to search for context-specific modules of genes that only co-vary within cell subsets^33^—particularly in cancer, where dysregulation can vary by patient, tumor site and other contexts. We therefore chose to use Hotspot^34^, which finds modules of genes that exhibit significant autocorrelation within local cellular neighborhoods of the phenotypic manifold. This approach is contrary to global measures such as Pearson correlation, which presume that relationships between features are consistent across a dataset^24,35^. Unlike other approaches, Hotspot does not entail full factorization of the data^32^.

To increase sensitivity, we applied Hotspot at fine resolution, using the k-nearest neighbor graph to define similarity between cells, and identified 37 gene programs across epithelial cells from all tumors, which we manually curated and annotated (**Supplementary Table 1k** and Methods). The fine resolution of our approach, coupled with the heterogeneity among patients and tissue contexts, led to redundancy between some modules. For example, Hotspot separated highly similar cell states from different patients into distinct modules. We therefore grouped modules based on their similarity and biological coherence (**Extended Data Fig. 4a**,b and Methods), highlighting ten annotated modules (**Fig. 3a** and **Extended Data Fig. 4a**) that are largely shared across multiple patients and are highly robust to parameter settings (**Extended Data Fig. 4c**,d). The local covariation employed by Hotspot was critical to identifying these modules, as these genes are not globally correlated (**Extended Data Fig. 4e**).

**Figure 3.**
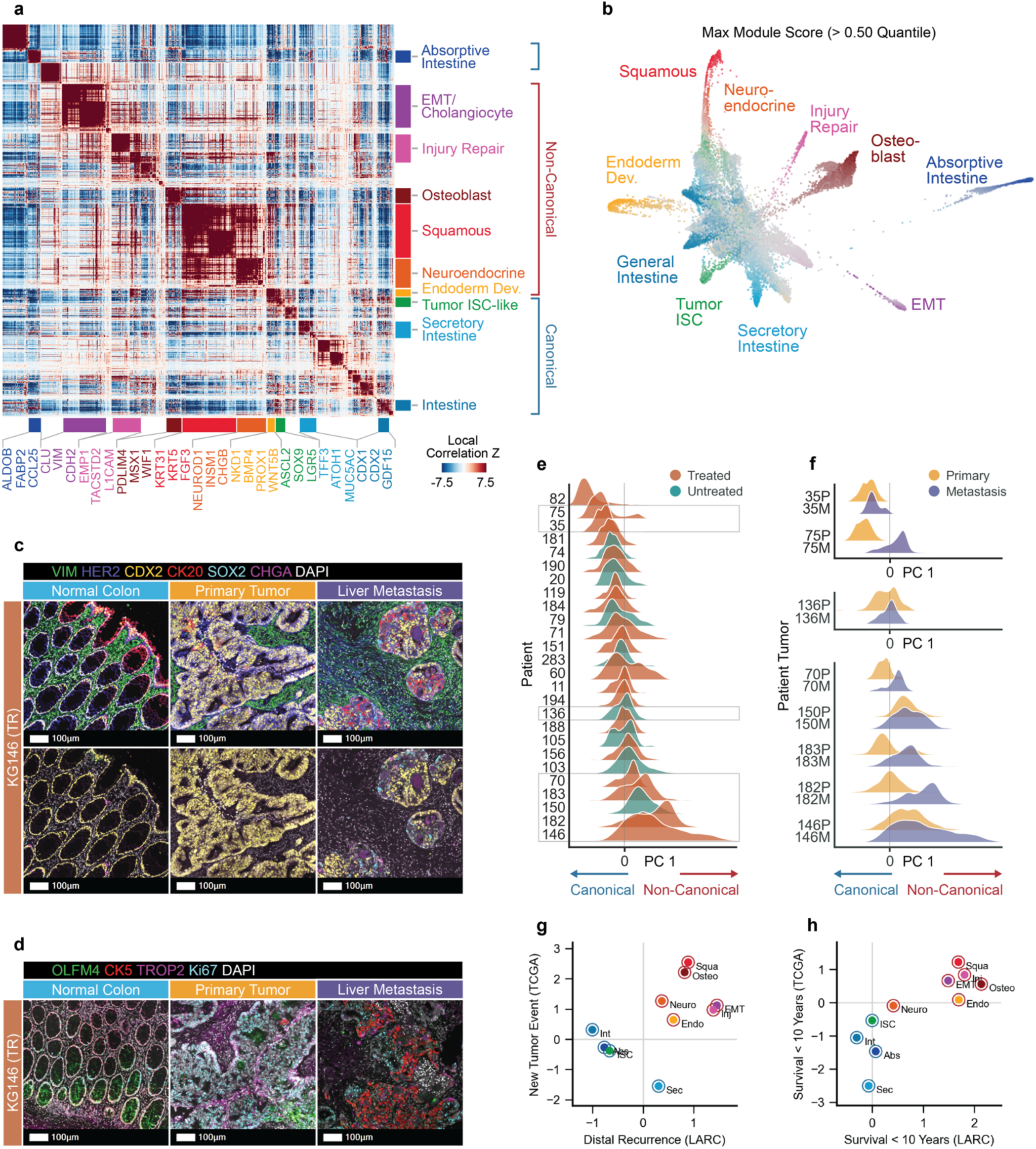
Non-canonical transcriptional programs in CRC are associated with metastasis and poor patient outcomes. **a**, Hotspot^34^ modules in all CRC tumor cells. Heatmap comprises 2003 highly variable genes with significant autocorrelation (FDR < 0.01), grouped into 10 CRC-derived gene modules (4 ‘canonical’ and 6 ‘non-canonical’ modules) (**Supplementary Table 1k** and Methods). **b**, FDL of all CRC tumor cells (26,145 cells), colored by gene module score for 10 CRC gene modules. Each cell is colored according to its maximum module score, or gray if no score exceeds the module’s 50th percentile across all cells. **c**, Vectra multiplex immunofluorescence from a representative matched trio of normal colon, primary tumor and liver metastasis from KG146) showing staining patterns for canonical intestinal and non-canonical genes: HER2 (pan- epithelial marker), CDX2 (intestinal lineage marker), CK20 (differentiated intestinal marker), SOX2 (multipotential stem cell marker), Chromogranin A (CHGA; neuroendocrine marker) and DAPI (nuclei), demonstrating decreasing order in morphology and increasing expression of non-canonical markers during tumor progression. **d**, Vectra multiplex immunofluorescence from the same patient (KG146) as in **c**, showing staining patterns for canonical intestinal and non-canonical genes: OLFM4 (ISC marker), CK5 (squamous cell marker), TROP2 (injury repair state marker), Ki67 (proliferation marker). **e**, Kernel density plot of tumor cells computed with *scipy* along a canonical: non-canonical axis (Methods) separated by patient and colored by treatment status. Patients with fewer than 50 cells are excluded. **f**, Kernel density plot depicting the distribution of tumor cells along the same canonical: non-canonical axis as in **e**, separated by primary (yellow) or metastatic (purple) tumors. **g**, Associations of module enrichments with survival status of patient donors in two independent clinical cohorts (Mann-Whitney U test). Associations are shown for 108 patients with rectal adenocarcinoma (LARC^46^) and for 445 patients with colon or rectal adenocarcinoma, published as part of The Cancer Genome Atlas (TCGA^45^). Enrichment scores are calculated using ssGSEA on bulk transcriptomic data from each patient (Methods). Each gene module is plotted according to the Mann Whitney U statistic of distal recurrence (x-axis) and new tumor event (y-axis). **h**, Same as **g** for associations of module enrichments with tumor recurrence.

We identified four tumor modules that express normal intestinal gene programs (**Fig. 2b**), including ISC-like (*LGR5*, *ASCL2*), differentiated absorptive (*FABP1*, *KRT20*) and secretory (*TFF1*, *LYZ*) states, which we term ‘canonical’ intestinal states. In addition, six ‘non-canonical’ modules express genes that are not expressed in the normal intestine, but exhibit tightly co-regulated expression corresponding to specific non-intestinal differentiated cell types (**Fig. 3a**). One module co-expresses *L1CAM* and *EMP1*, which have each been independently shown to mark *LGR5*^low^ CRC metastasis-initiating cells required for tumor regeneration^14,15^. This module contains additional genes associated with tissue regeneration after injury, drug-tolerant persister states and tumor progression, including *TACSTD2* (encoding cancer-associated trophoblast antigen TROP2)^36–38^, C*D70*^39^, and *OSMR*^40^. These observations support the emergence of a discrete metastasis- initiating cell population shown to have tumor regenerative properties, distinct from homeostatic ISCs or other epithelial cell types in advanced human cancer. We identified additional modules that cluster closely with the injury repair module, one expressing canonical markers of EMT (*CDH2*, *VIM*, *CLU*) and one expressing signaling genes associated with endodermal development (*WNT5B*, *BMP4*). Most strikingly, three modules express genes associated with differentiated non-intestinal cell states, representing squamous-like (*ELF5*, *SP5*), neuroendocrine-like (*NEUROD1*, *CHGB*) and osteoblast-like (*MSX1*, *DLX5*) lineages. While the normal colon contains a small population of enteroendocrine cells, differences in gene expression (for example, the tumor module lacks intestinal transcription factors *CDX1* and *CDX2*) and prevalence in cancer cells prompted us to classify the tumor neuroendocrine population as non-canonical.

Mapping each cell to its highest scoring module revealed that most modules reside at phenotypic extremities, which correspond to archetypes in this data (**Fig. 3b**). However, most cells express multiple modules— commonly combinations of tumor ISC-like, injury repair, EMT and endoderm modules—whereas more differentiated modules such as absorptive intestine tend to be more uniquely expressed in a cell (**Extended Data Fig. 4f**,g). We noted that most cells express canonical or non-canonical modules exclusively, except for a population that expresses both tumor ISC-like and endoderm modules (**Extended Data Fig. 4f**–h). Given the distinction between canonical and non-canonical modules, we wanted to visualize these spatially and validate non-canonical marker expression at the protein level. Therefore, we performed multiplex immunofluorescence on 73 tissue sections, including 65 derived from part of the same tissue samples used for scRNA-seq (**Fig. 3c,d**). While primary tumors display multilayered, dysplastic crypt architecture, they retain expression of the intestinal lineage-defining transcription factor CDX2^41,42^. In matched, synchronously resected metastases that express non-canonical modules in scRNA-seq data, expression of CDX2 and the ISC marker OLFM4 is downregulated, accompanied by higher expression of non-canonical markers including SOX2 (stemness), CHGA (neuroendocrine-like), CK5 (squamous-like) and TROP2 (encoded by *TACSTD2*, injury repair).

### Non-canonical gene expression varies across patients and is associated with poor clinical outcomes

Next, we sought to characterize patterns of canonical and non-canonical module expression across patients. We found that the Hotspot gene modules are expressed in multiple patients, and importantly, that expression of non-canonical modules occurs repeatedly in different metastasizing cancers (**Extended Data Fig. 4i**); moreover, the strongest principal component of variation in the full tumor cell dataset is closely associated with the transition from canonical to non-canonical module expression (**Fig. 3e,f**). Cells from some patients (eg. 82, 75, 35) skew largely canonical, while others (e.g. 183, 150, 182, 146) span the full spectrum of phenotypic states (**Fig. 3e**). However, in every primary–metastasis pair from individual patients, the metastatic tumor displays greater non-canonical expression than the simultaneously resected primary tumor (**Fig. 3f**). Importantly, we observed neuroendocrine-like and squamous-like gene expression in metastases of some untreated patients (e.g. 150), suggesting that metastasis *per se*, rather than therapy, is sufficient to induce entry into non-canonical phenotypic states. Together, these data demonstrate the progressive loss of canonical intestinal lineage identity, and the gain of non-canonical gene expression in the transition to CRC metastasis.

Lineage plasticity has previously been associated with therapy-resistant advanced disease in castrate- resistant prostate cancer and lung adenocarcinoma^43^, but has not been described as a feature of metastasis in the absence of selective pressure from therapy (**Fig. 3e**) or recognized as a feature of colorectal cancer. We asked whether canonical or non-canonical gene expression is associated with clinical or pathological variables in our fully annotated cohort of 31 patients with metastatic CRC, and found that non-canonical module expression is associated with metastasis and prior chemotherapy (**Fig. 3e,f** and **Extended Data Fig. 5a**–c). as well as other biomarkers of more aggressive disease including progression after surgery and poor overall survival (**Extended Data Fig. 5c**). Bulk RNA-seq has been used to classify CRC into CMSs, with CMS2 (representing high WNT signaling) associated with good outcomes and CMS4 (representing high mesenchymal gene expression) with poor outcomes^27^. Expression of our single-cell transcriptomics-based canonical modules corresponds with CMS2, while differentiated intestinal secretory module expression corresponds with CMS3 (**Extended Data Fig. 5d**). In contrast, diverse non-canonical modules correspond with CMS4.

To determine whether non-canonical module expression in untreated, non-metastatic primary tumors could serve as a biomarker of clinical outcome, we performed single sample gene set enrichment analysis (ssGSEA^44^) on two independent datasets of pre-treatment primary tumors, the cancer genome atlas (TCGA) cohort of 445 patients with Stage I-IV colon or rectal adenocarcinoma^45^, and an MSKCC cohort of 108 patients with locally advanced (Stage II-III) rectal cancers (LARC)^46^. Expression of non-canonical modules is associated with poor outcomes in both cohorts, including tumor relapse after surgery and overall survival <10 years (**Fig. 3g,h** and **Extended Data Fig. 5e**–g). Together, our analyses suggest that non-canonical module expression can exist in subpopulations within untreated primary tumors and undergo positive selection during metastasis, and that it is associated with negative clinical outcomes including poor overall survival.

### CRC undergoes stereotyped, sequential fetal and non-canonical cell state transitions during metastatic progression

We next sought to understand how the transition from canonical to non-canonical states occurs, using our matched samples that span metastatic progression in the same patient. We first focused on the patient whose tumor cells display the greatest phenotypic heterogeneity (KG146), including all canonical and non-canonical states, as well as the most differentiated non-canonical states (**Figs. 3e,4a**). As with PCA, we found that the first diffusion component (DC1) in this data corresponds to the canonical-to-non-canonical transition, revealing a progression from differentiated absorptive and secretory intestinal lineages in the primary tumor to an ISC state and subsequently to non-canonical endodermal development, neuroendocrine-like and squamous-like states in metastasis (**Fig. 4a,b**).

**Figure 4.**
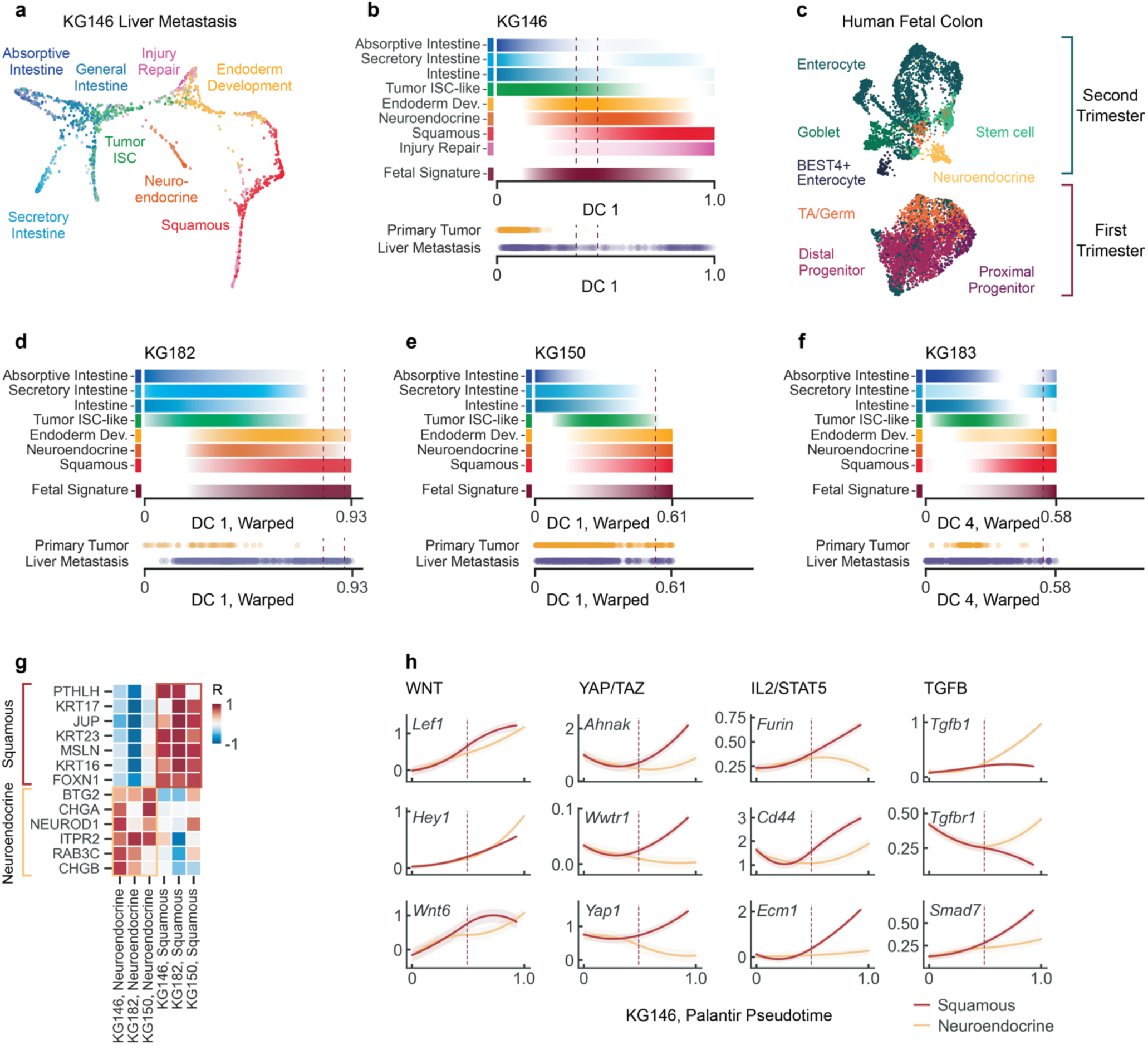
A conserved fetal progenitor intermediate bridges canonical and noncanonical states in CRC. **a**, FDL of cells from KG146 liver metastasis (1,279 cells), showing a diversity of canonical and non-canonical states. Each cell is colored according to its maximum module score, or gray if no module score exceeds its 25th percentile across all cells. **b**, Top, trends in gene module scores along DC1, observed in all tumor cells from patient KG146 (Methods). Each row depicts the module score along DC1 from the 20th percentile value (white) to the maximum value (highest saturation). Expression of the fetal signature peaks between canonical and non-canonical trends (vertical dashed lines correspond to 90th percentile). Bottom, positions of tumor cells along DC1. **c**, UMAP embedding of human fetal colon cells^50^. Cells are colored by cell type according to annotations in the original publication and organized as first trimester (6-11 post-conception wks, PCW) or second-trimester (12-17 PCW). **d–f**, Top, gene module scores as in **b** along DC1, observed in all tumor cells from patient KG182 (**d**), patient KG150 (**e**) and patient KG183 (**f**). DC values were first aligned to DC1 from patient KG146 using dynamic time warping (**Extended Data Fig. 6b**–e and Methods). Bottom, positions of tumor cells along DC1. **g**, Pearson correlations of Palantir^54^ branch probabilities with gene expression for genes associated with squamous (top) and neuroendocrine (bottom) transdifferentiation, across tumors from patients KG146, KG182 and KG150. **h**, Gene expression trends along Palantir pseudotime for selected signaling pathways during trans-differentiation of KG146 tumor cells toward squamous or neuroendocrine- like terminal states. Trends are computed using generalized additive models (GAMs) in Palantir. Solid lines represent mean expression, shaded regions represent 1 s.d.

The position of the endoderm development module as an intermediate state bridging canonical intestinal and non-canonical states caught our attention, since transient reversion to more developmentally primitive states has been associated with regeneration following intestinal epithelial injury^36,47,48^ as well as tumor biology^30,49^. To characterize the endodermal state systematically, we turned to a fetal development dataset from the Human Cell Atlas^50,51^. In the first trimester, colon epithelial cells assume an undifferentiated ‘fetal progenitor- like’ state, while by the second trimester, major organogenesis is largely complete and ISC and differentiated intestinal states can be clearly identified (**Fig. 4c**). We identified genes that are differentially expressed in first- trimester progenitors relative to second-trimester mature colonocytes in the developmental atlas (**Supplementary Table 1m** and Methods) and plotted expression of this fetal progenitor signature along DC1, finding that it marks a clear intermediate state between canonical intestinal and non-canonical differentiated states (**Fig. 4b**).

Next, we extended our analysis to the patients with the most heterogeneous tumors (KG146, KG182, KG150, KG183) and identified many fetal signature genes that are highly upregulated in a majority of these tumors (**Supplementary Table 1n** and Methods), including WNT signaling (e.g. *RBX1*, *PTK7*, *CTNNB1*), embryonic and neuronal development (e.g. *SOX4*, *BEX1*, *RARRES2*, *LMO4*), and adhesion (e.g. *MFGE8*, *BCAM*) genes. Comparing first-trimester colon cells to cancer cells that express the fetal signature, the most upregulated genes are, surprisingly, markers of intestinal lineage (e.g. *CDX1*, *LGR4*, *AGR2*, *HMGCS2*). Thus, our data suggests that cancer progression involves developmental reversion, characterized by a more primitive dedifferentiated state than that observed in first trimester colonic progenitors, with profound loss of intestinal lineage identity and upregulation of early developmental programs associated with WNT signaling.

Determining the extent to which this sequential cell-state trajectory exists across multiple patients presents multiple challenges, given the substantial inter-patient heterogeneity. First, patients exhibit different ranges of canonical and noncanonical states (**Fig. 3e**), requiring trajectory alignment across different start and end points. Second, alignment needs to account for differences in cell-state frequencies between patients. We thus used dynamic time warping^52^ a method that preserves cell order within each trajectory while allowing for different start points, end points and progression rates (**Extended Data Fig. 6a**), and that has been used to align single-cell trajectories while accounting for inter-individual variability^53–55^. We used dynamic time warping to align trajectories from each of the four patients whose tumors exhibit non-canonical differentiation to that of KG146 (possessing the broadest range of cell states) and ensured the robustness of these alignments (**Extended Data Fig. 6b**,c). Although tumors progress towards non-canonical states to different extents, we identified a common progression sequence from differentiated intestinal to ISC, fetal progenitor, and finally differentiated squamous or neuroendocrine-like states (**Fig. 4b,d–f**). Fetal progenitor signature expression increases as intestinal module expression declines, suggesting that a fetal-like intermediate state accompanies the loss of intestinal identity and precedes non-intestinal transdifferentiation (**Fig. 4b,d–f**).

To determine whether tumor evolution upon additional time or therapy leads to greater non-canonical differentiation, we analyzed secondary metastases collected from patients several months after the original resection and subsequent chemotherapy (**Extended Data Fig. 1a** and **Supplementary Data 1**). Indeed, secondary liver (KG185) and chest wall (KG182) metastases exhibit further progression towards non- canonical states along the DC (**Extended Data Fig. 6d**). A secondary lung metastasis (KG146) displays more non-canonical expression than the primary tumor, but less than the original liver metastasis, consistent with the typically more benign clinical course of CRC lung metastases relative to liver^56,57^.

Both squamous and neuroendocrine-like states are present in every tumor with non-canonical gene expression. To identify possible drivers of these divergent fates, we employed Palantir trajectory inference^54^, which confirmed the bifurcation of squamous and neuroendocrine trajectories and placed the fetal signature between branches (**Fig. 4g** and **Extended Data Fig. 6e**). Examining gene expression in branches along the canonical-to-non-canonical axis, we found that WNT signaling genes are coordinately upregulated in both branches, whereas squamous differentiation correlates with YAP and IL-2 signaling, and neuroendocrine differentiation correlates with TGF-β signaling, including the upregulation of TGF-β1 and TGF-βR1 and downregulation of TGF-β inhibitor SMAD7 (**Fig. 4h**).

Together, our data reveal a recurrent program of sequential cell-state transitions during CRC progression; canonical intestinal-like states are progressively lost as cancer cells revert to a fetal progenitor state, and then differentiate into non-canonical squamous-like and neuroendocrine-like states. In both the LARC and TCGA cohorts, high expression of the fetal progenitor signature in pre-treatment primary tumors is in fact associated with decreased disease-free survival, demonstrating its potential utility as a biomarker of clinical outcome (**Extended Data Fig. 6f**,g). Despite inter- and intra-tumoral heterogeneity within and across patients, the fetal progenitor state appears to serve as a convergent tumor-regenerative intermediate, which bridges canonical states with the aggressive non-canonical states associated with poor clinical outcomes.

### Cancer cell-intrinsic and extrinsic factors induce the emergence of non-canonical states during metastasis and therapy

While we observed increasing non-canonical differentiation during CRC progression, we do not know whether it is driven by cancer cell-autonomous changes during metastasis and therapy, or by differences between metastatic and colonic microenvironments. To address the mechanisms underlying phenotypic plasticity, we generated organoid models from the same tissue suspensions used for single-cell analysis. MSK-IMPACT sequencing confirmed that mutations are conserved between the patient tumor and both primary and metastasis-derived organoids (**Extended Data Fig. 7a**). We first focused on organoids derived from patient KG146 primary tumor (OKG146P) and liver metastasis (OKG146Li), as the originating primary tumor comprises largely canonical intestinal states (with a small proportion of cells in the injury repair state), while the metastasis spans the full spectrum of canonical and non-canonical states (**Fig. 4b** and **Extended Data Fig. 7b**, c). To control for microenvironment, we initially grew cancer cells in standard human intestinal stem cell (HISC) media, which contains intestine-specific niche factors that sustain ISCs (WNT, R-spondin, EGF, noggin, A-8301, FGF and IGF1) (**Fig. 5a**).

**Figure 5.**
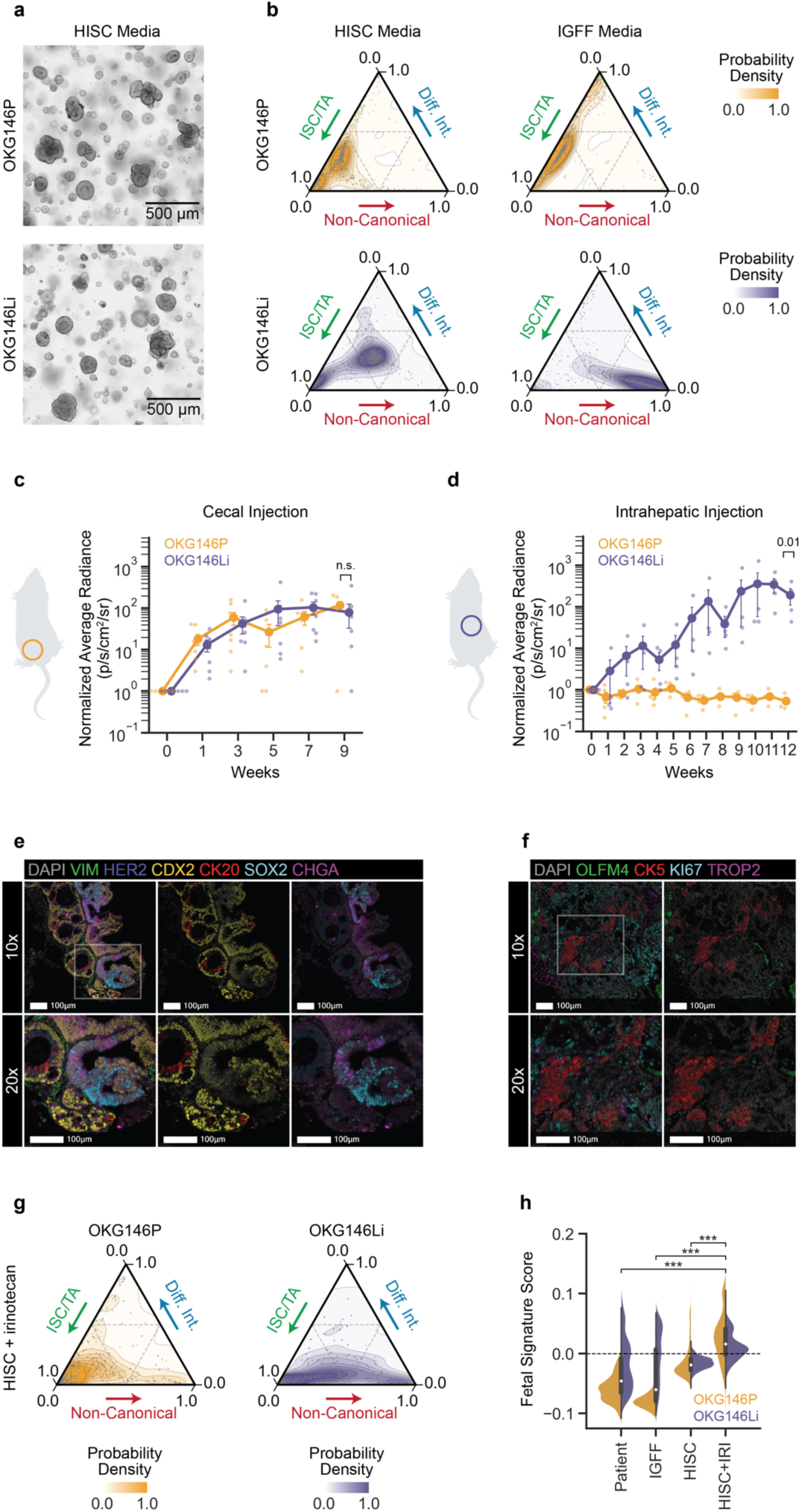
Organoid models reveal distinct contributions of tumor and microenvironment to metastatic plasticity. **a**, Brightfield microscopy showing morphology of paired primary rectal tumor (OKG146P) and liver metastasis (OKG146Li)-derived organoids, 7 days following seeding as single cells (2,000 cells/40 µl Matrigel). **b**, Cell state assignment probabilities per cell, calculated as Markov absorption probabilities (Methods), for OKG146P and OKG146Li organoids grown in human ISC (HISC) and intestinal growth factor free (IGFF) media. Lines indicate density contours within 5th and 95th percentiles of each distribution, dots indicate individual cells. TA, transit amplifying cells. Diff. Int., differentiated intestinal states. **c**,**d**, Normalized average radiance measured by weekly *ex vivo* bioluminescence imaging following cecal injection of 200,000 organoids (**c**) or intrahepatic injection of 500,000 organoids (**d**) from KG146P (primary tumor) and OKG146Li (metastasis) lines in NSG mice, normalized to signal immediately following injection (week 0). Error bars, s.e.m, n = 7 and 4; *p* = 0.2246 and 0.0143 between organoids in **c** and **d** respectively, Mann Whitney rank-sum test. **e**, Representative Vectra images of OKG146Li intrahepatic xenografts, harvested at 13 weeks post injection (wpi), stained for DAPI (nuclei), VIM (vimentin, stroma) HER2 (epithelial differentiation), CDX2 (intestine), CK20 (differentiated intestine), SOX2 (stem cell) and CHGA (neuroendocrine) markers. **f**, Representative Vectra images of OKG146Li intrahepatic xenografts, harvested at 13 wpi, stained for DAPI (nuclei), OLFM4 (ISC), CK5 (squamous), Ki67 (proliferation) and TROP2 (injury repair state) markers. **g**, Cell state assignment probabilities per cell for live cells from OKG146 organoids grown in HISC media and treated with 250 nM irinotecan chemotherapy for 7 days. Lines indicate density contours within 5th and 95th percentiles of each distribution, dots indicate individual cells. **h**, Fetal signature score distributions in primary (orange) and metastatic (purple) cancer cells from patient KG146 in patient tumor (left) and patient-derived organoids cultured in IGFF, HISC or HISC media containing 250 nM irinotecan. Fetal signature is the same as in Fig. 4 and calculated on z-normalized, imputed expression. ***, p < 0.001, two-sided t-tests comparing primary and metastasis samples between conditions.

Using a mutually nearest neighbor approach to map organoid phenotypic states to states in the tumors of origin (**Extended Data Fig. 7b**,c and Methods), OKG146P cells retain the largely ISC-like phenotype present in KG146P *in vivo* (**Fig. 5b** and **Extended Data Fig. 7d**). In contrast, OKG146Li cells adopt ISC-like and endodermal progenitor-like states when grown in HISC (**Fig. 5b** and **Extended Data Fig. 7d**), but demonstrate notably little of the non-canonical gene expression observed *in vivo* in KG146Li (**Figs. 5b** and **Extended Data Fig. 7c**,d). We therefore asked whether growth factors in the ISC medium might inhibit differentiation into non-canonical lineages. Indeed, withdrawing ISC niche factors from the medium causes OKG146Li cells to lose intestinal gene expression and gain non-canonical expression associated with injury repair and squamous-like, osteoblast-like and neuroendocrine-like lineages, whereas OKG146P cells retain intestinal epithelial gene expression—albeit less ISC-like and with higher levels of intestinal absorptive and secretory differentiation markers (**Fig. 5b** and **Extended Data Fig. 7d**). In liver metastases from two other patients (OKG182CW2, OKG183Li2), switching from HISC to intestinal growth factor-free (IGFF) medium likewise decreases expression of ISC marker *LGR5* and increases expression of non-canonical and some canonical differentiation markers (**Extended Data Fig. 7e**,f). Cancer cells from primary tumors thus appear to be largely restricted to an axis from ISC to differentiated intestinal states, whereas metastatic cells acquire greater cell-autonomous plasticity, including the ability to upregulate gene expression for a variety of non-canonical lineages in response to local microenvironmental signals.

To determine whether different intrinsic abilities to adapt to colon and liver microenvironments are retained *in vivo*, we xenografted organoids into the cecum (intestinal microenvironment) of NOD scid gamma (NSG) mice. Both primary- and metastasis-derived organoids grew into orthotopic cecal tumors, with similar kinetics (**Fig. 5c**); however, OKG146P organoids failed to grow out in the liver following intrahepatic injection, while OKG146Li organoids readily adapted to the liver microenvironment (**Fig. 5d**). Using Vectra multiplex immunofluorescence, we found that OKG146Li-derived xenografts retained the ability to differentiate into the full spectrum of canonical and non-canonical states (**Fig. 5e,f**), similar to the liver metastases in patients (**Fig. 3c,d**). Within the tumors, distinct individual gland-like structures adjoining one another either express high levels of canonical intestinal markers (HER2, CDX2, CK20, OLFM4), or high levels of non-canonical markers SOX2 (stem), Chromogranin A (neuroendocrine) and CK5 (squamous) with low intestinal lineage marker expression. Expression of canonical and non-canonical markers appears to be mutually exclusive at the level of the gland, suggesting the ability of individual metastasis-initiating cells to stochastically enter into either a canonical or non-canonical cell fate early during metastatic outgrowth.

To interrogate the relationship between therapy and non-canonical gene expression, we treated OKG146P and OKG46Li organoids grown in HISC with their IC50 dose of the first-line chemotherapeutic for metastatic CRC, irinotecan, and performed scRNA-seq on sorted, DAPI-viable residual cells after 7 days. Chemotherapy induced expression of non-canonical modules as well as the fetal signature in therapy-resistant residual cancer cells (**Fig. 5g,h** and **Extended Data Fig. 7g**). Thus, metastasis and chemotherapy—both forms of epithelial injury that disrupt contacts between neighboring epithelial cells—converge on the selection of highly plastic cell states with multilineage tumor regenerative potential.

### *PROX1* represses non-canonical differentiation and enforces intestinal lineage identity in the fetal progenitor state

We sought to understand how cancer cell identity is restricted to the intestinal lineage in the primary tumor, but gains lineage plasticity in metastasis. Given that the fetal state is observed in primary tumors and consistently appears as a bridge to non-canonical differentiated states, which are predominantly observed in metastases (**Fig. 4b,d–f**), we hypothesized that a transcription factor (TF) may act to restrict plasticity in invasive primary tumors. An ideal candidate TF should be expressed coordinately with the fetal state across multiple patients. Through *in-silico* screening, we found six TFs that are tightly co-expressed with the fetal signature along the canonical-to-non-canonical diffusion component in KG146, KG182, KG150 and KG183 (**Fig. 6a** and Methods). Among these, only three TFs are upregulated by irinotecan chemotherapy, which induces injury repair and fetal states (**Figure 5g,h** and **Extended Data Fig. 7g**), with *PROX1* being most induced (**Extended Data Fig. 8a**).

**Figure 6.**
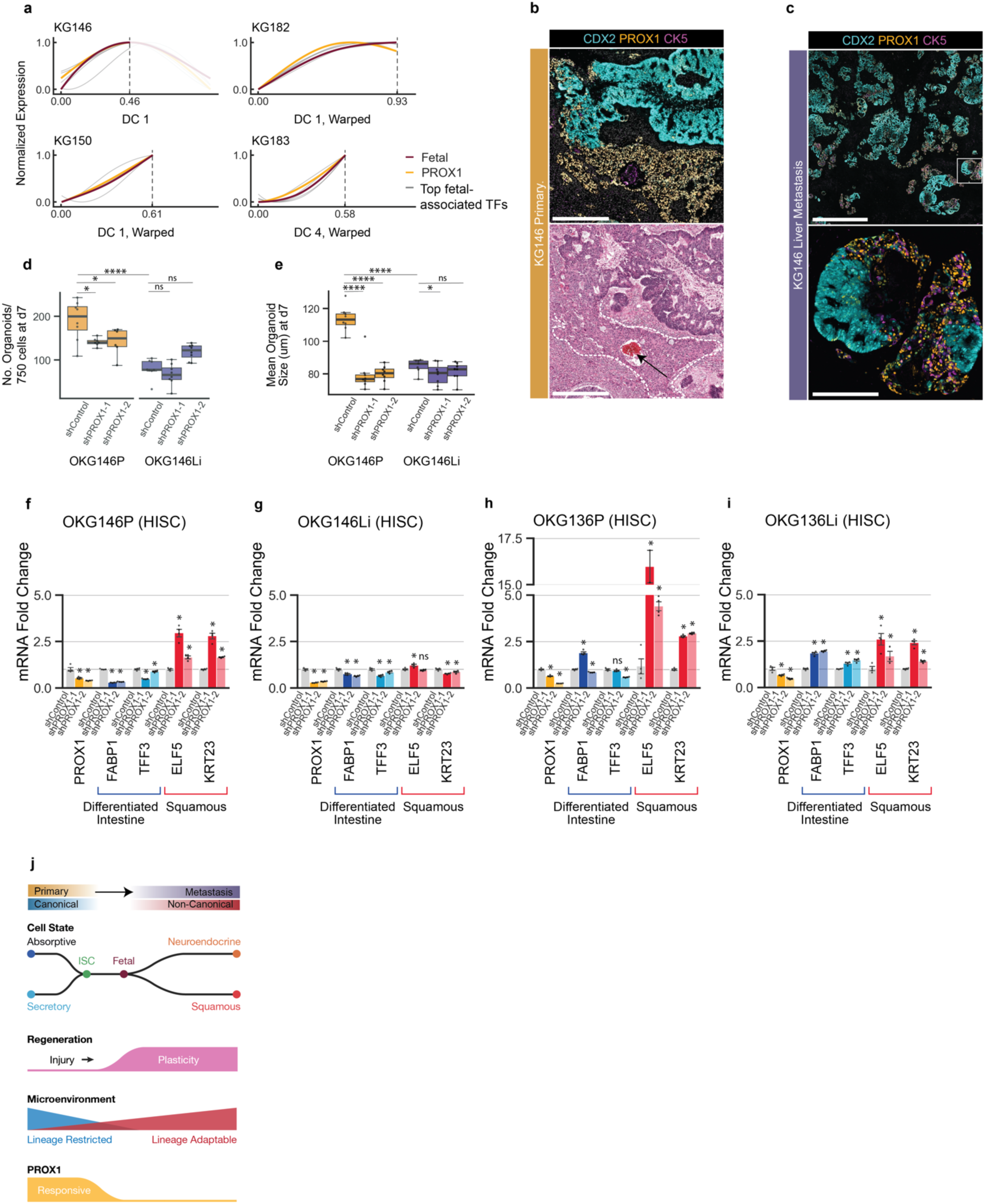
*PROX1* encodes a fetal-state-associated transcription factor that inhibits non-canonical transdifferentiation. **a,** Gene trends along aligned canonical-non-canonical axes, as in Fig. 4, of the 6 top-ranked fetal-associated human TFs in four patients. TFs are ranked according to their mean correlations of imputed gene expression with the fetal signature score in these patients. **b**, Lunaphore COMET immunofluorescence imaging of CDX2 (intestinal lineage), PROX1 (fetal state) and CK5 (squamous lineage) (top) and hematoxylin and eosin imaging (bottom) of the KG146P invasion front, showing the relationship between PROX1 expression and the transition from well-differentiated glandular morphology to poorly differentiated state. Arrow indicates a blood capillary completely encased by mucosa invasive cancer cells (scale bar = 500 μm). **c**, COMET immunofluorescence imaging of two KG146Li liver metastasis fields of view, showing spatially ordered transitions in expression of CDX2 (intestinal lineage), PROX1 (fetal state) and CK5 (squamous lineage) among metastatic tumor cells (scale bar = 1000 μm (top), 100 μm (bottom)). **d,e,** Organoid initiation capacity (number of organoids formed per 2000 single cells/40 μL matrigel) **(d)** and organoid outgrowth capacity (average organoid diameter at day 7) **(e)** of OKG146P and OKG146Li organoids expressing doxycycline- inducible shRNAs targeting *Prox1* or control shRNA at 7 days of culture in HISC media supplemented with 2 μg/ml doxycycline for 7 days. (scale bar = 1000 μm; n = 8; one-sided t-test, Benjamini-Hochberg correction, *, p < 0.05, ****, p < 0.0001). **f-i,** Relative mRNA expression of differentiated intestine and squamous differentiation markers in OKG146P **(f)**, OKG146Li **(g)** OK136P **(h)** and OK136Li **(i)** organoids expressing doxycycline-inducible shRNAs targeting *PROX1*, cultured in HISC media containing 2 μg/ml doxycycline for 7 days, in comparison with shControl organoids. RT-qPCR normalized to *GAPDH* mRNA expression (n = 4, t-test, Benjamini-Hochberg correction, * indicates p < 0.05). **j**, Model of cell-state transitions during metastatic tumor progression in CRC. Cancer cells in the primary tumor first enter into an ISC-like state. Cells at the primary tumor invasion front undergo developmental reversion into a highly plastic fetal progenitor-like state, enabling non-canonical differentiation into divergent states, including those resembling neuroendocrine and squamous cells, during metastatic outgrowth. Epithelial injury during tumor dissemination, metastasis and after therapy induces entry into the highly plastic fetal progenitor state, enabling tumor regenerative cells to express gene programs of non-canonical lineages, and dynamically adapt to diverse stresses during metastasis and therapy. Induction of the transcriptional repressor PROX1 serves to restrict non-canonical gene expression in injured normal epithelia, enabling tissue regeneration. PROX1-responsive intestinal lineage restriction is progressively lost during cancer progression, licensing non-canonical differentiation.

*PROX1* encodes a homeobox TF with roles in many lineages, including lymphatic, hematopoietic stem, neuronal, enteroendocrine and regenerative cells in the normal intestinal epithelium^58,59^. Its expression is induced by injury and is required for redifferentiation into canonical intestinal states during repair^59^. Moreover, it can act as a repressor or co-repressor, and increasing repressive activity in primary CRC tumors has been associated with more aggressive disease^60–62^. Using multiplexed immunofluorescence to define spatial context (Methods), we found that well-differentiated glandular regions of the primary tumor display high CDX2 and low PROX1 expression, whereas poorly differentiated CDX2-low cells invading blood vessels at the tumor margin express high PROX1, but retain expression of the intestinal differentiation marker CK20 (**Fig. 6b** and **Extended Data Fig. 8b**). This suggests that tissue-disruptive injury at the tumor invasion front induces PROX1, prompting us to investigate the relationship between PROX1 expression and the canonical-to-non- canonical differentiation trajectory observed in metastasis. In matched metastases, PROX1-expressing cells not only form spatially segregated subpopulations marked by low CDX2 expression; they also express the squamous-like marker CK5, with variable expression of the intestinal differentiation marker CK20 (**Fig. 6c** and **Extended data Fig. 8b**). We observe some regions with a spatially ordered transition from CDX2^high^ to CDX2^low^PROX1^high^ to PROX1^low^CK5^high^ cells, matching the observed trajectory in the RNA data (**Fig. 6c** and **Extended data Fig 8c**). Our observation of *PROX1* induction in cancer cells experiencing injury at the primary tumor invasion front suggests that it might act as a repressor of non-canonical differentiation in disrupted intestinal epithelia, in turn facilitating dedifferentiation into *LGR5*+ ISCs, and the restoration of epithelial structures once permissive niche signals become available. The corresponding prediction is that cancer cells must overcome this *PROX1*-dependent tissue regenerative lineage restriction mechanism to enable non- canonical differentiation in metastasis.

To test the function of *PROX1* in enforcing intestinal lineage identity, we engineered paired primary and metastasis (OKG146P and OKG146Li) organoids with doxycycline-inducible shRNAs targeting *PROX1* or a neutral control (**Extended Data Fig. 8d**,e). We cultured shControl and shPROX1 organoid lines in HISC media to induce ISC-like gene expression (**Fig. 5b**) and assessed their ability to regenerate organoids from single cells, an established assay of ISC function^63^. *PROX1* knockdown impairs OKG146P organoid formation and outgrowth, supporting a role for *PROX1* in regeneration along an intestinal lineage, and consistent with studies in primary tumor-derived organoids^64^ (**Fig. 6d,e**). On the other hand, dissociated cells from OKG146Li organoids, which have lost intestinal lineage restriction (**Fig. 5b**), form fewer organoids than their primary tumor counterparts, but *PROX1* knockdown has no effect on OK146Li organoid formation capacity (**Fig. 6d,e** and **Extended Data Fig. 8f**). Further, *PROX1* knockdown in OKG146P induces expression of non-canonical squamous (*Elf5*, *Krt23*) genes (**Fig. 6f**), suggesting that *PROX1* represses non-canonical gene expression in tumor cells that are in a regenerative state. Consistent with a loss of *PROX1*-dependent repression of lineage identity in lineage plastic metastatic cancer cells, *PROX1* knockdown in OKG146Li does not induce expression of non-canonical genes (**Fig. 6g** and **Extended Data Fig. 8g**). To further discern the cell state- dependent effect of *Prox1*, we engineered a pair of chemotherapy-naive primary and metastasis-derived organoids (OKG136P and OKG136Li), both comprised of cells largely in the canonical intestinal and fetal progenitor states, but not in differentiated non-canonical states (**Fig. 3e** and **Extended Data Fig. 8g**). In both of these largely canonical organoids, *PROX1* knockdown induces expression of non-canonical squamous markers, albeit to a greater extent in the more canonical OK136P (**Fig. 6h,i**). Thus, *PROX1* downregulation, independent of chemotherapy-induced or metastasis-dependent injury and priming, is sufficient to induce non-canonical differentiation. Together, our data support a model of context-specific dependency on *PROX1* lineage restriction, ISC function and tumor regeneration, with metastasis and treatment selecting for progressively less-*PROX1* dependent, increasingly lineage unrestricted states (**Fig. 6j**).

## Discussion

Our unique resource of biospecimen trios from normal colon, primary and metastatic colorectal cancer allowed us to observe and characterize the progression to metastasis in individual patients. Despite vast heterogeneity across samples, Hotspot gene module analysis charted trajectories that are broadly conserved across patients. As shown in previous studies, differentiated intestinal states first dedifferentiate into an *LGR5*+ intestinal stem cell-like state in the primary tumor, and invasive cancer cells enter into an injured state marked by high L1CAM/EMP1 expression, which is associated with metastasis-initiating cells^14,15^ (**Fig. 3a**). We now demonstrate that metastatic colonization is associated with progressive cell state transitions away from canonical intestinal states into a highly conserved fetal progenitor state, followed by lineage reprogramming into squamous-like, neuroendocrine-like and other states (**Figs. 3b–d** and **6j**). Our data unequivocally demonstrate that the selection of cell states in human CRC metastases is distinct from those that dominate primary tumors. Further, we delineate context- and site-specific functions of molecules and signaling pathways during tumor progression that could not be predicted using biospecimens from primary tumors alone. Our investigation of cancer cell-intrinsic and microenvironmental factors suggest a two-stage model of metastatic plasticity, whereby the loss of epithelial intercellular contacts during tumor dissemination or therapy first induces a multipotent state, then tumor microenvironmental factors drive differentiation towards diverse intestinal and non-intestinal lineages.

While plasticity is emerging as a hallmark of advanced cancer, our study provides a roadmap for studying the mechanisms and consequences of cell state reprogramming during cancer progression. We previously demonstrated that epithelial injury during tumor dissemination and therapy, acting via the loss of E-cadherin-mediated intercellular contacts, induces entry into an L1CAM+ tumor regenerative, metastasis-initiating state^14^. Here, we further demonstrate the selection of an L1CAM-expressing injury repair state that precedes non-canonical differentiation during metastatic outgrowth. Further, the increased cell-autonomous plasticity of metastatic tumors that we identify suggests a mechanistic explanation for the close link between metastasis and therapy resistance. The ability to dynamically express diverse gene programs in response to stresses and microenvironmental cues, unique to metastases in principle confers the ability to rapidly evade therapy. Whether specific pre-existing genomic alterations in primary tumors can endow cancer cells with greater epigenetic flexibility and likelihood of entering non-canonical states remains to be seen. We demonstrate that transcriptomic signatures of non-canonical states can serve as biomarkers of poor outcomes in two independent cohorts of pre-treatment primary CRC tumors. Thus, cells with the ability to enter into non- canonical states emerge in primary tumors and may be selected for during metastasis, although dynamic phenotypic adaptation into non-canonical states is likely further induced during tumor progression and by therapy. Our finding that non-canonical gene signatures can serve as prognostic biomarkers of future disease relapse and poor survival, could be clinically useful in identifying patients most likely to benefit from aggressive neoadjuvant or adjuvant therapy targeting non-canonical states to prevent future macrometastasis.

Unlike highly plastic metastases, primary tumor cells remain epigenetically fixed in an intestinal lineage, in part mediated through the repressor activity of *PROX1*. Our data are consistent with a model of context- dependent *PROX1* function along a continuum of canonical differentiation to dedifferentiation and non- canonical differentiation during tumor progression (**Fig. 6j**). In early tumorigenesis, as shown in mouse models of intestinal tumor initiation^64–66^, *PROX1* functions as a tumor promoter, with increasing levels of *PROX1* reinforcing ISC-like states by inhibiting canonical differentiation into enteroendocrine and other differentiated cell states^67^. As tumor cells progress into injury repair and fetal-like states at the primary tumor invasion front, *PROX1* stabilizes the phenotype and enables redifferentiation along a canonical trajectory in the presence of intestinal niche factors, accompanied by *PROX1* downregulation. Metastatic colonization, in contrast, selects for cells that are further along the canonical-non-canonical axis, that have downregulated *PROX1* and are insensitive to further inhibition of *PROX1* expression. In other words, the phenotypic consequences of *PROX1* levels in a given cancer cell depend on how far the cell has progressed along the canonical-non-canonical differentiation axis. Future work will likely uncover additional mechanisms by which plasticity is restricted in primary tumors and expands in metastasis.

The non-canonical states we identified in patient metastases were not captured in mouse models of CRC metastasis^15,16^. This could reflect the differences in timing of metastasis, which occurs over weeks to months in mice, but takes years in patients. At the time of resection, patient metastases are larger than 1 cm^3^ (frequently much larger) and therefore contain more cells that have undergone more cell divisions than their murine counterparts. Larger metastases may also experience greater intratumoral hypoxia or other sources of stress that could promote the emergence of non-canonical states. Further, our data suggest that injury to epithelial structures can induce non-canonical differentiation (**Fig. 6b**). In patients, metastases likely undergo variable rounds of attempted proliferation and immune editing, resulting in a dynamic equilibrium with multiple injury-repair cycles and clinical dormancy before the eventual outgrowth of macrometastases^1^. This period of injury-repair may be important in enabling or selecting for non-canonical differentiation, and may be poorly captured in mouse models selected for rapid metastatic outgrowth. Most clinical cancer genomics studies, including the TCGA, have focused their analysis on primary tumors, with the underlying assumption that metastases are likely to have similar phenotypes. Yet, new cancer therapeutics are almost always first tested in patients with advanced metastatic disease in clinical trials^1,68^. The fact that we identify cell states in metastases that we do not find in primary tumors from the same patients underscores the limitations of extrapolating from primary tumors, and highlights the need to study metastatic tissue and patient metastasis-derived *ex vivo* models to delineate therapy response and plasticity mechanisms.

Phenotypic plasticity poses a major challenge to cancer therapy^54^, but the identification of cell states and trajectories that are conserved across multiple patients suggest future opportunities to target plasticity. One approach is to target conserved regenerative intermediates such as injury repair or fetal-like states. The clinical challenge in this case would be to identify time points where sufficient cells are in these specific states. Identifying conserved trajectories, on the other hand, may pave the way for combination or sequential therapy with drugs that anticipate and preemptively target resistance states. For example, LGR5^+^ ISC- targeting drugs could be followed with drugs targeting squamous-like or neuroendocrine-like states, or perhaps more impactfully, it could be possible to target the molecular machinery that enables dynamic entry into resistant states. While identifying mechanisms that are unique to cancer cell reprogramming and dispensable for normal tissue homeostasis remains an important challenge, the current study provides a roadmap for understanding and eventually targeting progressive plasticity in advanced cancer.

## Acknowledgements

This work was supported by NIH grants U2CCA233284 (D.P., K.G.), U54CA209975 (D.P., K.G.), R37CA266185 (K.G., J.S.), K08CA230213 (K.G)., T32GM007739 (E.K.B., K.L.), U01CA23844401A1 (W.J.), R01EB027498A1 (W.J.), P30CA008748 (MSKCC), Howard Hughes Medical Institute Investigator (D.P.) and Gilliam Fellowship (A.M.), NSF GRFP (A.M.), Damon Runyon Clinical Investigator Award (K.G.), Burroughs Wellcome Career Award for Medical Scientists (K.G.), AACR NextGen Grant for Transformative Cancer Research (K.G.), Stand Up to Cancer Convergence 3.1416 Award (K.G.), Pershing Square Sohn Prize (K.G.), Starr Cancer Consortium (K.G.), Josie Robertson Investigator Award (K.G.) and Gerry Metastasis and Tumor Ecosystems Center grants (Q.J., D.P., K.G.).

## Declaration of Interests

D.P. is on the scientific advisory board of Insitro. K.G. is a consultant for Seres Therapeutics and is an inventor on patents related to targeting metastasis. J.S. is a consultant for Paige AI.

---

**Extended Data Figure 1.**
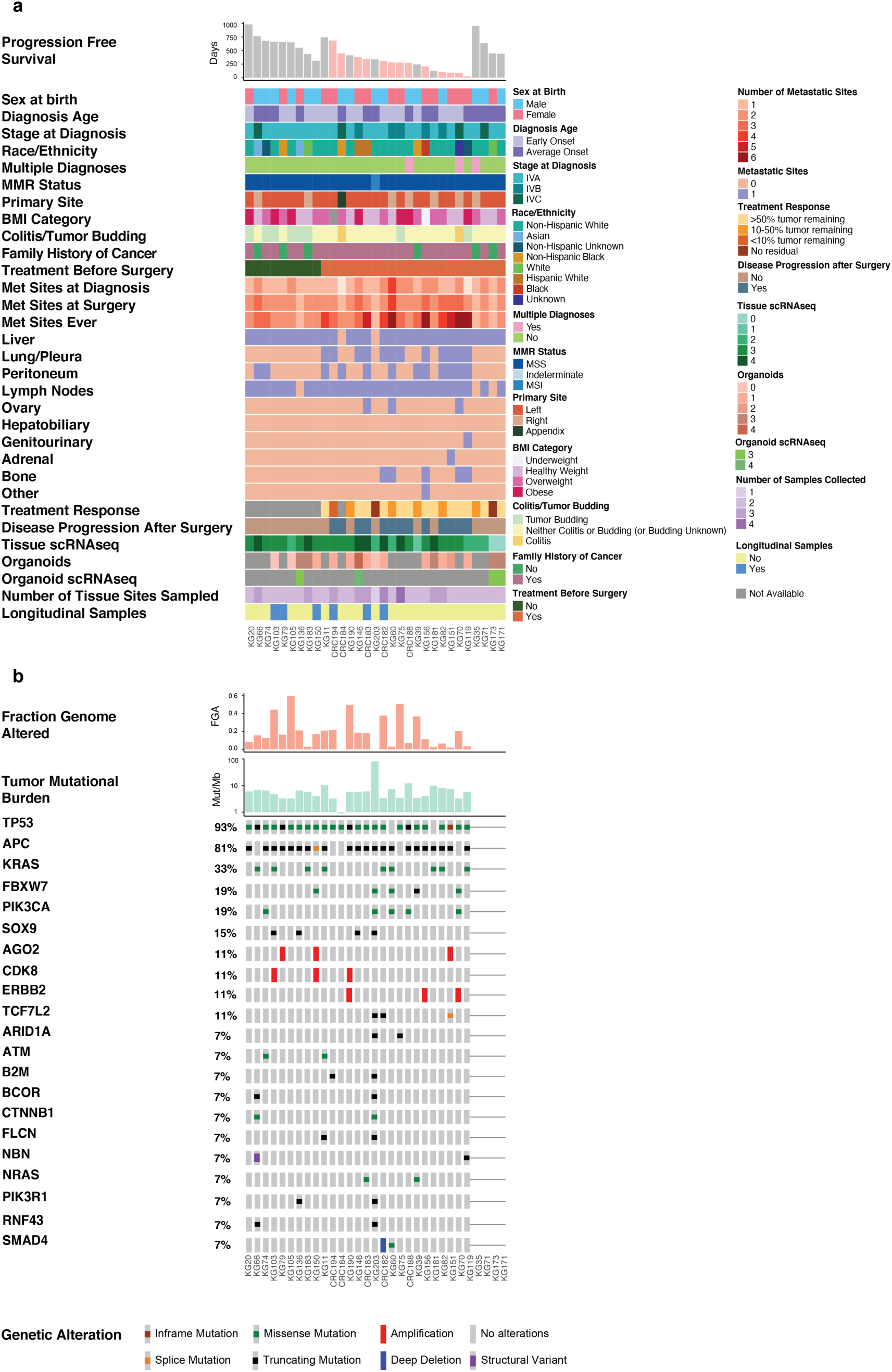
Clinical and genomic characteristics of metastatic colorectal cancer biospecimen cohort. **a,** Clinical characteristics of patients in the study cohort, including key demographic, clinical, metastatic site, treatment and outcome variables, as well as whether scRNA-seq data was collected from each biospecimen or organoid. Longitudinal samples refer to additional metachronous tumor samples collected at time of progression, subsequent to initial synchronous tissue collection. No more than one biospecimen was sequenced per site. **b**, Genomic features of tumors sequenced using the MSK-IMPACT platform^69^. Tumor- associated mutations were filtered based on OncoKB annotations. Percentage of tumors with a mutation in a given gene is indicated at left. The most frequently mutated genes were *TP53* (93%), *APC* (81%) and *KRAS* (33%). The most frequently amplified genes were *AGO2* (11%), *CDK8* (11%), and *ERBB2* (11%). Fraction genome altered and tumor mutational burden are shown at top. Mut/Mb = average number of mutations per megabase.

**Extended Data Figure 2.**
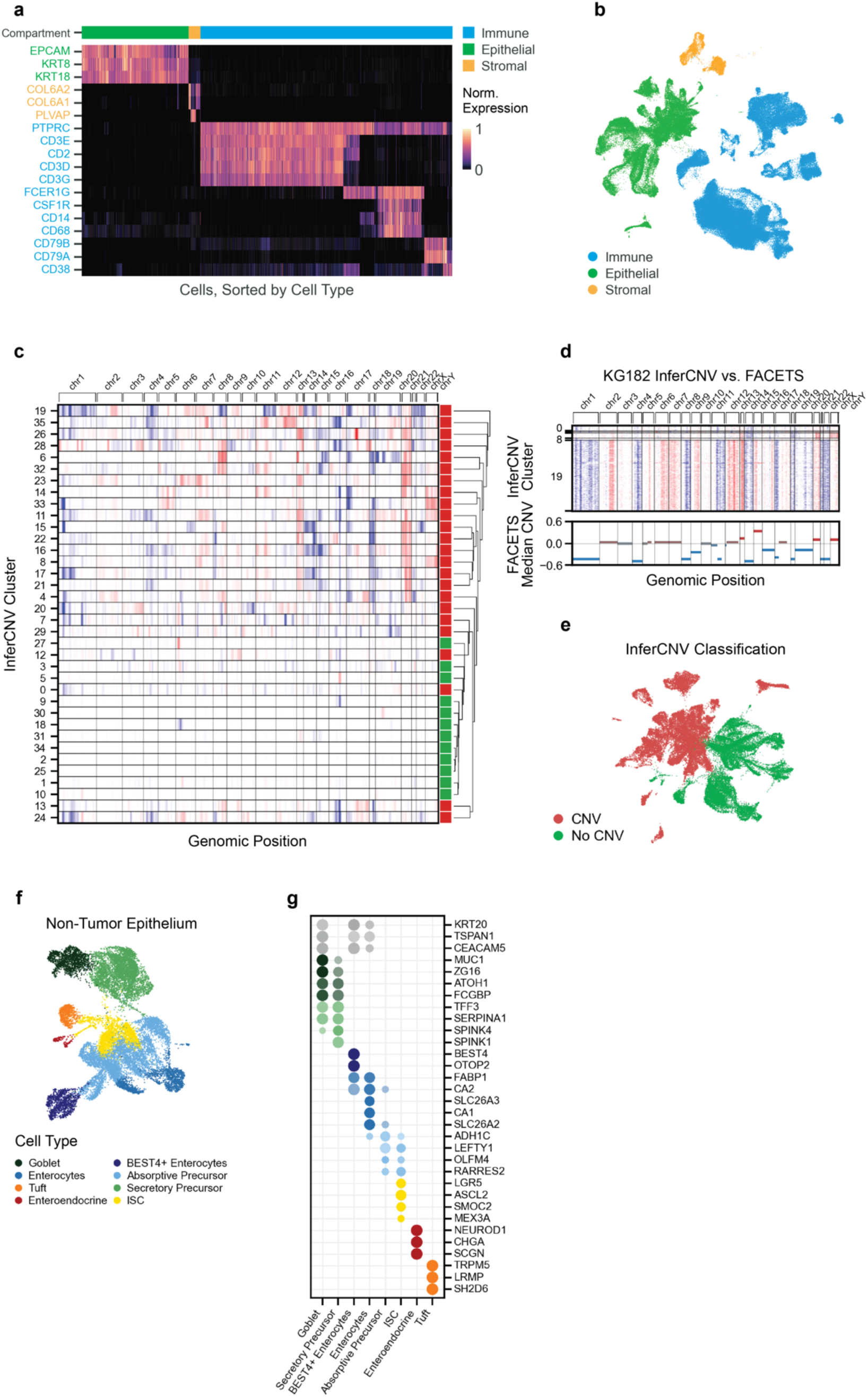
Clinical and genomic features of patient cancers and collected biospecimens. **a**, Expression of canonical marker genes across all cells grouped by compartment, cell type and cluster. Each column is a cell and each row is a gene; color values represent normalized transcript counts (Methods). **b**, UMAP embedding of all cells colored by immune (111,609 cells), epithelial (47,437 cells) or stromal (5,258 cells) compartment. **c**, Copy number changes calculated with InferCNV^23^ and binned by genomic region (Methods), per epithelial cell cluster. Rows represent the mean inferred copy number relative to a diploid reference population of non-tumor epithelium, for each cluster of cells. Clusters are calculated according to cells’ inferred copy number profiles. Cancer cells are called on a per-cluster basis according to their mean copy number profile, shown in the right column as green (no CNV) or red (CNV). **d**, Inferred copy number of tumor cells for a representative patient (top, mapping to three inferCNV clusters) compared to copy number values estimated by FACETS^70^ using bulk sequencing for a targeted gene panel (bottom; see Methods). **e**, UMAP of all epithelial cells as in Fig. 1d, colored by CNV classification. **f**, UMAP of all non-tumor epithelial cells (21,297 cells), colored by cell type annotation (Methods). **g**, Expression of colon epithelial cell type markers across annotated cell types. Dot size scales with the proportion of cells in a cell type that express each gene; color intensity indicates the mean *z*-scored, log-normalized expression of each gene. Rows (genes) are colored by the cell type (as in **f**) for which they are a marker.

**Extended Data Figure 3.**
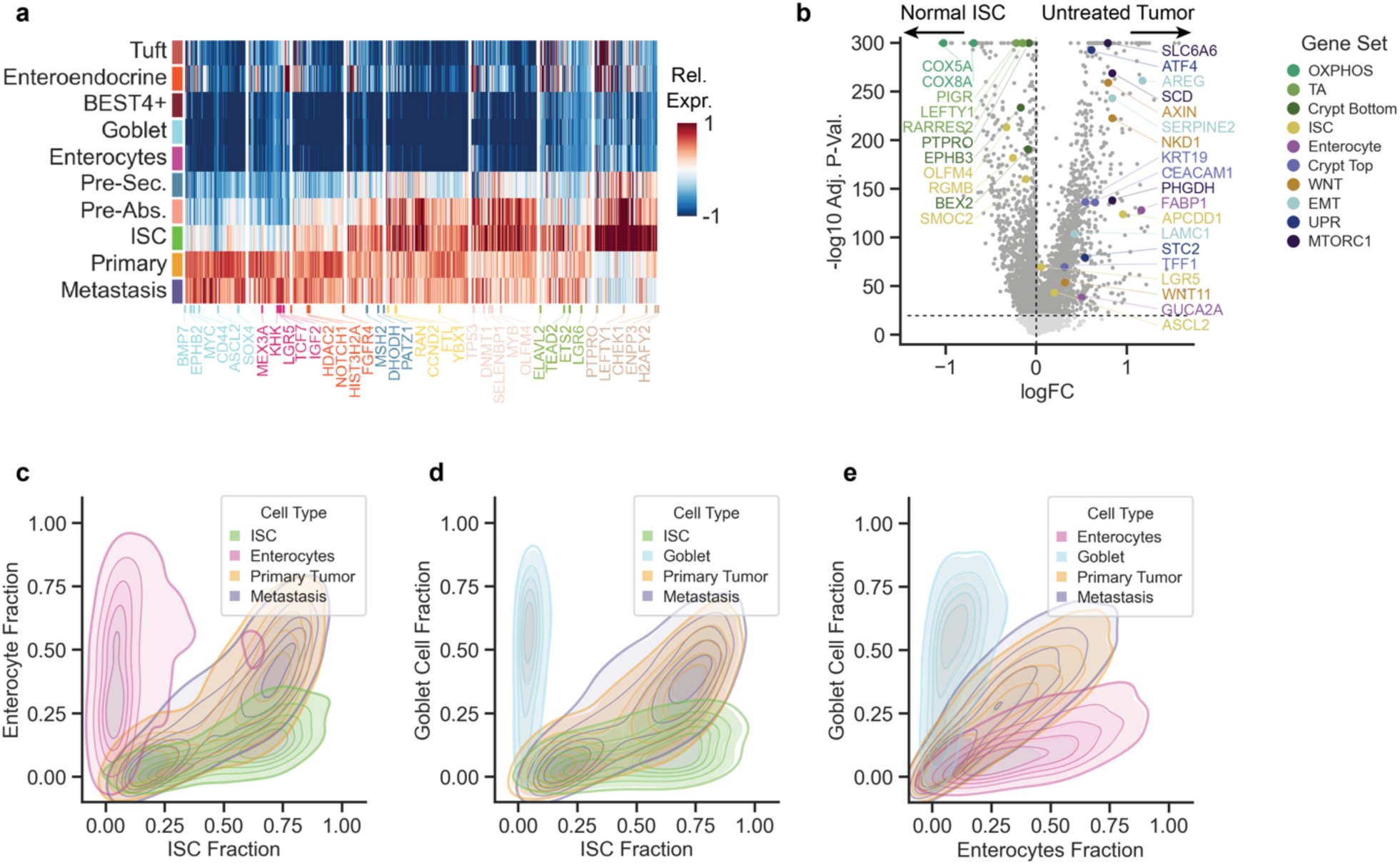
Untreated CRC tumors express early stem cell and mixed lineage programs. **a**, Mean expression of ISC gene signature genes, *z*-normalized per gene across cells, within all treatment- naive epithelial cell types (**Supplementary Table 1k** and Methods). **b**, Genes differentially expressed between ISCs and all treatment-naive tumor cells. Highlighted gene programs are among the top-enriched gene sets according to GSEA performed on genes ranked by differential expression in tumor cells (Methods), and highlighted genes are from leading edge subsets. OXPHOS, oxidative phosphorylation; TA, transit amplifying; ISC, intestinal stem cell; WNT, WNT-beta-catenin signaling; EMT, epithelial-mesenchymal transition; UPR, unfolded protein response; MTORC1, MTORC1 signaling. logFC, log fold-change. -log10 Adj. P-Val., -log10 adjusted P value. **c**, Two-dimensional density plot displaying the abundance of ISC- and enterocyte-specific DEGs in normal ISCs and enterocytes, and primary tumor and metastasis cells from untreated patients (Methods). Only markers that show high expression in tumors (in the top quartile per gene across all tumor cells) are plotted. **d**, Same as **c**, for ISC and goblet cell markers. **e**, Same as **c**, for enterocyte and goblet cell markers.

**Extended Data Figure 4.**
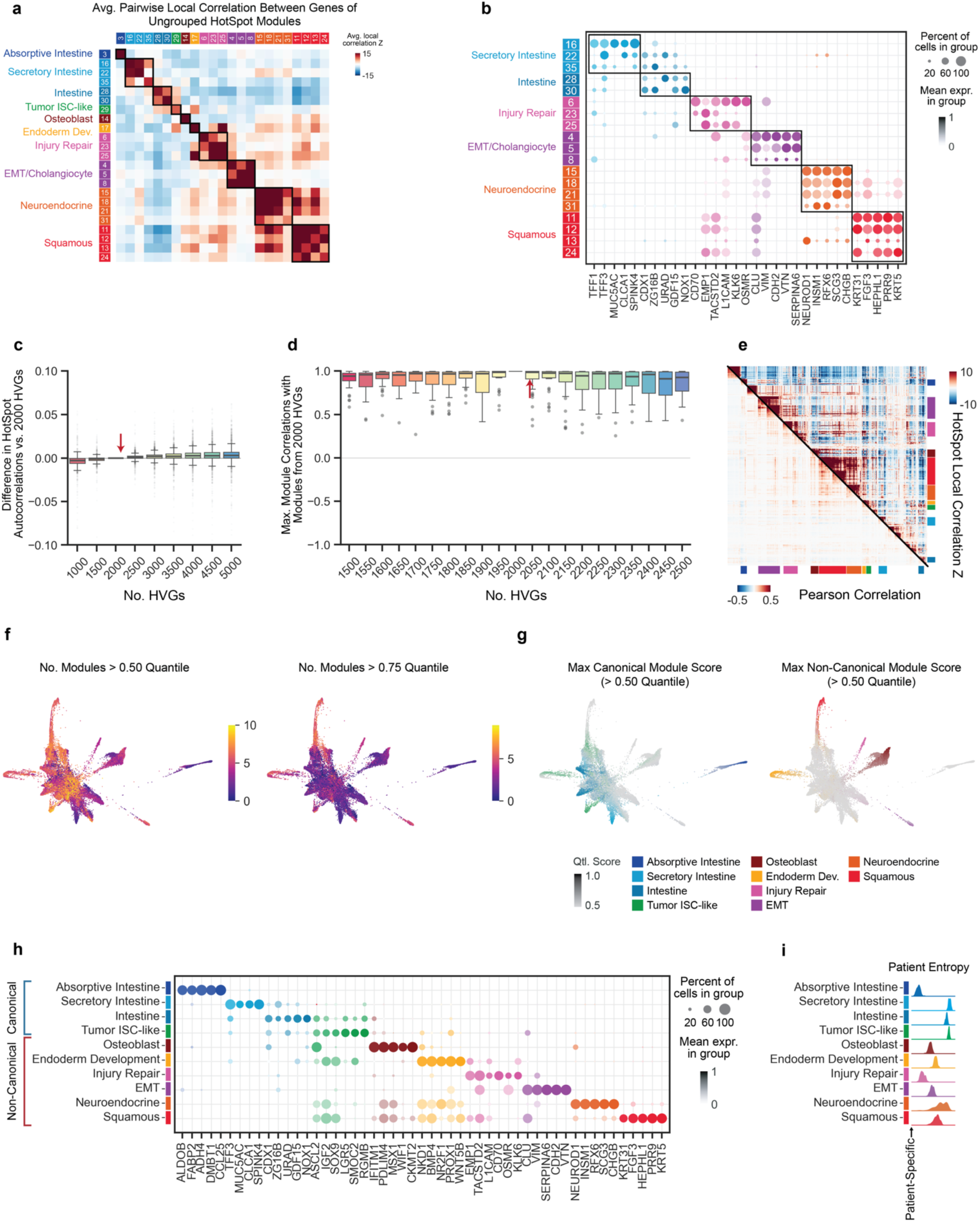
Hotspot identifies canonical intestinal and non-canonical gene expression modules in tumor cells from the full metastatic CRC cohort. **a,** Heatmap showing average pairwise local correlation score (Methods) between HotSpot modules before grouping. x-axis and y-axis show each module as annotated. Each entry is the average local correlation between all pairs of genes assigned to the corresponding modules of the x-axis and y-axis. Squares above and to the left are colored according to the final annotation of the modules after grouping. **b**, Z-normalized gene expression within groups of cells (y-axis) which include the top-scoring cells for each ungrouped HotSpot module (Methods). Columns of dots are colored according to the HotSpot module assignment of the gene on the x-axis. Dot size indicates proportion of cells in a cluster that score for that gene module; color intensity indicates the mean module score of that cluster. Squares to the left are colored according to the final annotation of the modules after grouping. **c**, Robustness of Hotspot local autocorrelation as a function of the number of highly variable genes (HVGs) chosen (Methods). y-axis depicts the difference in gene autocorrelation score using varying numbers of HVGs (x-axis) and using 2,000 HVGs (used throughout this manuscript). Plots display median (line), interquartile range (boxes) and 1.5x interquartile range (whiskers). **d**, Robustness of Hotspot gene modules as a function of the number of HVGs used. y-axis depicts Pearson correlation between Hotspot gene modules in this study (based on 2,000 HVGs) and the most similar gene modules produced using an alternate number of HVGs (x-axis) (Methods). **e**, Hotspot local correlation is more sensitive than global correlation. Each row and column is a gene and each entry quantifies gene-gene local correlations in CRC tumors computed with Hotspot (upper triangle) or Pearson correlations (lower triangle), for all 1,962 genes among the top 2,000 HVGs in the complete CRC tumor dataset, with a false discovery rate (FDR) < 0.01, as determined by Hotspot. **f**, FDL of all tumor cells (26,145 cells) as in Fig. 3b. Each cell is colored by the number of HotSpot modules for which it scores higher than the 0.5 (left) or 0.75 ^59^ quantile module score across all cells. **g**, FDL of all tumor cells as in **f** colored by gene module score for 4 canonical (left) or 6 non-canonical ^59^ CRC gene modules. Each cell is colored according to its maximum module score, or gray if no score exceeds the module’s 50th percentile across all cells. **h**, Z-normalized gene expression within groups of top-scoring cells for each grouped HotSpot module as in **b**. **i**, Kernel density plots depicting the entropy across patients within high-scoring tumor cells for each module (Methods). High entropy indicates that cells which score highly for a module come from a well-mixed set of patient samples; low entropy indicates that cells primarily originate from a single patient. A cell is considered high scoring for a module if its module score is greater than one s.d. above the mean, calculated across all tumor cells.

**Extended Data Figure 5.**
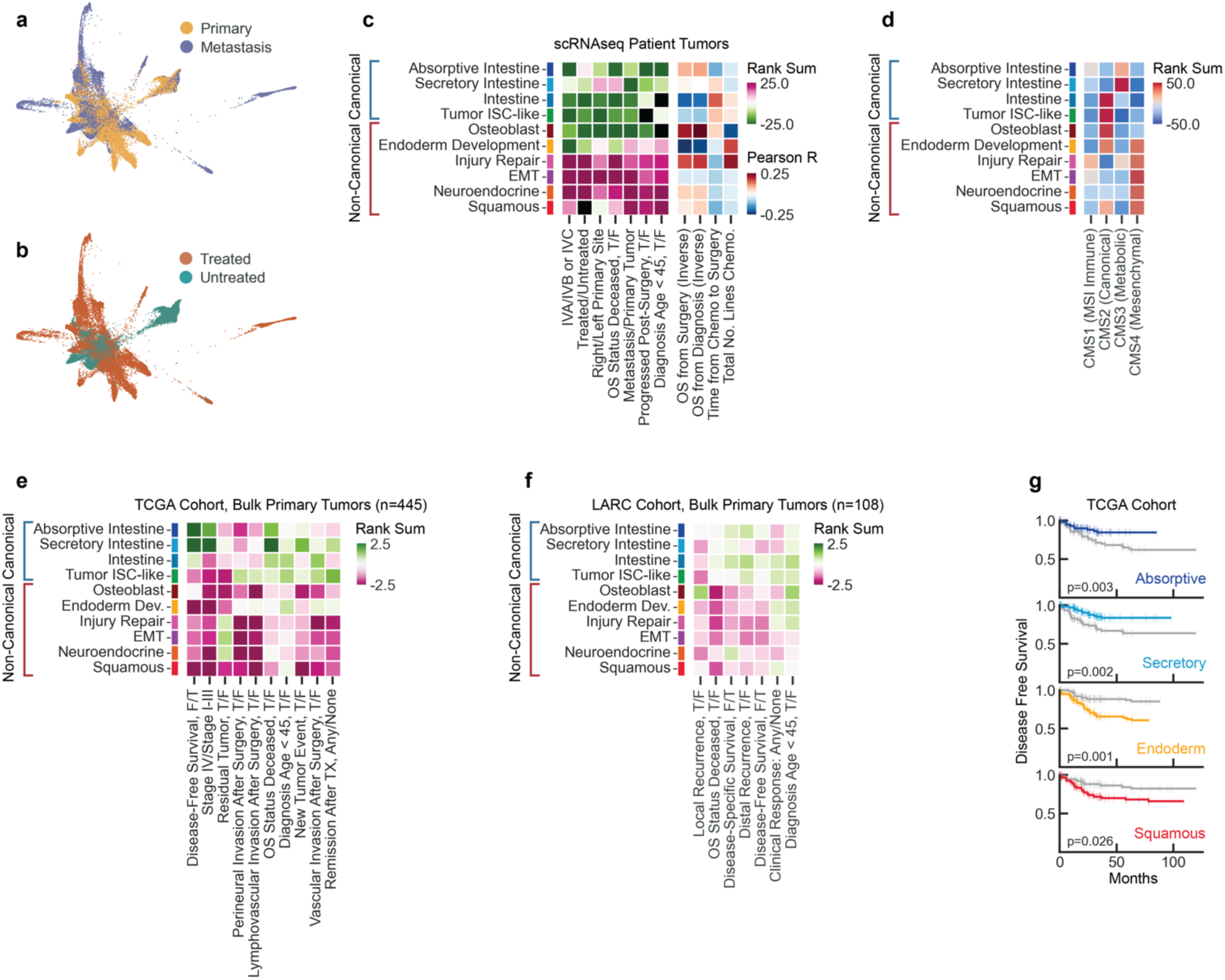
Clinical features and outcomes associated with canonical and non-canonical module expression. **a**, FDL of all tumor cells (26145 cells) as in Fig. 3b, colored by primary or metastasis. **b**, FDL of all tumor cells, colored by treatment status. **c**, Associations of patient baseline clinical, pathological, treatment and survival attributes with gene module scores in scRNA-seq data. Mann-Whitney U test is used for all binary attributes, Pearson correlation for all continuous attributes. **d**, Associations of CRC consensus molecular subtype (CMS) classifications^27^ with gene module scores in scRNA-seq data (Methods). **e**, Associations of patient baseline clinical, pathological, treatment and survival attributes with module ssGSEA enrichment scores for 445 patients with colon (COAD) or rectal (READ) adenocarcinoma from the TCGA cohort^45^. **f**, Same as **e**, for 108 patient samples with locally advanced rectal cancer (LARC^46^). **e**,**f**, Mann-Whitney U test is used for all binary attributes. **g**, Kaplan-Meier plots showing disease-free survival for patients in the TCGA cohort with high or low ssGSEA enrichment for the indicated signatures. A patient is signature-high if the enrichment score of their tumor is > 1 s.d. above the mean, calculated across all patients, and signature-low if it is < 1 s.d. below the mean (n = 147, 158, 144, and 123 for Absorptive, Secretory, Endoderm, and Squamous, respectively)

**Extended Data Figure 6.**
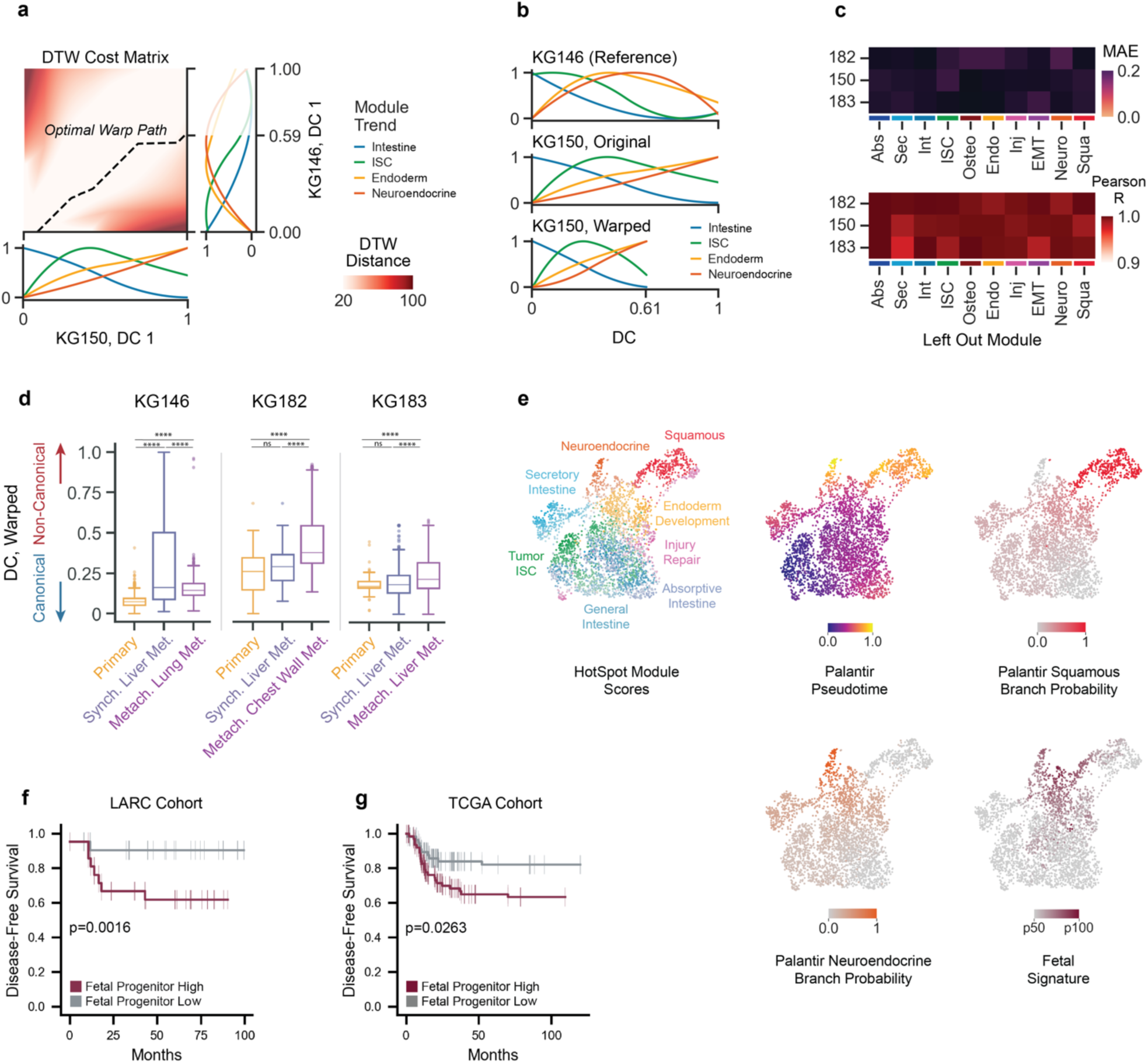
Dynamic time warping identifies conserved progression of cell states across patients. **a,** Accumulated cost matrix for all dynamic time warping (DTW) paths between DCs for patients KG146 and KG150. The matrix is calculated for trends in patients’ module scores (at right and below). Dashed black line depicts the optimal warping path between DCs. **b**, Top, gene module score trends calculated over the first DC in patient KG146, which is used as a reference for DTW alignment of other patient DCs; for example, DC1 for KG150, shown before (middle) and after (bottom) alignment. **c**, Top, mean absolute error (MAE) between the DTW path for each patient DC and a DTW path computed after leaving out one module score. Values were only calculated on timepoints shared by both sequences (Methods). Bottom, Pearson correlations computed for the same paths as above. **d**, Distributions of cells from primary tumor, synchronous (Synch.) liver metastasis, and longitudinally collected metachronous (Metach.) metastasis from lung (KG146), chest wall (KG182), or liver (KG183) along the DTW-aligned DC representing the canonical-non-canonical axis (t-test, Bonferroni correction, ****, p < 0.0001). Timing of longitudinal metachronous metastasis collection, and treatments administered between initial and secondary metastasis collection are indicated in Supplementary Data Figure 1. **e**, UMAP of 3351 cells from KG146 primary tumor, synchronous liver metastasis, and metachronous lung metastasis. Cell are colored by their maximum module score or in gray if no module score exceeds its 25th percentile (top-left), Palantir pseudotime (top-middle), Palantir branch probability for the squamous-annotated (top-right) and neuroendocrine-annotated (bottom-left) branches, and the fetal signature (bottom-left). **f,** Kaplan-Meier plots showing disease-free survival for patients in the LARC cohort with high or low ssGSEA enrichment for the fetal progenitor signature. A patient is fetal-progenitor-high if their enrichment score is > 1 s.d. above the mean, calculated across all patients, and fetal-progenitor-low if it is < 1 s.d. below the mean (log-rank, n = 42, *p* = 0.0350). **g**, Same as **f**, for patients in the TCGA cohort (log-rank, n = 134, *p* = 0.0016).

**Extended Data Figure 7.**
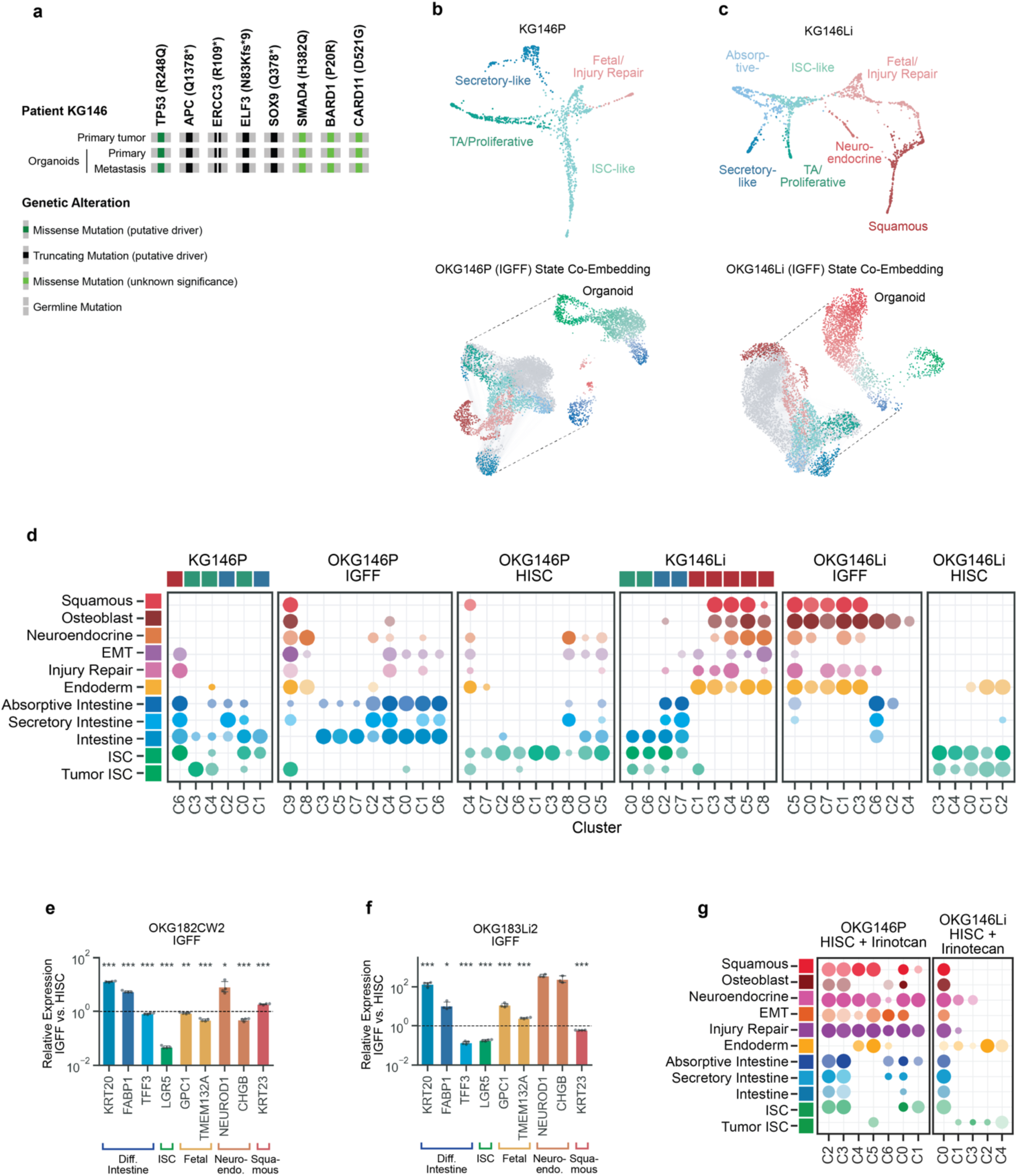
Transcriptomic comparison of patient tumors and organoids deconvolves cell-autonomous and microenvironmental contributions to cell state. **a,** MSK-IMPACT results showing concordant mutation profiles of primary tumor and patient-derived organoids for patient KG146. **b**, Top, FDL of KG146 primary CRC tumor cells (880 cells) colored by cluster-level cell state annotations (Methods). Bottom, UMAP of OKG146P organoid cells cultured for 7 days in IGFF (upper right) and UMAP of these cells co-embedded with KG146P primary tumor cells (lower left). An augmented k-nearest neighbor graph is used to calculate organoid cell state probabilities; lines denote graph edges and cells are colored by maximum probability (Methods). **c**, Same as **b**, for KG146Li patient tumor cells and OKG146Li patient-derived organoid cells (1,279 cells) cultured for 7 days in IGFF. **d**, CRC gene module scores within cell clusters for patient tumor and organoid scRNA-seq samples (Methods). Dot size indicates proportion of cells in a cluster that score for that gene module; color intensity indicates the mean module score of that cluster. Cluster assignments are made within each sample, and scores are calculated separately for all patient tumor samples and all organoid samples. Squares above patient tumor samples indicate the cluster classification (blue, differentiated intestine-like states; green, ISC-like and TA-like states; red, non-canonical states). **e**,**f**, Relative expression of differentiated intestine, ISC, fetal and non-canonical differentiated states in patient-derived organoid line OKG182CW2 (secondary metastasis from chest wall) (**e**) or OKG183Li2 (secondary metastasis from liver) (**f**) cultured for 7 days in IGFF compared to HISC media. RT-qPCR normalized to *GAPDH* mRNA expression level (n = 4, Mann Whitney rank-sum test, *, p < 0.05, **, p < 0.01, ***, p < 0.001****, p < 0.0001). **g**, Gene module scores within cell clusters, as in **d**, for patient tumor and organoids cultured in HISC media and treated with irinotecan.

**Extended Data Figure 8.**
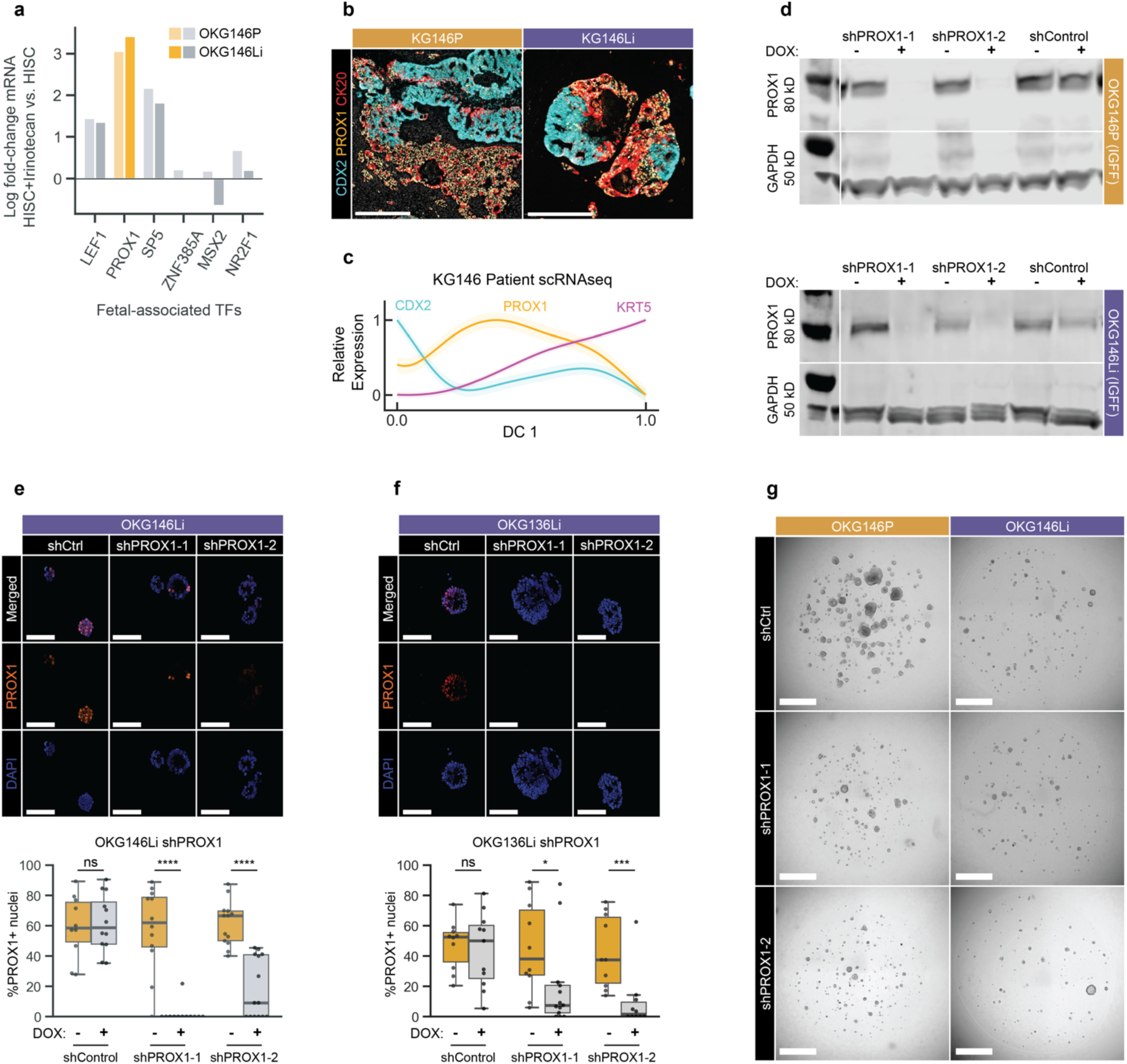
*PROX1* is a context-dependent stabilizer of intestinal identity in an epithelial injury-associated, fetal-like state. **a**, Relative expression, from scRNA-seq data, of the top 6 fetal-associated TFs in OKG146P (light bars) and OKG146Li (dark bars) organoids cultured for 7 days in HISC media containing 250 nM irinotecan, compared to HISC media alone. **b**, Representative COMET immunofluorescence images of CDX2 (intestinal lineage), PROX1 (fetal state) and CK20 (intestinal differentiation) staining of KG146P invasion front KG146Li liver metastasis fields of view (scale bar = 1000 μm (left), 100 μm (right)). **c**, Trends in gene expression of CDX2, PROX1, and KRT5 along DC1 observed in all tumor cells from patient KG146. **d,** Top: Western blot showing expression of PROX1 or GAPDH loading control in OK146P (top) or OKG146Li (bottom) organoids expressing doxycycline-inducible shRNAs targeting *PROX1* or control, cultured in IGFF medium with 2 μg/ml doxycycline for 7 days. **e,** Representative images (top) and quantification (bottom) of whole mount immunofluorescence of OKG146Li organoids expressing shRNAs targeting *PROX1* or control, cultured in HISC media with 2 μg/ml doxycycline for 7 days, stained for PROX1 (red) and DAPI (blue, scale bar = 100 μm). **f,** Representative images (top) and quantification (bottom) of whole mount immunofluorescence of OKG136Li organoids, as in **e**, (scale bar = 100 μm) **g,** Representative images of OKG146P and OKG146Li organoids, 7 days after plating 750 single cells/15 μL matrigel in HISC+2 μg/ml doxycycline (scale bar = 1000 μm).

## Methods

### Patient biospecimen procurement and processing

#### Tissue collection

Patients undergoing synchronous colorectal resection and metastasectomy at MSKCC were identified by chart review, and those who had signed pre-procedure informed consent to MSK IRB protocols #06-107, #12- 245, #14-244 and #22-404 for biospecimen collection were selected for this study. Freshly resected surgical tissue in surplus of clinical diagnostic requirements was processed into single-cell suspensions for single cell RNA sequencing, and where sufficient tissue was available, processed to generate organoids. Portions were also fixed in formalin and embedded in paraffin. Tissue was generally processed within 1 hour of surgical resection. Archival formalin-fixed, paraffin-embedded (FFPE) clinical tissue blocks for immunostaining were identified by database search and chart review. Tissue processing and histopathological data interpretation were overseen by an expert gastrointestinal pathologist (J.S.). Where trios of normal colon, primary CRC and metastatic CRC were successfully collected, the patient was longitudinally tracked through their clinical course at MSK using MSK Darwin^71^, and tumor tissue surplus to diagnostic requirements was collected from any subsequent procedures.

#### Patient metadata

Clinical data, including baseline demographic data and prior treatments, were abstracted via manual review of patient electronic medical records by board certified medical oncologists (M.L.; K.G.), collected as part of institutional review board approved protocols (MSK IRB #14-244 and #22-404). Time to each treatment event was calculated from the date of diagnosis to allow for comparison across patients. Study data were collected and managed using REDCap electronic data capture tools hosted at MSKCC on secure central servers. 17/31 patients had multiple metastatic sites at the time of surgery, and had >50% of tumor sites remaining following surgery. 17/31 patients had early onset colorectal cancer (age of diagnosis <50). Clinical MSK- IMPACT targeted exon sequencing was performed on tumor/normal tissue from 27/31 patients and revealed expected mutations (**Figure S1B**)^69^. Consistent with the low percentage (<5%) of metastatic CRC that are mismatch repair deficient/microsatellite instability high (dMMR/MSI-H)) CRC, 1/31 patients in our cohort had an MSI-indeterminate tumor. Clinical data collection was censored on September 30, 2022.

#### Tissue processing

We collected 50–300 mg of freshly resected surgical tissue in 5 ml of intestinal growth factor free (IGFF) organoid media ((Advanced DMEM/F12 (AdDF12; Thermo Fisher Scientific), GlutaMAX (2 mM, Thermo Fisher Scientific), HEPES (10 mM, Thermo Fisher Scientific), N-Acetyl-L-cysteine (1 mM, Sigma-Aldrich), B27 Supplement with Vitamin A (Thermo Fisher Scientific)) supplemented with Primocin (100 μg/ml, InvivoGen), Plasmocin (50 μg/ml, InvivoGen), Pen/Strep (100 μg/ml, Thermo Fisher Scientific)), Amphotericin B (2.5 μg/ml, Cytiva), Nystatin (250 units/ml, Millipore Sigma)). For primary and metastatic tumors, specimens were placed into a 15 cm petri dish using sterile forceps and washed 3x with DPBS (Thermo Fisher Scientific) supplemented with the above antibiotic cocktail, and minimally chopped with sharp sterile blades to enable transfer of tumor fragments using a pre-wet 25 ml serological pipette.

Tumor fragments were transferred into a gentleMACS type C tube (Miltenyi) pre-filled with 5 ml of IGFF media supplemented with antibiotics, DNase I (100 units/ml, Millipore Sigma) and a commercial cocktail of tissue digestion enzymes (Tumor Dissociation Kit, Miltenyi). Tumors were digested using the gentleMACS Octo Dissociator on the manufacturer’s 37C_h_TDK_1 protocol for a maximum of 30 minutes. Considering the heterogeneity of tissue specimens (i.e. cell viability, immune infiltration, blood content, necrosis, and calcification), the digestion state of the tumor fragments was assessed every 10 minutes under an inverted microscope. Digestion was interrupted before 30 minutes if at least 50% of the tumor material appeared broken into clusters of 1-to-10 cells. Next, cell cluster solutions were filtered through a 100 μm cell strainer, and washed 3x with DPBS supplemented with antibiotics, with each centrifugation step carried out at 100g for 3 minutes at room temperature. The final cell suspension was filtered through a 100 μm cell strainer, washed, and centrifuged for 5 min at 500 *g* and 4°C.

Non-tumor tissue was transferred into a 50 ml tube pre-filled with 25 ml dissociation/chelation buffer (8 mM EDTA, 0.5 mM DTT, Dnase I (100 units/ml, Millipore Sigma)). Mucosal fragments were incubated with gentle rotation at 4°C for a maximum of 30 minutes. The dissociation state of the tissue fragments was assessed every 10 minutes under an inverted microscope. Dissociation was interrupted before 30 minutes if at least 30% of the mucosal material appeared broken into clusters of 1-to-5 colonic crypts. Next, the crypt solution was filtered through a 1 mm cell strainer (PluriSelect) to separate individual crypts or small crypt clusters from large chunks of undissociated mucosa. Dissociation was quenched using an equal volume of DPBS supplemented with antibiotics. At this point, the 1 mm filter was flipped and inverted into a fresh 50 ml tube. Up to 25 ml of DPBS supplemented with antibiotics was flashed through the inverted filter to recover the undissociated mucosal tissue. Upon manually shaking such suspension of mucosal tissue fragments (e.g., approximately 5 times), the collection of clusters of 1-to-5 colonic crypts was reattempted as described above. Based on the iteration of filtration and manual agitation steps, up to three additional fractions of crypt suspensions were collected. Crypt suspensions were washed 3x with DPBS supplemented with antibiotics, with each centrifugation step carried out at 100g for 3 min at room temperature. Based on visual inspection under an inverted microscope, one or more crypt suspensions were selected for subsequent processing according to the size and integrity of the crypts, and either processed separately or pooled together if individual suspensions were assessed to have low-crypt content.

For both tumor and normal tissue, if blood traces were visible under an inverted microscope, the cell pellet was resuspended in 1-5 ml ACK lysis buffer (Lonza), according to the pellet size, and incubated for 5 minutes at room temperature. Quenching was performed with 3x volumes of DPBS supplemented with antibiotics, followed by an additional wash to remove ACK traces. The resulting cell pellet was further processed for either scRNA-seq, organoid generation, or both. Tissue processing protocols were extensively and iteratively optimized to maximize retrieval of high quality (low mitochondrial and ribosomal content) viable single cell suspensions for downstream analyses.

#### Single-cell RNA sequencing

Cell suspensions were filtered through a 40 μm cell strainer and incubated in FACS buffer (10 mM HEPES, 0.1 mM EDTA, 0.1% FBS) with DAPI (1 ug/ml, Thermo Scientific) and calcein AM (Invitrogen) for 5 min on ice. Viable (calcein-positive) cells were sorted using a 130 μm nozzle (SH800S SONY sorter) and collected in DPBS with 0.04% bovine serum albumin (BSA). Single-cell RNA-seq was performed on a Chromium instrument (10x Genomics) following the 3′ RNA v3.1 user manual. In brief, FACS-sorted cells were washed once with DPBS containing 0.04% BSA and resuspended to a final concentration of 700–1,300 cells/μl. Cell viability was above 80%, as confirmed with 0.2% (w/v) Trypan Blue staining (Countess II). Cells were captured in droplets and subjected to reverse transcription and cell barcoding; emulsions were then broken and cDNA was purified using Dynabeads MyOne SILANE followed by PCR amplification per manual instructions. Up to 10,000 cells were targeted for each sample. Final libraries were sequenced on the Illumina NovaSeq S4 platform (R1 – 28 cycles, i7 – 8 cycles, R2 – 90 cycles).

#### Organoid generation and culture

Primary and metastatic CRC and normal colon organoid lines were established as previously described^14,72,73^. Cells processed as described above were centrifuged at 600*g* for 5 min at 4°C and resuspended at 2,000 cells per 40 μl of Matrigel. After Matrigel domes solidified at 37°C, HISC media supplemented with Y-27632 was added to wells. Organoids were passaged every 7–10 days, and were considered established after 3 passages. For non-tumor organoid culture, HISC media was supplemented with human R-spondin 1 (1 μg/ml; Peprotech), and NGS-WNT (0.5 M, ImmunePrecise N000). Media was changed every 3–4 days. Organoid lines were expanded and early passage stock vials cryopreserved in liquid nitrogen.

For validation, organoids underwent targeted exome sequencing via MSK-IMPACT^69^ and key oncogenic genomic alterations were identified by OncoKB^74^ (see below). Diagnostic tissue from originating tumors was sequenced to confirm that these alterations were conserved in each derived organoid line. Organoids were STR-verified at the time of establishment and before every experiment, and were routinely tested for mycoplasma contamination (MycoALERT PLUS detection kit, Lonza).

#### MSK-IMPACT

Tumor and organoid targeted exon sequencing was performed using MSK-IMPACT^69^. The OncoKB precision oncology knowledgebase, an FDA-recognized human genetic variant database curated by experts at MSK^74^, was used to distinguish between oncogenic alterations (presumed “drivers”) and variants of unknown significance (presumed “passengers”). Only somatic alterations labeled as “oncogenic”, “likely oncogenic”, or “predicted oncogenic” by OncoKB were included for analysis. The MSK-IMPACT data analysis pipeline can be found at https://github.com/rhshah/IMPACT-Pipeline. Genomic alterations were annotated with information from OncoKB using the OncoKB annotator tool (https://github.com/oncokb/oncokb-annotator).

#### FACETS

Copy number alterations in solid tumors were computed from MSK-IMPACT using the FACETS (Fraction and Allele-Specific Copy Number Estimates from Tumor Sequencing) algorithm^70^, which provides allele-specific copy number estimates at the level of both gene and chromosome arm. FACETS was also used to generate purity-corrected segmentation files, for detection of whole-genome duplication events, to infer the clonality of somatic mutations, to assess arm-level copy-number changes and to generate mutant allele copy-number estimates.

### Computational Data Analysis

#### scRNA-seq data pre-processing

##### Alignment of sequencing reads

All scRNA-seq datasets were pre-processed as follows: FASTQ files from patient samples were processed with the SEQC pipeline^75^ using the hg38 human genome reference, default parameters, and platform set to 10x Genomics v3 3’ scRNA-seq kit. The SEQC pipeline performs read de-multiplexing, alignment, and UMI and cell barcode correction, producing a preliminary count matrix of cells by unique transcripts. By default, the pipeline will remove putative empty droplets and poor-quality cells based on 1) the total number of transcripts per cell (cell library size); 2) the average number of reads per molecule (cell coverage); 3) mitochondrial RNA content; and 4) the ratio of the number of unique genes to library size (cell library complexity). However, due to the sensitivity of the colorectal epithelium to dissociation, we observed increased indicators of cell stress, apoptosis, and droplet contamination in many samples, including high mitochondrial and ambient RNA expression, which can obscure statistical inference from meaningful biological gene expression. As such, typical ad hoc cell filtering based on identifying a steep dropoff in the number of transcripts per droplet (that is, a deviation leading to a ‘plateau’ in ambient RNA levels), could impair the extraction of meaningful biology. We therefore sought to systematically evaluate and correct for ambient RNA expression and filter for real single cells using CellBender^76^ as described below.

##### CellBender to subtract ambient RNA

CellBender is an unsupervised method for removing ambient RNA from single-cell RNA-seq data. It first infers rates of ambient RNA and barcode swapping per gene and droplet, respectively, from an unfiltered cell-by- gene count matrix. This probabilistic model is then used to generate a denoised (i.e., ambient RNA-corrected) count matrix, as well as the probability that each droplet contains a cell, which can be used for calling real cells. We ran CellBender on the unfiltered counts matrix of each sample produced by SEQC with the following parameters: 1) set the expected number of cells as the number of cells loaded into each 10x Chromium lane per sample (typically 5000-10000 cells), 2) set total droplets used to estimate ambient background RNA to 30,000, and 3) set training epochs to 100. We used the denoised count matrix produced by CellBender for all subsequent analyses.

##### Removing low quality cells

Based on CellBender-corrected expression counts, we sought to identify and filter out low quality cells from downstream analysis. Since our study focuses on epithelial cells, which are known to be more sensitive to single-cell dissociation than other cell types, we paid special attention to droplet quality by performing three filtering steps:

Step 1. Remove all droplets with posterior probability of containing cells ≦0.5 according to CellBender. This lenient filtering ensured that no biologically relevant cells were removed, at the cost of retaining some cells with worse technical characteristics.

Step 2. Remove droplets with <200 total counts, <200 total genes expressed, or whose libraries comprised >50% mitochondrial RNA.

Step 3. Iterative rounds of clustering and filtering to remove low quality or apoptotic cells that group together to create unstructured “junk” clusters. We carried out this filtration by combining count matrices from all patients for each sample type (non-tumor, primary tumor, and metastasis), clustering cells using PhenoGraph^22^ (*k* = 20), and studying the covariance structure of highly expressed genes within each cluster. We reasoned that highly expressed genes are not co-regulated in cells undergoing apoptosis, nor in droplets containing ambient RNA, motivating the removal of droplets lacking meaningful covariance structure. We repeated the process of clustering and filtering until only cells residing in structured clusters remained, after which all datasets were combined.

Cells passing all above criteria were retained for downstream analysis.

#### scRNA-seq data analysis

##### Data normalization and dimensionality reduction

Raw count matrices containing pre-processed data were normalized to the median library size and log- transformed with a base of *e* and pseudocount of 0.1. We then selected highly variable genes (HVGs) using the *highly_variable_genes* function in Scanpy and *flavor* = seurat_v3 (we chose *bins* = 40). We kept the top 50 genes within each bin, for a total of 2000 HVGs. In addition, for all datasets (besides the fetal colon dataset) we included genes with known relevance to normal colonic cell types (41 genes) and cell states associated with inflammatory disease, injury of the colon, and regulation of REST and EMT (56 genes) (see **Supplementary Table 1a** for complete list of genes). Going forward, we will denote the 2097 genes including both HVGs and manual additions as HVGs. Next, we performed principal component analysis (PCA) on log- normalized matrices using only HVGs and retained the number of principal components (PCs) that explain 75% of variance (112 PCs).

##### Data visualization

For all two-dimensional embeddings, we used the Scanpy *neighbors* function to compute a *k*-nearest neighbor graph on the PCs based on Euclidean distance and *k* = 30. To visualize the global CRC cell atlas (**Fig. 1d-f, Extended Data Fig. 2b**,e), the non-tumor epithelium (**Extended Data Fig. 2f**) and human fetal gut cell atlas (**Fig. 4c**), we generated projections using the UMAP implementation in Scanpy, with *min_dist* = 0.3–0.5, and *init_pos* = paga. To visualize epithelial cell subsets including all untreated epithelial cells (**Fig. 2a,c**), all tumor cells (**Fig. 3b, Extended Data Fig. 5a**,b), and patient KG146 cells (**Fig. 4a, Extended Data Fig. 7f**,g) we utilized force-directed layouts, which provide a more intuitive representation of cell state transitions and the local relationships between subpopulations, using Scanpy with the ForceAtlas2 layout and *init_pos* = paga.

##### Gene expression denoising and imputation

We applied MAGIC imputation^77^ to normalized, log-transformed count matrices to denoise and recover missing transcript counts due to dropout. Imputation was performed using conservative parameters (*t* = 3, *ka* = 5, *k* = 15). Imputed values are used for visualization of gene expression or gene signature expression (described in text and figure legends where used), as well as for analyzing mixed-lineage gene correlations in untreated patient tumors (section “*Gene correlations in normal intestine and untreated tumor*”).

##### Gene signature scores

To generate all gene signature scores in our study, we utilized the Scanpy *score_genes* function, which calculates the mean expression of genes of interest subtracted by the mean expression of a random, expression-matched set of reference genes. To account for expression-level differences across genes within signatures, we provided *z*-normalized expression data as input for this function.

#### Cell annotation

##### Partitioning cells into epithelial, stromal, and immune compartments

We clustered our combined dataset of all cells using PhenoGraph with the Louvain algorithm (*k* = 45) on the PCs obtained above. To ensure robustness to the choice of *k*, we repeated PhenoGraph clustering for all values of *k* between 20 and 100 in increments of 5 and calculated the adjusted Rand index between each pair of clusterings. We chose a *k* value that, within small variation, generated a Rand index > 0.9, indicating that cell assignments to clusters remain mostly unchanged and are thus robust to the choice of *k*.

Next, we partitioned clusters into epithelial, stromal and immune compartments based on marker gene expression (**Extended Data Fig. 2a**,b). Specifically, we used the *score_genes* function in Scanpy^78^ to score expression of compartment-specific gene signatures from Smilie et al.^79^ similar to the strategy used in that study (see **Supplementary Table 1b** for signatures for each compartment). Each cluster was assigned to the compartment with the maximal score.

##### Analysis of epithelial compartment

We filtered the epithelial compartment to discard any remaining low-quality cells, by removing the lowest modes in the distributions of log library-size, log number of genes expressed, and fraction mitochondrial RNA (see **Supplementary Table 1c** for exact thresholds) resulting in 67,534 epithelial cells. We chose to assign thresholds separately for each compartment due to their different sensitivities to dissociation and sample preparation, as well as inherent biological differences between compartments, e.g., tumor cells often have very large library sizes compared to immune or stromal cells.

Within the epithelial compartment, we then recomputed HVGs (2,097 HVGs), re-performed PCA (210 PCs), and clustered cells with PhenoGraph (k=30) and removed 4 remaining outlier clusters containing cells belonging to patients KG103, KG105, and KG66. These clusters were characterized by very low library size; little block structure in their gene covariance matrices, primarily containing mitochondrial and ferroptosis- associated genes; and a strong overlap between cells originating from the patients’ non-tumor, primary, and metastatic samples. In line with our scRNA-seq data, we observed very aberrant mucosa in histological images of non-tumor samples for these patients, noting an association with prior disease conditions, which may explain the poor sample quality. Together, these observations suggested that the clusters likely represent highly stressed or dying, disease-associated cells which would not be informative to our study. After removing them, 47,437 cells remained; all downstream analysis of the epithelial compartment was performed on these cells.

##### Tumor cell identification using single-cell CNA calls

We identified cancer cells in the epithelial compartment (**Extended Data Fig. 2c**-e) using the following criteria: 1) evidence of copy number alterations (CNAs) compared to cells originating from non-tumor colon samples, and 2) clustering that is distinct from non-tumor epithelial cells.

We identified CNAs at the single-cell level with infercnvpy, a Python implementation of InferCNV^23^ using a sliding window of 200 genes and default parameters. Mean expression of the reference diploid was determined using all available normal tumor-adjacent samples. InferCNV assumes that gene expression is proportional to copy number; thus, CNA inference can be sensitive to the choice of reference dataset, particularly if expression in the reference is sparse (e.g., due to few cells, giving noisy results) or overrepresented by a few phenotypes (e.g., a single cell type, yielding false positives). Since non-tumor epithelial cell types are differentially distributed across patients in our matched adjacent non-tumor samples, we chose to use all non-tumor epithelial data as a reference.

We performed Leiden clustering on the inferred copy number matrix and called cancer cell clusters if they harbored <25% normal tumor-adjacent cells and an average CNA score ≧1 s.d. from the diploid mean. As a result, 3102 cells derived from tumor samples with no CNA were reclassified as non-tumor epithelial cells (we perform cell type annotation on these cells in the section below), and 26,145 cells derived from tumor samples with CNA remained classified as tumor cells. We also produced an independent CNA estimate using the FACETS pipeline^70^ for samples from patients with targeted DNA panel sequencing data. In many cases, patient CNA estimates by FACETS were consistent with the most abundant single-cell CNA profiles for the same patients (see **Extended Data Fig. 2c**,d for an example).

##### Cell type annotation in the non-tumor epithelial compartment

To annotate epithelial cell types, we retained the subset of normal epithelial cells (21,297 cells), computed a PC representation (249 PCs) of the resulting log-normalized count matrix, and clustered cells with PhenoGraph using Leiden and *k* = 15 on the obtained PCs. We ensured robustness to the choice of *k* as described above. This process resulted in 48 clusters of cells, which were annotated into cell type based on two criteria: 1) similarity of mean *z*-normalized expression to that of canonical marker genes for major colon epithelial cell types (**Extended Data Fig. 2g**), and 2) gene set enrichment analysis (GSEA) of relevant cell type gene sets from the literature (**Supplementary Table 1d**), based on differentially expressed genes in each cluster compared to the rest, computed using the R package MAST^80^. GSEA was performed using the Python package gseapy^81^ with 10000 permutations and default parameters. Based on both criteria, we manually annotated the clusters as ISC (2,580 cells), absorptive precursor (6,874 cells), enterocyte (2,115 cells), BEST4^+^ enterocyte (1,367 cells), secretory precursor (5,573 cells), goblet (1,751 cells), tuft (848 cells), and enteroendocrine (189 cells). These cell type annotations are shown in **Fig. 1d** and **Extended Data Fig. 3f**.

#### Comparison of normal ISCs and treatment-naive tumors

##### Creation of an ISC-specific gene signature

To determine ISC-specific marker genes, we performed one-versus-rest differential expression analysis of ISC cells against all other non-tumor cells using MAST on the normalized, log-transformed count matrix for non-tumor epithelial cells, and calculated gene rankings according to the *-log(p-value)*logFoldChange* value for each gene (**Supplementary Table 1e**). The final ISC gene signature consisted of the top 100 ranked differentially expressed genes (*p-value* < 0.01 for all genes included). All gene rankings and list of ISC signature genes are listed in **Supplementary Table 1e**. We calculated a gene signature score on the *z*- normalized expression of these genes using the *score_genes* function in Scanpy (**Fig. 2c**).

##### Principal Component Analysis and Annotation of PC1

We took a subset of the single cell dataset containing all untreated normal and tumor epithelial cells (13,935 cells) and performed PCA on the log-normalized expression matrix. We focused on the first principal component (PC1), which we note was responsible for 13.5% of the variance in the dataset (compared to 8.96% of variance described by PC2). To annotate PC1, we ordered all genes according to their feature loadings on PC1, excluding genes with non-zero loadings which cannot be ordered. Using this ordering, we performed GSEA using the *prerank* function of the Python package gseapy with relevant cell type gene sets from the literature (**Supplementary Table 1d**) and default parameters as inputs. GSEA results can be found in **Supplementary Table 1f**.

##### DEG and GSEA analysis between untreated tumor and ISC cells

We performed differential expression analysis of ISC cells against all untreated tumor cells using MAST (**Supplementary Table 1g**) and GSEA using relevant cell type gene sets from the literature (**Supplementary Table 1d**) as well as all Hallmark^82^ and KEGG^83^ gene sets. GSEA was performed using the *prerank* function of the Python package gseapy with 10000 permutations and default parameters (**Supplementary Table 1h**).

##### Identification of ISC phenotypic admixture in treatment-naive tumors

To assess whether the cell type promiscuity observed at the cluster level (**Extended Data Fig. 3b**) is also present in individual cells, we used MAST^80^ to compute differentially expressed genes (DEGs) enriched in ISCs, enterocytes, and goblet cells in the non-tumor epithelial dataset; ranked each DEG by *-log(p- value)*logFoldChange* value; and used the top 300 genes as lineage markers. For enterocytes and goblet cells, DEG analysis was restricted to differentiated cell types (that is, excluding precursor cell types). This allowed genes shared between precursor and differentiated cell types of the same lineage to be recovered by DEG analysis. For example, the expression of enterocyte marker *SLC26A3* in late-absorptive precursor cells lowered its observed differential expression in enterocytes compared to all other cells when absorptive precursors were included in DEG analysis, despite abundant SLC26A3 expression among enterocytes. All DEG results are reported in **Supplementary Table 1e,i,j**.

Because tumor cells do not emulate the complete phenotype of normal differentiated cell types^84^, we restricted the 300 cell lineage markers from above to genes which are also abundantly expressed in more than 20% primary tumor or 20% metastasis cells. We consider a lineage marker “abundantly expressed” in a cell if its normalized expression in that cell was greater than or equal to the bottom quartile of expression from the lineage for which it is a marker. To ensure the specificity of markers, we also removed any gene which is abundantly expressed in other lineages. Finally, we determined the fraction of the remaining lineage markers which are abundantly expressed in each normal cell and treatment-naive tumor cell. The distributions for each cell type and tumor type are visualized using the *sns.kdeplot* function in Python Seaborn in **Extended Data Fig. 3c**-e.

##### Gene correlations in normal intestine and untreated tumor

To understand whether the co-expression of conflicting cell type markers (**Extended Data** Figure 3c-e) was associated with the loss of cell-type-specific gene regulation in tumors, we first computed Pearson correlations between all pairs of top ranking DEGs for ISCs, enterocytes, and goblet cells as described above (300 genes total) using the imputed expression matrix for (i) all non-tumor epithelial cells and (ii) all treatment-naive tumor cells (**Fig. 2e**). To account for heterogeneity in gene regulation across tumors, correlations were first computed for each patient tumor separately, resulting in 12 correlation matrices, and averaged. In this way, the observed strong correlations correspond to those gene pairs with strong positive or negative correlations across multiple patient tumors, suggesting consistent gene dysregulation compared to the non-tumor setting.

#### Identification of informative gene modules in CRC tumor data

We chose to apply HotSpot^34^, to identify shared gene modules in our patient tumor dataset. HotSpot is an algorithm for identifying informative gene modules within single cell datasets given a user-provided metric for local cell-cell similarity. Specifically, Hotspot finds genes with high local autocorrelation, evaluating the pairwise local correlation between genes on the local neighborhoods of cells along the cell-cell similarity graph (constructed from the k-nearest neighbor graph). Importantly, the way significant local autocorrelation is detected is suited to scRNA-seq and resilient to issues such as gene drop-out in individual cells. The computed gene-gene affinity matrix is clustered to output a set of gene modules. Unlike *global* measures of correlation (e.g., Pearson’s correlation) which presume relationships between features are consistent across a single-cell dataset, HotSpot’s *local* correlation measures are computed on the local neighborhoods in the cell knn-graph.

While previous work^24,35^ used NMF based factor analysis, Hotspot was better suited for our goals. NMF is a linear decomposition that requires a consistent and complete decomposition of the entire dataset and can be sensitive to batch and other variation. Hotspot is based on gene-gene covariance, which better represents a set of genes that work together towards a common function. Moreover, covariance has been shown to be robust to batch effects^75^ and is likely more robust to inter-tumor variation. Most importantly, Hotspot gene modules are based on covariance that can be localized to a cell subpopulation or exhibit potentially nonlinear graded expression over regions of the manifold. In cancer in particular, heterogeneous cell states adapted to different patient tumor contexts, such as metastatic sites, are likely to correspond to differences in gene covariance. Additionally, Hotspot is well suited to handle both gene pleiotropy and rare populations, which also play important roles in tumor context. For this reason, we believed HotSpot would be adept at characterizing sources of inter- and intra-tumor phenotypic heterogeneity in our dataset alongside global, manifold-defining modules of genes.

To apply Hotspot, we first partitioned the data to only include tumor cells (26145 cells), their top 2000 HVGs, and genes of known biological relevance, for a total of 2097 genes (see “Data normalization and dimensionality reduction” above). After re-normalization, we performed PCA and retained enough PCs to explain 75% of the variance (233 PCs). We then identified a subset of significantly autocorrelated features with respect to the PC latent space, by running HotSpot using the depth-adjusted negative binomial (danb) observation model and 30 neighbors. The danb model^85^ comprises the background null distribution against which expression counts are normalized, and is used to avoid flagging genes as significant due to local autocorrelation in the library size of cells.

We retained 2,003 genes with a false discovery rate (FDR) <0.01 for calculating local correlations and downstream clustering. We used the *create_modules* function in HotSpot (default parameters except *minimum_gene_threshold =* 20 and *core only* = False) to obtain a preliminary set of 37 co-varying groups of genes, called modules. Module assignments of genes can be found in **Supplementary Table 1k**. Further, below we provide further details on this clustering procedure and justify our choice of parameters.

##### Robustness of HotSpot gene modules

We evaluated the robustness of the HotSpot analysis to the number of HVGs used as input features, based on 1) consistency of gene autocorrelation, and 2) consistency of obtained modules.

Consistency of gene autocorrelation to evaluate the input cell-cell similarity matrix faithfully captures the structure of the data across different input features (genes): Given an input gene set, HotSpot removes genes with low autocorrelation along the kNN graph. This ensures that only genes which vary along the manifold in an informative manner are selected for module detection. To ensure that genes passing this criterion are robust to the number of selected HVG, we recomputed modules with 1,000-5,000 HVGs in increments of 500 genes, keeping all other parameters constant. For each combination, we calculated the difference in HotSpot local autocorrelations for each gene against its autocorrelation score when we input 2,000 HVGs (the value used in this study) and visualized the average as a boxplot (**Extended Data Fig. 4c**). These differences were minimal across HotSpot runs for all HVG pairs (maximum difference of 0.07), suggesting the cell similarity graph retained its structure regardless of how many HVGs were chosen.

Consistency of modules: To verify that the set of genes comprising each module was robust to the number of features, we generated HotSpot modules for 1,500–2,500 HVGs in increments of 50 genes, with *minimum_gene_threshold* set to 20 and *core_only* set to false. We then calculated Pearson correlations between the module scores of each set of HotSpot modules and the set obtained with 2,000 HVGs, as used in this study (**Extended Data Fig. 4d**). For each original HotSpot module obtained with 2,000 HVGs, we report the correlation of the module used in this study and the best-matching module – the module that is most highly-correlated with the original module in this study – in **Extended Data Fig. 4d**. In general, every set of gene features identified a subset of modules showing close correspondence to our final set of modules based on max correlation.

##### HotSpot module clustering

HotSpot clusters the gene-gene local correlation matrix into modules using an agglomerative hierarchical clustering procedure which, at each step, merges two genes/modules with the highest pairwise Z-score. Once a merged module contains more genes than a minimum threshold, it is labeled and cannot be combined with other labeled modules. The process ends when the highest pairwise Z-score between unmerged modules falls below a minimum value; at that point, all genes which are unlabeled are not assigned to any module.

In practice, we found choosing a large value for the minimum gene threshold left too many genes unassigned and created modules which, due to their high numbers of genes, were difficult to interpret. However, choosing too small of a threshold would fail to merge some intercorrelated, biologically similar modules. We opted to use a lower threshold of 20 genes minimum - to err on the side of more manageable, smaller modules - and then manually group modules with similar biological interpretations, after ensuring their genes were also correlated. This process is described in detail below.

##### HotSpot module grouping and annotation

All original 37 HotSpot modules were annotated manually based on known canonical gene markers for intestinal and non-intestinal cell types and from literature. A subset of genes used to annotate modules are shown in **Fig. 3a** and **Extended Data Fig. 4b**,h, and all genes used for annotation are highlighted in **Supplementary Table 1k**. For HotSpot modules that were more difficult to identify, we performed gene set over-representation analyses using the Python package gseapy and the GO Biological Process gene sets to give a direction to investigate. They were otherwise used to give more evidence supporting our initial annotations. All module annotations can be found in **Supplementary Table 1k**, and over-representation analysis results for our final module annotations are provided in **Supplementary Table 2**.

We chose to focus on 23 of the 37 modules (1,201 genes) which we found to represent meaningful biological gene programs. The 14 modules (722 genes) which were not explored further were annotated as cell cycle/proliferation (2 modules), cell stress (4 modules), leukocyte (3 modules), or cilia (1 module) or else could not be interpreted (4 modules). We then manually grouped 19 modules into 6 module groups after 1) arriving at the same biological interpretation for all modules within a group, and then 2) ensuring the local correlations between genes of grouped modules were high on average (**Extended Data Fig. 4a**,b). This resulted in 10 final gene modules; all module groupings and annotations are shown in **Extended Data Fig. 4a**,b**,h** and **Supplementary Table 1k**.

Once the modules were annotated and grouped using the above-described strategies, we categorized the modules into two distinct categories:

1. Canonical: Modules describing canonical intestinal cell types and processes such as epithelial differentiation, mucus production, and small molecule transport, which are critical to maintaining normal intestinal function.
2. Non-canonical: Modules describing processes not typically seen in the healthy intestine, such as keratinization, inflammatory response, and wound healing.

##### HotSpot gene module scores

HotSpot evaluates per-cell scores for each module using the following procedure: For all genes in a module, expression counts were first mean-centered using the danb null model and smoothed using the weighted average of their nearest neighbors. The background null distribution factors cell library size differences into account; thus, centering based on the null ensures that correlations are not impacted by library size differences. PCA was then performed on the resulting centered counts and the first PC values were used as the module scores for each cell. We used HotSpot module scores to visualize and summarize the patterns of module expression within cell groupings (e.g., clusters), and to relate gene modules to gene signature scores for pre-existing gene sets as well as to sample metadata (**Fig. 3,4** and **Extended Data Fig. 4**–7).

**Extended Data Fig. 4b**,h are plotted using the *dotplot* function in scanpy where the groups of cells (y-axis) include the top-scoring cells for each module (i.e., cells which are specifically high for that module). To assign cells as “top-scoring” for a HotSpot module, we z-normalize all HotSpot module scores across cells and then make sure a cell 1) has a score > 1 s.d. above the mean for that module score, and 2) has a score < 1 s.d. above the mean for all other modules not given the same annotation and assign it to the module for which it has the highest score. In **Extended Data Fig. 4b**, cells are assigned to one of the 23 ungrouped modules on which we focus. In **Extended Data Fig. 4h**, cells are assigned to one of the 10 final gene modules.

#### Axis of canonical and non-canonical phenotypes

We computed PC1 of the dataset using only the 1201 genes belonging to our 10 gene modules. We found the scores of differentiated intestinal modules were anti-correlated with PC1 and the scores of trans- differentiated cell states were positively correlated with PC1 (Spearman correlation, all p-values < 0.05; **Supplementary Table 1l**). Moreover, upon ordering modules by the positions of their highest-scoring cells along PC1, we found the ordering reflected a canonical to non-canonical spectrum of states: in order, Absorptive Intestine, Secretory Intestine, Intestine, EMT, Tumor ISC-like, Injury Repair, Osteoblast, Endoderm Development, Neuroendocrine, Squamous (peak PC1 positions in **Supplementary Table 1l**). Accordingly, we used this PC to summarize the canonical or non-canonical orientation of patient samples in our single-cell data (**Fig. 3e,f**).

#### Interpatient entropy of gene modules

We used entropy to evaluate the patient-specificity of each gene module using the following procedure: 1) To correct for imbalance among patient sample sizes in our dataset, we first re-sampled our patient tumor dataset to 10000 cells such that cells were evenly distributed among patients; 357 cells were sampled with replacement per patient. 2) For each module, we determined a subset of high-scoring cells whose module scores were greater than 1 s.d. above the mean. 3) We computed the Shannon entropy over the distribution of patients in the high-scoring subset of cells for each module. We repeated this procedure 100 times and visualized the entropy distribution with kernel density plots using the *kdeplot* function in Python Seaborn (**Extended Data Fig. 4i**).

#### Association of gene modules in single-cell data

##### Associations with clinical covariates

To test for associations between the HotSpot gene module scores in our scRNAseq data and clinical features associated with the patient samples from which the cells were collected, we performed the following procedure: 1) To control for bias owing to uneven numbers of cells collected across patients, we re-sampled our patient tumor dataset to 10000 cells such that cells were evenly distributed among patients; 357 cells were sampled with replacement per patient. 2) For binary clinical features (e.g., treated vs. untreated), we separated all cells within the re-sampled dataset into two groups based on the status of the patient from which the cells were collected. 3) We compared the HotSpot gene module scores between the two groups using the Mann Whitney U test. This procedure was repeated 100 times for each clinical covariate and the mean values taken across all 100 repetitions are shown in **Extended Data Fig. 5c (left)**.

For continuous clinical features (e.g., overall survival time), we performed step 1 from above and calculated the Pearson correlation between the feature values and HotSpot gene module scores. Mean values following 100 repetitions of this procedure are shown in **Extended Data Fig. 5c**^59^.

##### Associations with consensus molecular subtype (CMS) classifications

For CMS classification of single-cell data, we repeated the procedure as described^86^ to apply the bulk RNAseq CMSclassifier^27^ to scRNAseq data: We first obtained CMS centroid data from the CMSclassifier package, including 20 centroids (4 CMS types from 5 datasets) defined on 695 genes. We calculated Pearson’s correlation coefficient values between the 20 centroids and all tumor cells using z-normalized expression of the same 695 genes on which the centroids are defined. After calculation, each cell was annotated as the CMS type with the highest average correlation across all five datasets.

To find associations between HotSpot module scores and CMS types, we performed the same procedure as above in “*Associations with clinical covariates”*, separating cells into two groups based on whether they were annotated as a CMS type. Mean values from Mann Whitney U tests following 100 repetitions of this procedure are shown in **Extended Data Fig. 5d**.

#### Association of gene modules with clinical covariates in bulk RNA-seq data

To test the associations of HotSpot modules with clinical covariates in bulk cohorts, we performed the following general procedure: 1) We ran single-sample GSEA on two bulk RNA-seq datasets, LARC and TCGA-COAD, collected from CRC primary patient tumors using the lists of genes in our HotSpot modules as the input gene sets. 2) For each dataset, we tested the association of patients’ clinical features with the enrichment scores of their tumor samples.

##### Single-sample GSEA (ssGSEA) analysis

LARC: We analyzed 108 locally advanced rectal cancer (LARC) tumor samples with available RNA- sequencing data^46^. Genes were retained if they had a count per million (CPM) greater than 1 across more than 50% of the samples were retained. The edgeR v3.40.2 package^87^ was used for TMM normalization and FPKM transformation and the org.Hs.eg.db v3.16.0 package was used for gene annotation. Genes with multi mapped Ensembl IDs or that mapped to multiple Ensembl IDs were removed. We then performed ssGSEA analysis using the R package hacksig v0.1.2 R package^44^ and all HotSpot modules.

TCGA: We downloaded and analyzed RNA- sequencing data for 445 tumor samples from the TCGA-COAD study^45^. RNA raw counts were retrieved using TCGAbiolinks_2.26.0^88^ and genes with a count of 0 across all the samples were removed, as well as genes that had multiple associated gene symbols or no gene symbol. The VST transformation was performed using the DESeq2_1.38.3 package^89^. Subsequently, ssGSEA analysis was conducted utilizing the R package GSVA_1.46.0^90^.

##### Associations with clinical covariates

For binary clinical features (e.g., disease-free survival), we separated the enrichment scores into two groups based on the status of the patient from which the bulk RNAseq data was collected and compared the enrichment scores for each HotSpot module between the two groups using the Mann Whitney U test. For continuous features, we calculated the Pearson correlations between the enrichment scores of each HotSpot module and feature values. Results are shown in **Fig. 3g,h, Extended Data Fig. 5e**,f.

##### Kaplan-Meier analyses in TCGA-COAD cohort

For each HotSpot module, we collected two groups of samples from the TCGA-COAD cohort: 1) highly enriched samples with ssGSEA enrichment scores greater than 1 s.d. above the mean enrichment score among all samples, and 2) lowly-enriched samples with enrichment scores less than 1 s.d. below the mean. We then performed a log-rank test on disease-free survival between these groups using the *lifelines* package in Python^91^ Survival curves are shown in **Extended Data Fig. 5g** for all modules with significant results (p-value < 0.05).

#### Creation of a human fetal colon progenitor gene signature

We used a gut cell atlas containing scRNA-seq data from dissected human embryos aged 6.1 to 17 weeks, which captures the development of human intestinal cells from a fetal progenitor state to differentiated crypts^50^. We downloaded a raw H5AD file from the authors containing all epithelial cells from fetal donors and limited our analysis to 8,408 cells originating from the large intestine of first and second trimester samples.

##### Data reprocessing

Because the large intestine cells represent a minority of the fetal gut cell atlas (8,408 of 52,184 total cells), we reprocessed the partitioned dataset to focus our analysis on these cells specifically. We ran HVG selection (2,000 HVGs), PCA (167 PCs explaining 75% variance), and UMAP projection (*min_dist* = 0.5) using scanpy as described in the sections, “*Data normalization and dimensionality reduction*” and “*Data visualization”*. Upon inspection, UMAP projection revealed that the week 11.1 sample separated from all others. Cells from this sample were characterized by heat-shock genes *DNAJB1*, *HSP90AA1*, *HSPE1*, *HSPA8*, *HSPA1A* among the top 10 DEGs (compared to all other samples, by MAST analysis) indicating cell stress, so it was removed resulting in 7,984 cells.

In the remaining dataset, we retained the authors’ original cell type annotations, but merged the enteroendocrine subtypes (M/X, D, β, L, N, K, I, and enterochromaffin cells) into one enteroendocrine group. According to the authors’ annotations, we found the first trimester samples predominantly consisted of progenitor cells, with proximal progenitor, distal progenitor, and stem cells comprising 88% of the cells. In contrast, second trimester samples exclusively consisted of mature colon mucosal cell types and exhibited strong expression of *LGR5*, *TFF3*, *SLC26A3*, *NEUROD1*, and *POU2F3* (corresponding to ISCs, goblet cells, mature enterocytes, enteroendocrine, and tuft cells, respectively). Accordingly, we reasoned that the separation between first and second trimester samples sufficiently captured the distinction between progenitor-like cell types and the formation of colonic crypts.

To determine marker genes specific to the first trimester cell population, we first performed a differential expression analysis of first versus second trimester cells using MAST on the normalized, log-transformed count matrix and identified 173 DEGs with logFC > 2 and adjusted *p*-value < 1e-5. Earlier cells are more proliferative, therefore we removed genes related to cell cycle or proliferation from our first-trimester gene list. Specifically, we calculated the Pearson correlation of all first trimester DEGs with 445 genes belonging to the Reactome “Cell Cycle, Mitotic” and “Cell Cycle, G1-G1/S Phase” gene sets and the Hallmark “Cell Cycle, G2M Checkpoint” gene set^92^. 60 genes with a correlation greater than 0.25 for at least one gene belonging to the gene sets were removed. Our fetal gene signature comprises the remaining 113 genes, which are reported in **Supplementary Table 1m**.

##### Comparison between fetal signature and cancer fetal like state

From the differential expression analysis of first versus second trimester cells described above, we selected a set of 1712 genes upregulated in the first trimester colon (logFC > 0.5 and adjusted p-value < 1e-5) and compared the expression of these genes to the expression of our fetal signature in patient tumors. For the datasets of patients KG146, KG182, KG183, and KG150, we calculated the Pearson correlation between a gene signature score calculated using our fetal gene signature (see section “*Gene signature scores*” for details on calculating gene scores) and the imputed expression of each gene. In **Supplementary Table 1n**, we report the logFC from DE analysis as well as the average correlation for 193 genes with consistent trends (abs(R) > 0.25, positive or negative in all patients, p-value < 0.05) in at least three of four patients.

#### Alignment of canonical to non-canonical tumor axes across patients

##### Identification of canonical-non-canonical axes in multiple patient datasets

While the canonical to non-canonical states globally defined the main axis of variation in our dataset across patients (**Fig. 3e,f**), we performed a more detailed comparison of this transition in the 4 patients most progressed towards non-canonical states, KG146, KG182, KG183, and KG150. To best capture a faithful trajectory within each individual patient, we reprocessed each patient individually. Each complete patient tumor dataset, consisting of primary tumor, synchronous metastatic tumor, and metachronous metastatic tumor samples, was subset and separately processed as in the section “Dataset normalization, imputation, and dimensionality reduction”.

To establish the patient tumor datasets’ canonical-non-canonical axes, we reasoned DCs would best capture the nonlinear transformation from canonical to non-canonical states in one dimension in our patient datasets, computing this DC independently for each patient to avoid any possibility of artificial imposition of the same ordering in other patients, e.g., wherein one patient dataset assumes the structure of another due to a disparity in size between datasets.

We computed diffusion maps (*knn* = 30) for each patient and retained a subset of DCs based on the eigengap of the ranked components’ eigenvalues for the first 20 DCs (KG146, 4; KG182, 6; KG183, 8; KG150, 4 DCs). For each patient dataset, we ranked its DCs by the difference between the average Spearman correlation of a DC with all non-canonical modules and the average Spearman correlation with all canonical modules and selected the DC with the highest difference between the two averages as the axis for canonical to non- canonical transformation. For patients KG146, KG182, and KG150, the first DC was selected, while the fourth DC was selected for patient KG183. The consistent and independent selection of the first DC as the canonical to non-canonical transition in 3/4 patients further supports the importance of this axis. We note that patient KG183 had fewer non-canonical cells compared to the other patients and therefore this transition was not the top DC as in other cases.

##### Alignment across patients by dynamic time warping

To directly compare DCs across patients, we sought to align all patient DCs to a common reference axis, selecting DC1 from patient KG146 which spans the greatest phenotypic range (**Figure 3e)**. We assume that each patient axis represents a sequential transformation of cell states and require any alignment to preserve this per-patient pseudo-temporal ordering of states. However, the alignment must also account for different starting and end states due to different extent of progression into non-canonical states, as well as different durations for each state.

Therefore, we chose to use dynamic time warping (DTW), which has previously been used in single cell data analysis for trajectory and sample alignment^53–55^. DTW is a time-series analysis algorithm used to measure the similarity between two temporal sequences which may vary in speed, start and end points, while conserving temporal order. In DTW, the sequences are warped non-linearly in the time dimension to produce the best match (smallest absolute difference) between sequence values, subject to the constraint that the warping function must be monotonically increasing. DTW is very similar to the Needleman-Wunsch algorithm and nearly a special case of the Wasserstein metric from optimal transport^93^.

##### Choosing features for alignment across patients

We chose to focus on features relating to our HotSpot modules for a more faithful alignment of the canonical to non-canonical axis across patients as these represent the gene programs which we would like to best track across trajectories. Specifically, we chose to align DCs using the HotSpot module scores for our canonical and non-canonical HotSpot modules, rather than individual gene expression. Aligning based on module scores had several advantages over aligning over individual genes. First, focusing on 10 modules allows for more robust computation than an alignment over ∼1,000 and moreover is not sensitive to the number of genes in the module. Second, using gene module scores “denoises” the trends in gene expression on which we’re aligning and allows us to incorporate much more information (e.g., multiple gene programs) captured by a small set of features (10 gene module scores). Finally, this approach is less sensitive to inter-patient differences, considering independently evolved tumors, which were previously observed as biologically similar gene programs across patients before grouping HotSpot modules (**Extended Data Fig. 4ab**). We noticed that gene covariance across these similar modules is better conserved than individual gene trends.

We used DTW to warp the patient DCs to optimally match DC1 of patient KG146 based on trends in gene module scores between patients. Trends were calculated as in “Visualization of Expression Trends” on gene module scores and z-normalized to be comparable across patient datasets. We then used the DTAIDistance library in Python to compute n-dimensional warping paths between DCs (where n is the number of features considered). This version implements DTW with prefix- and suffix- invariant (psi-) relaxation, which allows start and end points of the sequence to be ignored (i.e., partial matches between sequences) if this would lead to a lower cost; we used a psi-relaxation value of 100 (out of 500 DC “timepoints” representing the x-axis of the gene trends). After computing the optimal warping path, we performed linear interpolation between the original and warped DC trend values using the “interp1d” function from the Python package SciPy to construct a continuous function and warped all patient DC values to their aligned value in DC1 of KG146 using this function.

##### Robustness of DTW alignments

We performed a leave-one-out analysis to ensure no individual gene module score had a disproportionate influence on the alignment result. For each of the 10 gene modules, we recomputed the optimal warping path of each patient DC and DC1 of patient KG146 using only the other 9 gene modules. We then computed the mean absolute error (MAE) and Pearson correlation of each *leave-one-out* warping path (the re-mapped values of each patient DC onto DC1 of patient KG146 with one module score excluded) compared to the path obtained using all features (the re-mapped values of each patient DC onto DC1 of patient KG146 with all modules). Because psi-relaxation could result in mismatches between the start and end points of the sequences, MAEs and correlations were only calculated on timepoints shared by both sequences. For all patients and gene modules, the absence of any gene module gave rise to a maximum error of 0.050 (**Extended Data Fig. 6c top**) out of 1.0, where 1.0 would indicate an average error at each timepoint equal to the length of DC 1. Additionally, we observed a minimum correlation of 0.973 (**Extended Data Fig. 6c bottom**), indicating strong similarity between all *leave-one-out* warping paths and the one used in this study.

##### Visualization of module trends

We use generalized additive models (GAMs) with cubic splines as smoothing functions as in Palantir^54^ to analyze module score trends along original DC (KG146) or DTW-warped DC (KG182, KG150, KG183) axes (**Fig. 4b,d-f**). GAMs increase robustness and reduce sensitivity to density differences, and cubic splines are effective in capturing non-linear relationships. In our work, trends for a module score were fitted using a regression model on the DC values (x-axis) and module score values (y-axis). The resulting smoothed trend was derived by dividing the data into 500 equally sized bins along the DCs and predicting the module score at each bin using the regression fit. In **Fig. 4b, d-f**, module score trends are visualized from the 20th percentile value (white) to the maximum value (highest saturation).

#### Palantir pseudotime and branch calculations

Not only was the sequential progression of canonical to non-canonical states conserved in the for most non-canonical patients (KG146, KG182, KG150 and KG183), but also all 4 patients had both squamous and neuroendocrine states. We used Palantir^54^ to investigate these two fates and genes underlying each of these fate transitions. Palantir requires a starting state as input. As output, Palantir computes the terminal fates from the data and provides a cell fate map that assigns a probability for each cell to differentiate into each terminal fate. Additionally, Palantir also outputs a pseudotime alignment of cells from start to each of the terminal states and can provide branching gene trends leading to each terminal fate. Palantir was run separately on the tumor datasets of patients KG146, KG182, and KG150, excluding KG183 because of the limited number of non-canonical cells in this patient. We provided Palantir with LGR5^+^ cells (cells with the highest imputed expression of LGR5) as the input starting point motivated by their identification as cells of origin in CRC in other studies. Moreover, we note Palantir has been shown to be robust to the exact choice of starting cell. We ran Palantir with 500 waypoints and a number of DCs chosen by eigengap to identify trajectories originating from the LGR5^+^ starting position. In all three patients, we discovered two distinct branches that led to CDX2^−^, non-intestinal terminal cells. We disregarded three additional branches in patients KG146 and KG150 as the terminal cells for these branches were CDX2^+^ and showed expression of various markers of the differentiated intestine, including FABP1 and TFF3, suggesting these represented trajectories toward canonical states.

To annotate the two non-canonical terminal states identified by Palantir, we compiled known markers for squamous or neuroendocrine cells (observed in healthy cells or non-CRC cancers, all genes listed in **Fig. 4g**), and calculated the Pearson correlations between their imputed gene expression and the probabilities for the non-intestinal branches, these correlations are shown in **Fig. 4g**. To ensure the inclusion of only cells belonging to a given branch and to avoid interference by other cell states, we excluded all cells with a probability of less than 0.5 for that branch when calculating correlations. This analysis identified branches corresponding to neuroendocrine-like and squamous-like states in each patient. To further support our annotations, we calculated Pearson correlations between the non-intestinal branch probabilities in each patient and imputed expression for all genes observed in all three patients. We found that the top 5 genes ordered by average correlation among the three squamous annotated branches in KG146, KG182, and KG150 were associated with squamous epithelium and keratinization (*DMRTA1*, *NECTIN4*, *DLX3*, *CXCL14*, *LYPD3*; all genes in **Supplementary Table 1o**). Similarly, 4 of the top 5 genes among the three neuroendocrine annotated branches were associated with glial and neural cells (*TRPM3*, *ITPR2*, *PLPPR1*, *PPFIA2*; all genes in **Supplementary Table 1p**).

##### Visualization of Palantir pseudotime trends

Palantir gene trends were calculated like in “*Visualization of module trends”* using GAMs to fit gene expression along pseudotime as determined by Palantir. All expression trends of individual genes were calculated on MAGIC imputed data (see “MAGIC imputation”), and the standard deviation of expression along each bin was determined by the standard deviation of the residuals of the fit.

##### Kaplan-Meier analyses of fetal signature in bulk data

For all samples in the LARC and TCGA-COAD cohorts, we calculated ssGSEA enrichment scores as described in “*Single-sample GSEA (ssGSEA) analysis*” using our fetal gene signature as an input. For each cohort, we then collected two groups of samples: 1) highly enriched samples with ssGSEA enrichment scores greater than 1 s.d. above the mean enrichment score among all samples, and 2) lowly enriched samples with enrichment scores less than 1 s.d. below the mean. We performed a log-rank test on disease-free survival between these groups using the *lifelines* package in Python. Survival curves are shown in **Extended Data Fig. 6e**,f.

#### Cell state classifications in KG146 patient tumors

We first isolated the KG146 primary tumor data (880 cells) and determined principal components that explain 75% of the variance (119 PCs). We then clustered the cells using PhenoGraph (*k* = 30) and identified 6 clusters. We repeated this process for the liver metastasis data (1,279 cells, 200 PCs) and obtained 9 clusters. We calculated the average HotSpot gene module scores (e.g., module scores calculated in “*HotSpot gene module scores*”) for each cluster and annotated the clusters as “ISC-like”, “Absorptive-like”, “Secretory-like”, “Fetal”, “Injury-Repair”, “Neuroendocrine-like”, and “Squamous-like” based on high scores for specific gene modules associated with respective labels (**Extended Data Fig. 7d**). We further re-classified one cluster of “ISC-like” cells as “Proliferation/ TA-like” based on their unique expression of the proliferation markers such as *MKI67* and *PCNA*. **Extended Data Fig. 7d** is plotted using the *dotplot* function in scanpy where the groups of cells (y-axis) are PhenoGraph clusters and features are HotSpot module scores. Cluster assignments are made within each sample using k=30.

#### Mapping organoid data to patient tumor

To map cells of each organoid sample to the phenotypically closest tumor cell state from the full KG146 patient dataset, we developed a manifold-based classifier that combined methods from Harmony^94^ and PhenoGraph^22^ to transfer labels between datasets. The process involved three distinct steps—feature selection, co-embedding, and classification.

##### Feature selection

We identified the top 100 DEGs for each annotated cell state in the KG146 patient tumor dataset resulting in a total of 800 genes. We then subsetted the resulting lists to only include those that are highly significant, i.e. with a log-fold change greater than 3 and Benjamini-Hochberg-adjusted *p*-value less than 0.001, resulting in 753 genes total. These genes were used for PCA and to produce neighbor graphs in the following steps.

##### Co-embedding

Using a standard co-embedding approach consisting of a joint PCA and UMAP, we observed substantial “batch” effects between the two datasets, making label transfer between similar cells ineffective. Therefore we followed the approach outlined in Garg et al.^95^ to bridge between datasets. Specifically, we first computed the nearest neighbor graph (neighbors’ function in Scanpy with k = 30) in each dataset separately. Next, we computed mutual nearest neighbors (MNNs) between the samples using Harmony^94^. Importantly, we utilized cosine metric to quantify distance between cells across samples, as this metric is less sensitive to technical artifacts and better reflects conserved biological states in both the in-vivo and in-vitro samples^96^. We chose to use a higher number of mutual neighbors (mknn=60) because it is more robust to sparsity in the MNN graph.

Next, both the within-sample nearest neighbor and between-sample mutual nearest neighbor graphs were converted to a within-sample and between-sample affinity matrix respectively using the adaptive Gaussian kernel (default parameters) as implemented in Harmony^94^. The resulting matrices were concatenated to obtain an “augmented” cell-cell affinity matrix. In summary, it consists of three components: A) similarity between cells within the in-vivo data, B) similarity between cells within the in-vitro data, C) similarity between cells across the in-vitro and in-vivo data. This matrix was input to Phenograph classification (see below) to propagate labels from the reference (KG146) dataset to the unlabeled dataset (organoid) and generate UMAP co-embeddings of patient and organoid datasets as shown in **Extended Data Fig. 7b**,c.

##### Classification

Finally, we supplied the augmented affinity matrix from Step 2 directly to PhenoGraph’s “classify” function^22^ with default parameters. Briefly, this function converts the affinity matrix into a row-normalized Markov matrix and computes the probability of random walks starting from unlabeled cells from the in-vitro samples reaching a class of labeled cells in the in-vivo sample. Finally, each unlabeled cell is assigned the cell state label with the maximum probability.

To summarize our organoid sample classifications, we aggregated the probabilities of different cell states into three generalized categories. Specifically, we summed the probabilities of ISC-like and TA/Proliferative cell states to form ISC/TA, the probabilities of absorptive-like and secretory-like states to form differentiated intestine, and the probabilities of fetal/injury repair, neuroendocrine, and squamous cell states to form non- canonical. These groupings can be interpreted as the likelihood of a cell belonging to any of the several cell states which were combined. Like in Chan et al.^84^,the resulting probabilities for each of these categories are displayed in **Extended Data Fig. 7b**,c and in the ternary plots created using the Python package python- ternary^97^ in **Fig. 5b,g**.

#### Identification of fetal-state-associated transcription factors

We aimed to generate a ranked list of human transcription factors (TFs) that are associated with the fetal progenitor state in a conserved manner across non-canonical patient tumors. To do this, we first obtained 1,099 human transcription factors^98^. We restricted our list of potential targets to TFs for which we observe 1) at least 5 UMIs in any cell in all four patient datasets (leaving 527 TFs), and 2) at least 50 TF-expressing cells in all four patient datasets (leaving 508 TFs). This was intended to remove sparsely expressed TFs which may not be as easily targeted, and to restrict ourselves to TFs whose correlations with our fetal signature are more reliable.

Next, we used Palantir to compute the expression trends for all the TFs and the fetal progenitor signature score along the canonical-to-non-canonical DC (see *Visualization of expression trends* for details on computation of expression trends). Because we are interested in TFs which drive the transition from canonical cell states into fetal cell states, we focused on those whose expression peaks in each patient along their DC prior to entry into non-canonical states. For patients KG146, KG182, KG150, and KG183, we first identified the position along their DTW-warped DC of the first maxima of the trend in the fetal progenitor signature calculated along this DC (indicated in vertical dashed lines in **Figure 6a**). The maxima were found by computing the numerical derivative of the trend and identifying the first position at which the sign of the derivative transitions from positive to negative. If no such position existed (i.e., KG150, KG183), we used the position of the maximum value. We then calculated the Pearson correlation between the expression of each TF and the fetal progenitor gene signature score using only cells at positions along a patient’s DC which precede the peak in the signature score. This yielded 4 correlation values total for each TF, one for each patient. We then focused only on TFs with a minimum correlation of at least R=0.5 in patient KG146 (leaving 14 TFs) and R=0.2 in all four patients (leaving 6 TFs).

For the remaining 6 TFs, we compared the treatment response of expression in organoids by computing the log fold change in organoids treated with irinotecan and cultured in HISC media compared to those just cultured in HISC media with no treatment. Log-fold-changes were calculated from single cell data only using cells classified as non-canonical (see section “*Mapping organoid data to patient tumor*”), and values are shown in **Extended Data Fig. 8a**.

### Multiplexed immunofluorescence

#### Multiplexed tissue staining and imaging

Primary antibody staining conditions were optimized using standard immunohistochemical staining on the Leica Bond RX automated research stainer with DAB detection (Leica Bond Polymer Refine Detection DS9800). Using 4 µm formalin-fixed, paraffin-embedded tissue sections and serial antibody titrations, the optimal antibody concentration was determined followed by transition to a seven-color multiplex assay with equivalency. Optimal primary antibody stripping conditions between rounds in the seven-color assay were performed following 1 cycle of tyramide deposition followed by heat-induced stripping (see below) and subsequent chromogenic development (Leica Bond Polymer Regine Detection DS9800) with visual inspection for chromogenic product with a light microscope (TH). Multiplex assay antibodies and conditions are described in **Table 1**.

| Antigen | Antibody Clone | Manufacturer | Concentration |
| --- | --- | --- | --- |
| HER2 | 29D8 | CST | 0.64 µg/ml |
| SOX2 | SP76 | Abcam | 1.26 µg/ml |
| CK20 | D9Z1Z | CST | 0.208 µg/ml |
| CDX2 | D11D10 | CST | 0.0575 µg/ml |
| VIM | D21H3 | CST | 0.0375 µg/ml |
| Chromogranin A | EP1030Y | Abcam | 2.705 µg/ml |

**Panel 2**
| Antigen | Antibody Clone | Manufacturer | Titration |
| --- | --- | --- | --- |
| TROP2 | SP294 | Abcam | 3.29 µg/ml |
| Ki67<br>(Culture supernatant) | SP6 | Biocare | 1:100 |
| CK5 | E2T4B | CST | 0.142 µg/ml |
| TP63 | D9L7L | CST | 0.15ug/ml |
| OLFM4 | D1E4M | CST | 0.27ug/ml |
| PLCG2 | E5U4T | CST | 0.2 ug/ml |

#### Seven-color imaging assay

4 µm FFPE tissue sections were baked for 2.5 hours at 63°C in vertical slide orientation with subsequent deparaffinization performed on the Leica Bond RX followed by 30 min of antigen retrieval with Leica Bond ER2 followed by 6 sequential cycles of staining with each round including a 30-minute combined block and primary antibody incubation (Akoya antibody diluent/block ARD1001EA) except HER2. HER2 uses one hour incubation.

For Chromogranin A and OLFM4, detection was performed using a secondary horseradish peroxidase (HRP)- conjugated polymer (Akoya Opal polymer HRP Ms + Rb ARH1001EA; 10-min incubation). Detection of all other primary antibodies was performed using goat anti-rabbit Poly HRP secondary antibody (Invitrogen B40962; 10 min incubation). The HRP-conjugated secondary antibody polymer was detected using fluorescent tyramide signal amplification using Opal dyes 520, 540, 570, 620, 650 and 690 (Akoya FP1487001KT, FP1494001KT, FP1488001KT, FP1495001KT, FP1496001KT, FP1497001KT). The covalent tyramide reaction was followed by heat induced stripping of the primary/secondary antibody complex using Akoya AR9 buffer (AR900250ML) and Leica Bond ER2 (90% AR9 and 10% ER2) at 100°C for 20 min preceding the next cycle (1 cycle of stripping for HER2, CDX2, Tp63, PLCG2 and Chromogranin A, two cycles for Sox2, CK20, Vim, Trop2, CK5, and OLFM4, 2.5 cycles of stripping for Ki67). After 6 sequential rounds of staining, sections were stained with Hoechst (Invitrogen 33342) to visualize nuclei and mounted with ProLong Gold antifade reagent mounting medium (Invitrogen P36930).

#### Multispectral imaging, spectral unmixing and cell segmentation

Seven color multiplex-stained slides were imaged using the Vectra Multispectral Imaging System version 3 (Akoya). Scanning was performed at 20x (200x final magnification). Filter cubes used for multispectral imaging were DAPI, FITC, Cy3, Texas Red and Cy5. A spectral library containing the emitted spectral peaks of the fluorophores in this study was created using the Vectra image analysis software (Akoya). Using multispectral images from single-stained slides for each marker, the spectral library was used to separate each multispectral cube into individual components (spectral unmixing) allowing for identification of the seven marker channels of interest using Inform 2.4 image analysis software. Images were exported to Indica Labs Halo image analysis platform and cell segmentation and signal thresholding was performed separately on each case using a supervised algorithm.

#### Twelve-color imaging assay

4 µm FFPE tissue sections were baked for 1 hour at 62°C in vertical slide orientation with subsequent prestaining of deparaffinization (Leica, Bond Dewax Solution, AR9222) and 35 minutes of Bond ER2 (Leica, AR9640) antigen retrieval at 100°C was performed on the Leica Bond RX automated research strainer. The Lunaphore Comet imaging chip (MK03) was placed over the acquisition region based on alignment with regions selected on H&E and then loaded into the Comet device followed by a first cycle of autofluorescent image acquiring and preceding 9 cycles of staining, imaging and elution. Each cycle contained one rabbit primary antibody and/or one mouse primary antibody detected with an Alexa Flour conjugated species- specific secondary antibody (see **Table 1**). The cyclic staining, imaging and elution was automatically performed by COMET. The optimal concentration, position in the panel and incubation time (4 minutes for each cycle except the HER2 was 8 minute incubation) of each marker in the panel was optimized such that the results of the multiplex imaging were concordant with the optimized single diaminobenzidine IHC stain for each marker which was selected by an MSK pathologist (T.H.) according to the DAB staining result which was performed on the Leica BondRX Stainer using the Leica BOND Polymer Refine Detection Kit, DS9800. For the COMET, all the antibodies were pre-diluted with Intercept Antibody Diluent (LI-COR, 927-65001). After each staining cycle, elution of primary and secondary antibodies was performed with Elution Buffer Solution (Lunaphore, BU07-L). For each sample, a 0.9 mm^2^ region of interest was imaged under the Lunaphore slide cover chip at 20x magnification, and a full channel stacked OME.tif file was generated automatically once all cycles were completed. The OME.tif image was exported after background subtraction with Lunaphore COMET Viewer software which was visualized and assessed per channel using Indica Lab Halo software.

| Antibody | Company | Clone | COMET Conc. | Secondary Ab (Invitrogen) | Cycle |
| --- | --- | --- | --- | --- | --- |
| HER2 | CST | D8F12 | 1 µg/ml | D anti-R AF647 (A32795, 10 µg/mL) | 1 |
| CDX2 | CST | D11D10 | 0.46 µg/ml | D anti-R AF647 (A32795, 10 µg/mL) | 2 |
| PROX1 | CST | D2J6J | 1.5 µg/ml | D anti-R AF647 (A32795, 10 µg/mL) | 3 |
| Ki67 | Abcam | EPR3610 | 1 µg/ml | D anti-R AF647 (A32795, 10 µg/ml) | 4 |
| Vimentin | Thermofisher | V9 | 0.33 µg/ml | D anti-M AF647 (A32787, 10 µg/ml) | 5 |
| PLCG2 | CST | E5U4T | 1 µg/ml | D anti-R AF555 (A32794, 10 µg/ml) | 5 |
| CHGA | CST | 5H7 | 0.13 µg/ml | D anti-M AF647 (A32787, 10 µg/ml) | 6 |
| CK5 | CST | E2T4B | 0.213 µg/ml | D anti-R AF555 (A32794, 10 µg/ml) | 6 |
| SYNC | ThermoFisher | Poly | 2 µg/ml | D anti-R AF647 (A32795, 10 µg/ml) | 7 |
| CK20 | CST | D9Z1Z | 0.21 µg/ml | D anti-R AF647 (A32795, 10 µg/ml) | 8 |
| GPC1 | Abcam | EPR22580-72 | 6.47 µg/ml | D anti-R AF647 (A32795, 10 µg/ml) | 9 |

### Organoid experimental methods

#### Organoid culture

For experiments, organoids were harvested from Matrigel with 3 mM EDTA in DPBS and, where indicated, were treated with TrypLE (Thermo Fisher Scientific) for 5-10 min at 37 °C and filtered through a 40-μm cell strainer to generate single cells. Cells were plated at a density of 2,000 cells per 40 μl of Matrigel and incubated at 37 °C for 30 minutes until domes solidified. Organoids were cultured in HISC (Advanced DMEM/F12 (AdDF12; Thermo Fisher Scientific), GlutaMAX (2 mM, Thermo Fisher Scientific), HEPES (10 mM, Thermo Fisher Scientific), N-Acetyl-L-cysteine (1 mM, Sigma-Aldrich), B27 Supplement with Vitamin A (Thermo Fisher Scientific), Primocin (100 μg/ml, InvivoGen), EGF (50 ng/ml, Peprotech), Noggin (100 ng/ml, Peprotech), A8301 (500 nM, Sigma), FGF2 (50 ng/ml, Peprotech), IGF-I (100 ng/ml, Peprotech)) or IGFF (HISC without EGF, Noggin, A-8301, FGF2, IGF-I) media supplemented with Y-27632. Media was changed every 3-4 days. Organoids were harvested at 7 days for downstream assays. Where indicated, small intact organoids (2-3 days post passage) were treated with 250 nM irinotecan added to HISC media for 7 days and then harvested for downstream assays. Lentiviral transduction of organoids was performed as described^99^. For single cell RNA sequencing of organoids, organoids were harvested from Matrigel with 3 mM EDTA in DPBS, dissociated to single cells with Accutase (Sigma-Aldrich), for 30-45 minutes at room temperature, washed with IGFF media and processed as described above.

#### Inducible knockdown

For doxycycline inducible PROX1 knockdown experiments, de novo 97-mer mirE shRNA sequences were synthesized (IDT Ultramers) and PCR amplified using the primers miRE-Xho-fw (5′- TGAACTCGAGAAGGTATATTGCTGTTGACAGTGAGCG-3′) and miRE-EcoOligo-rev (5′- TCTCGAATTCTAGCCCCTTGAAGTCCGAGGCAGTAGGC-3′) as described^100^. Sequences (shPROX1-1: TGCTGTTGACAGTGAGCGCGAGGACCAAGATGTCATCTCATAGTGAAGCCACAGATGTATGAGATGACATCTTGGTCCTCATGCCTACTGCCTCGGA; shPROX1-2: TGCTGTTGACAGTGAGCGCCCCCGAGAAAGTTAC AGAGAATAGTGAAGCCACAGATGTATTCTCTGTAACTTTCTCGGGGATGCCTACTGCCTCGGA) were cloned into the LT3GEPIR backbone^100^ (Addgene; 111177) and used to generate lentiviral particles to transduce into organoids as described^99^. The original plasmid containing an shRNA sequence to Renilla 306 luciferase (shRen.713) was used as a control. Transduced organoids were selected using HISC media supplemented with 2 μg/ml puromycin for 7 days. For inducible knockdown experiments, organoids were dissociated into single cells, plated at a density of 2,000 cells per 40 μL Matrigel and maintained in HISC or IGFF media supplemented with 2ug/ml doxycycline (Fisher Scientific) unless otherwise specified for 7 days before downstream assays. For organoid initiation and outgrowth assays, organoids containing inducible PROX1 or control shRNA cultured in HISC media supplemented with 2ug/ml doxycycline for 7 days were dissociated into single cells, and plated at a density of 2,000 cells per 40 ul matrigel in HISC media without Y-27632 supplemented 2ug/ml doxycycline, and imaged at 7 days (BioTek).

#### Western blotting

Approximately 4,000 organoids (3-4 million cells) were recovered from Matrigel using 3 mM EDTA in DPBS, washed, centrifuged (200*g*, 5 min, 4°C) and lysed with 1x RIPA buffer supplemented with PPI (1:100, Sigma- Aldrich, 04693132001), and benzonase (1:100, Fisher Scientific, 70-664-3) on ice for 30 minutes. Protein concentration was determined by Pierce BCA assay (Thermo Scientific, 23227). 10 ug protein per sample was separated by SDS-PAGE on Bis-Tris polyacrylamide gels (Fisher Scientific, NW04120BOX), transferred to activated PVDF membranes (Millipore, IPFL00010) and blocked in 3% BSA-TBST solution for 30 minutes. Membranes were incubated overnight at 4°C with the following antibodies: mouse anti-Beta-actin (1:1000, Thermo Fisher Scientific, AM4302) and rabbit anti-PROX1 (1:1000, Abcam, ab199359), then secondary antibody incubation with 488 anti-mouse and 680 anti-rabbit secondary antibodies (1:5000, LI-COR Biosciences, 1 hour, room temperature) prior to imaging (Odyssey CLx). Western blots were quantified using ImageJ software(v1.53t) ^101^.

#### Organoid whole mount immunofluorescence

Whole organoids were plated in 10% Matrigel and HISC media on chamber slides with removable wells (Thermo Fisher Scientific, 177380), and incubated at 37°C for 1 hour. Organoids were then fixed in 4% paraformaldehyde in PME solution for 10 minutes, washed twice with IF buffer (0.2% Triton X-100, 0.05% Tween in DPBS), permeabilized (0.5% Triton X-100 in DPBS for 10 minutes, room temperature), blocked in 10% Normal Goat Serum (Thermo Fisher Scientific, 50062Z) for 30 minutes prior to overnight primary antibody incubation with 1:500 rabbit anti-PROX1 (Abcam, ab199359) in 10% Normal Goat Serum at 4°C. Organoids were incubated with 1:400 of 594 anti-Rabbit antibody (Invitrogen, A11012) for 1 hour at room temperature, washed 3 times with IF buffer and 1 time with cold DPBS. Wells were removed and samples mounted using medium containing DAPI (Novus Biologics, H-1200-NB). Imaging was performed on a Zeiss Axio Imager 2 with Apotome structured illumination microscope. PROX1 expression was quantified using the CellProfiler (v4.2.5) software^102^.

#### Reverse transcription and qPCR

Total RNA was extracted from organoids, crypts or dissociated cells with the RNeasy Mini kit (Qiagen). Total RNA (3–4 μg) was used to prepare cDNA with the Transcriptor First-Strand cDNA synthesis kit (Roche). qPCR was performed with TaqMan gene expression assay primers (Thermo Fisher Scientific; *PROX1*: Hs00896293_m1; *GAPDH:* Hs02758991_g1; *ELF5*: Hs01063022_m1; *KRT23*: Hs00210096_m1; Hs00167549_m1; *TFF3*: Hs00902278_m1; *FABP1*: Hs00155026_m1), Relative expression was quantified using the ddCT method, normalized to the expression of *GAPDH* on a QuantStudio 6 and 7 Pro real-time PCR system (Applied Biosystems).

#### Orthotopic xenograft experiments

All animal experiments were performed in accordance with protocols approved by the Memorial Sloan Kettering Cancer Center Institutional Animal Care and Use Committee (IACUC). NSG (NOD.Cg- Prkdc^scid^IL2rg^tm1Wjl^/SzJ; stock 005557) mice were obtained from the Jackson Laboratory and were transplanted at 6 weeks of age^103^. For orthotopic cecal injections, KG146P or KG146Li organoids were transduced with pLenti-PGK-Akaluc (AkaLuc) or pLenti-PGK-tdTomato-Akaluc (TdT-AkaLuc) virus (subcloned from pLenti-PGK-Venus-Akaluc(neo)) and HR180-LGR5-iCT plasmids through Gibson assembly (Addgene 124701; 129094). For cecal injections, 200,000 cells from each organoid line were mixed with 50% Matrigel in 10 μl of DPBS and injected into the cecal submucosa of NSG mice. For hepatic injections, 500,000 cells from each organoid line were mixed with 50% Matrigel in 70 μl of DPBS and injected under the liver capsule of NSG mice. Weekly bioluminescence imaging was performed on an IVIS Spectrum Xenogen instrument (Caliper Life Sciences) and analyzed with Living Image software v2.50. Experimental group sizes were practically determined on the basis of cages (five mice per cage), with *n* ≥ 5 mice per group. Animals were euthanized at the humane endpoint, and tissues harvested for downstream assays. Where needed, tissues were fixed in 4% paraformaldehyde prior to downstream assays.

## Data and Code Availability

All raw data and code are available by request and will be uploaded to public databases prior to publication.

---

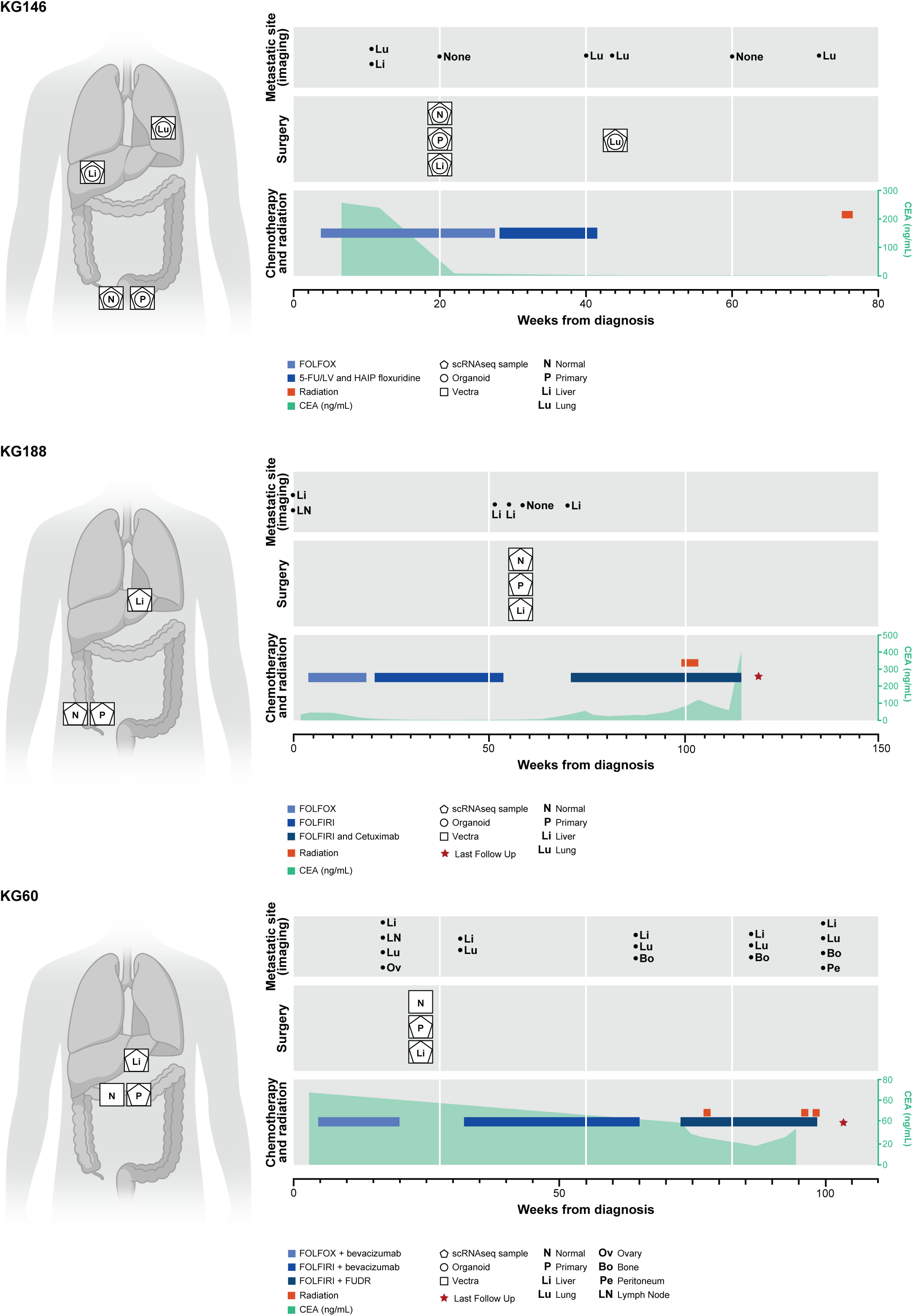
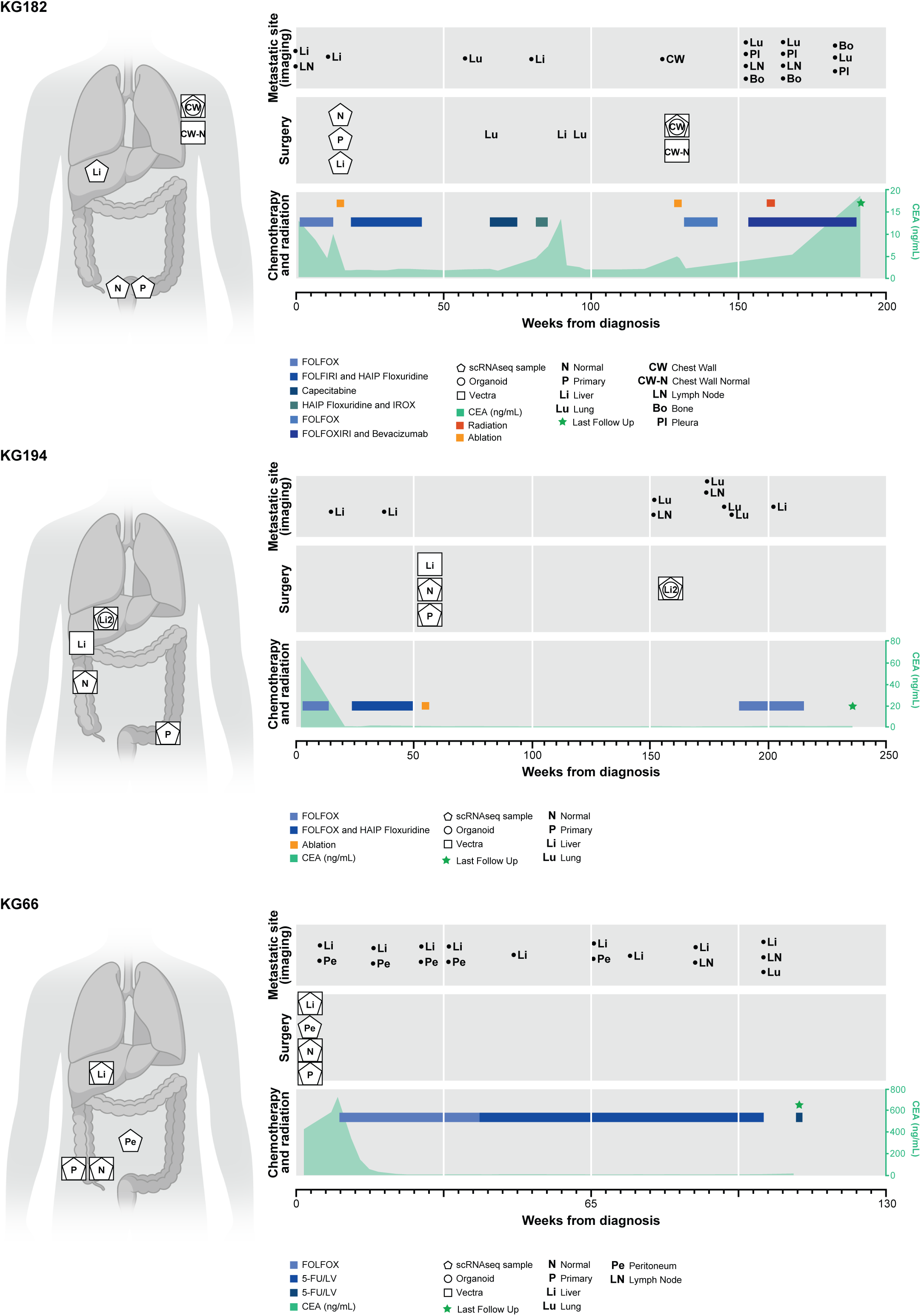
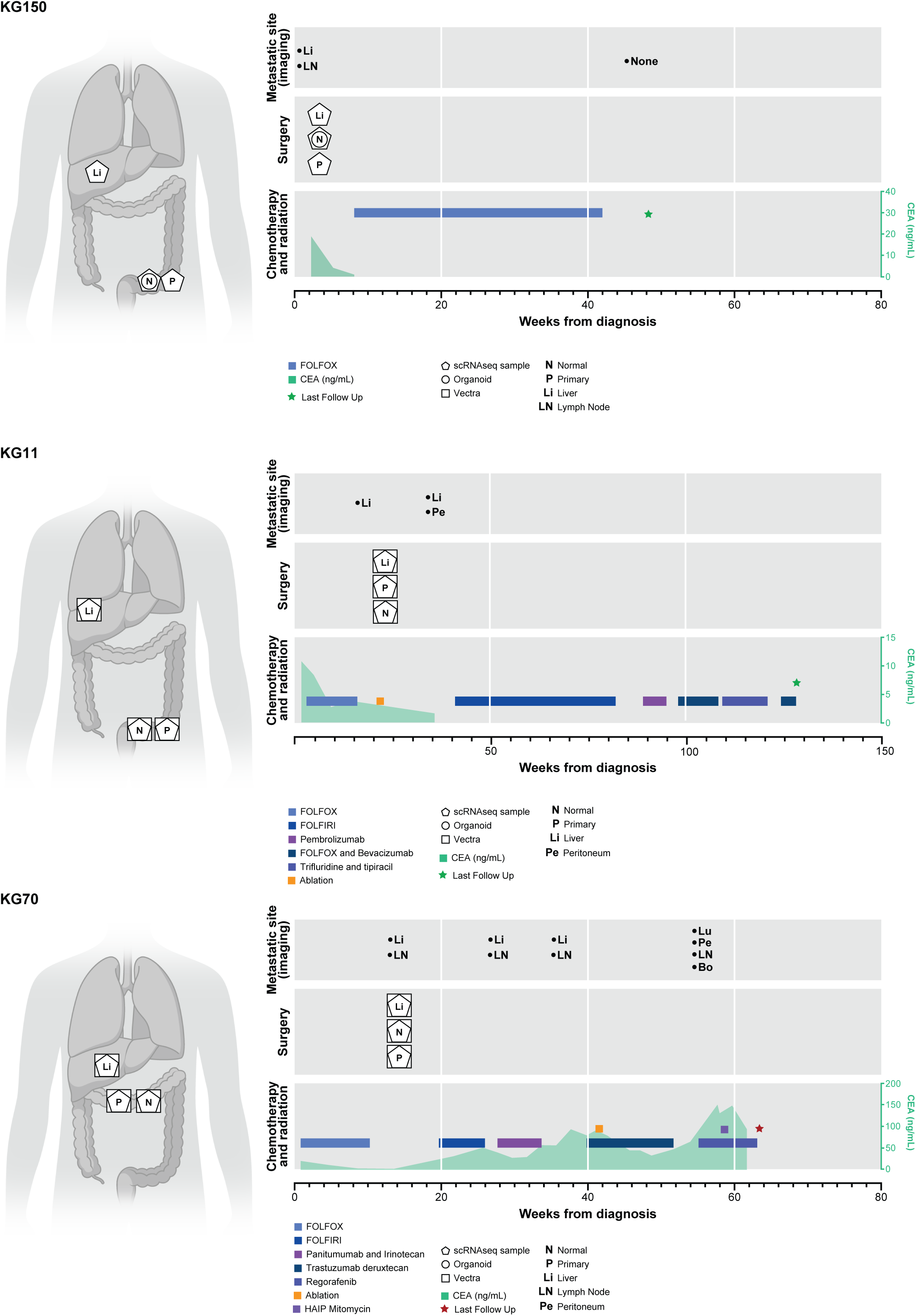
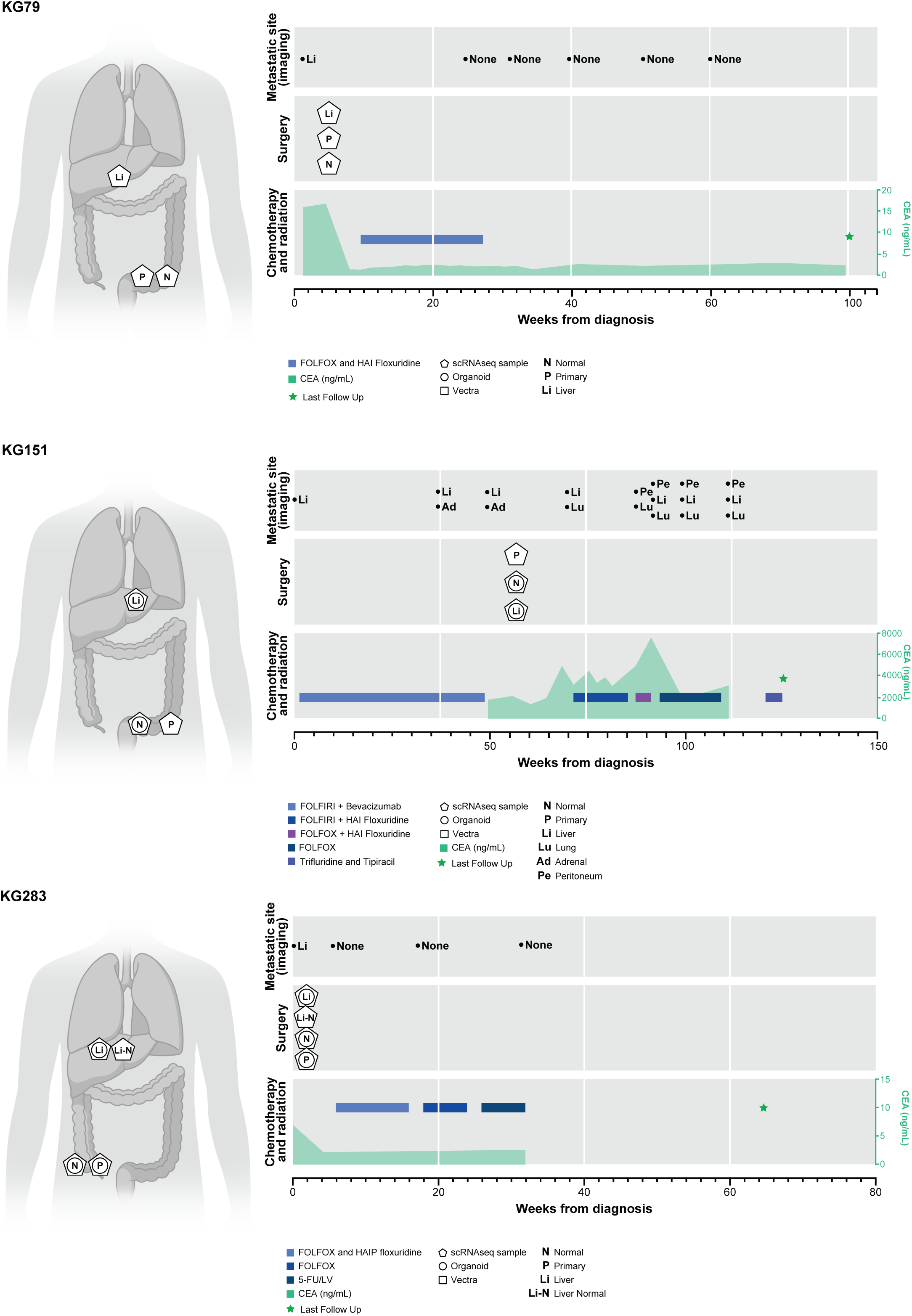
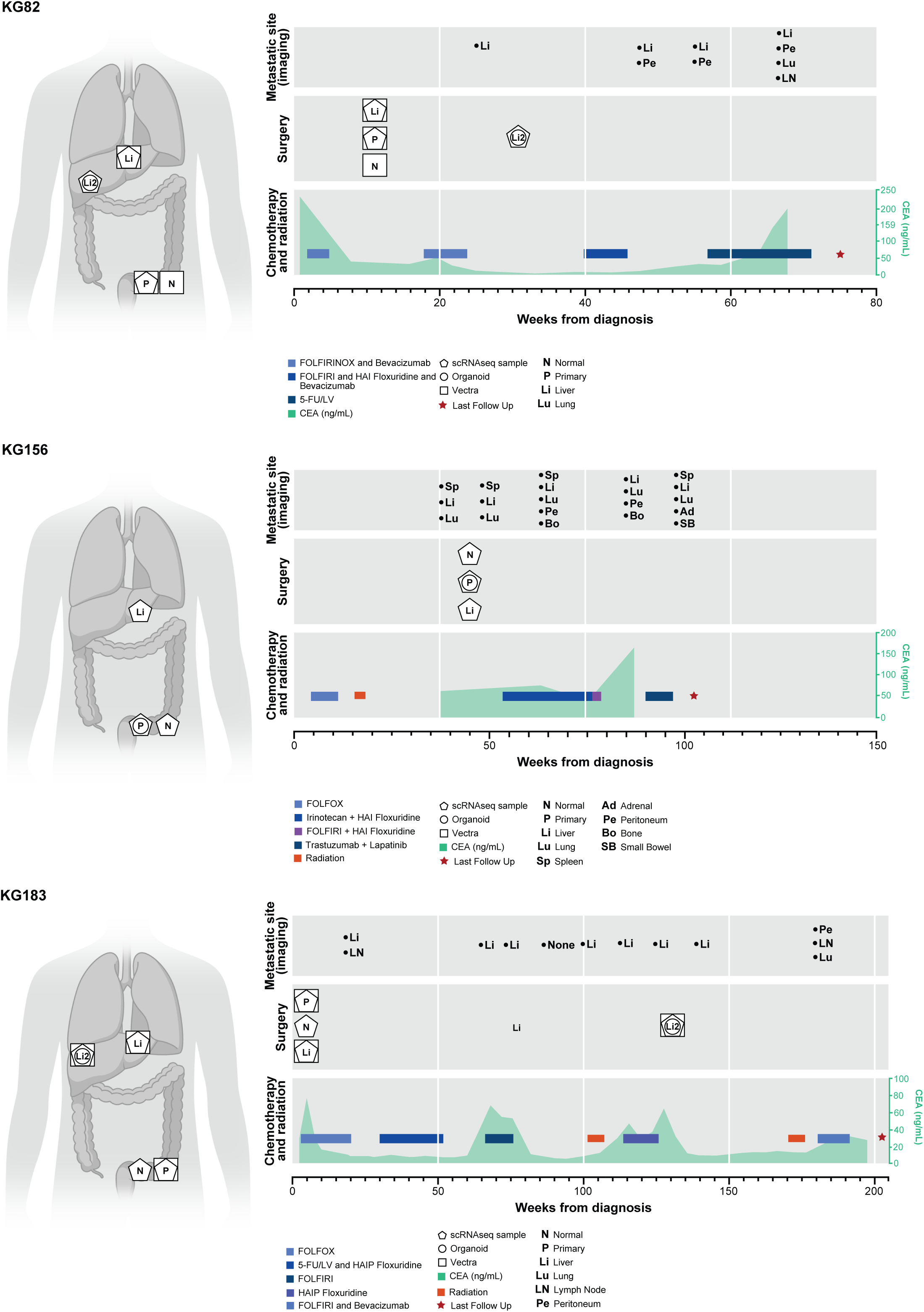
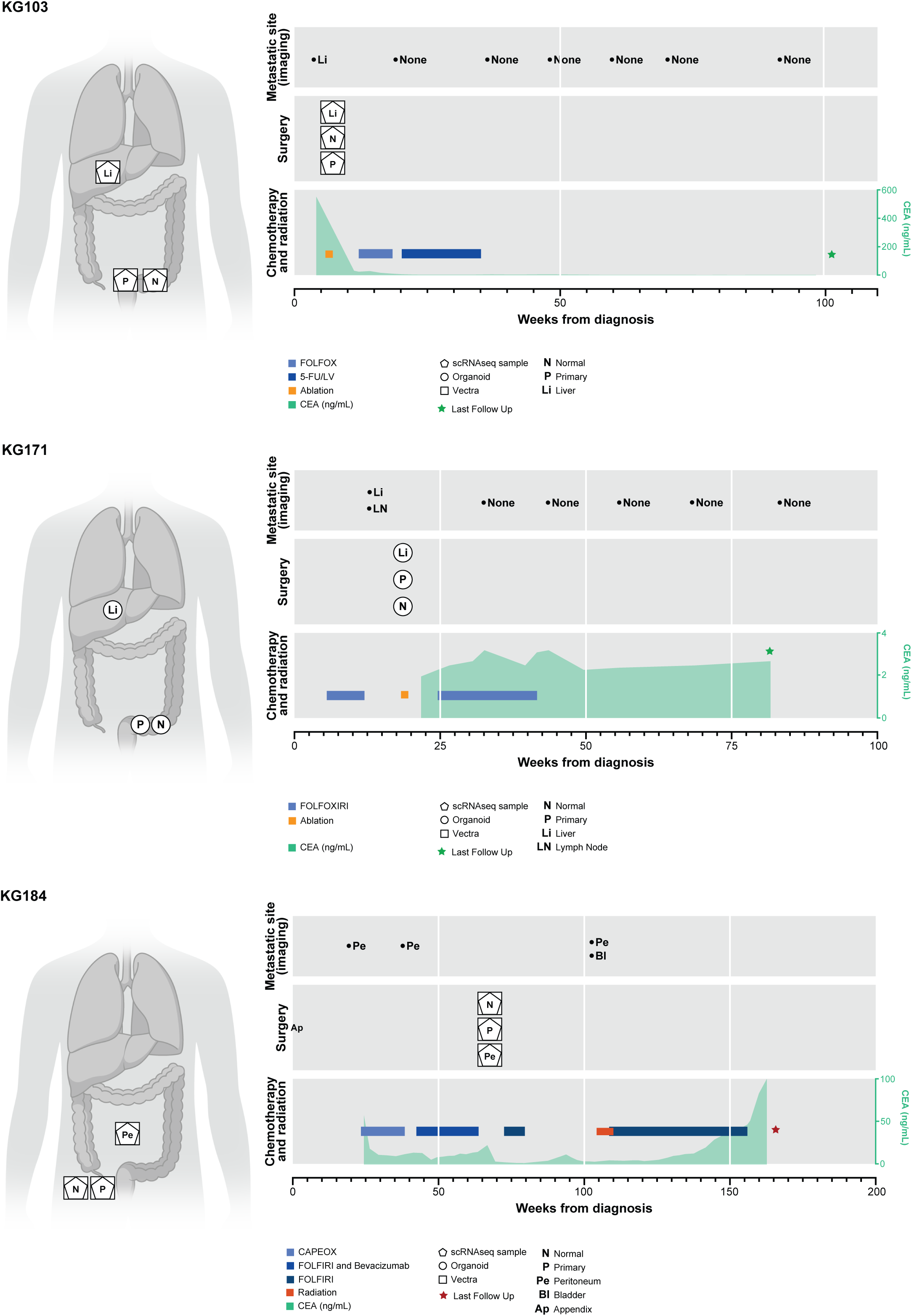
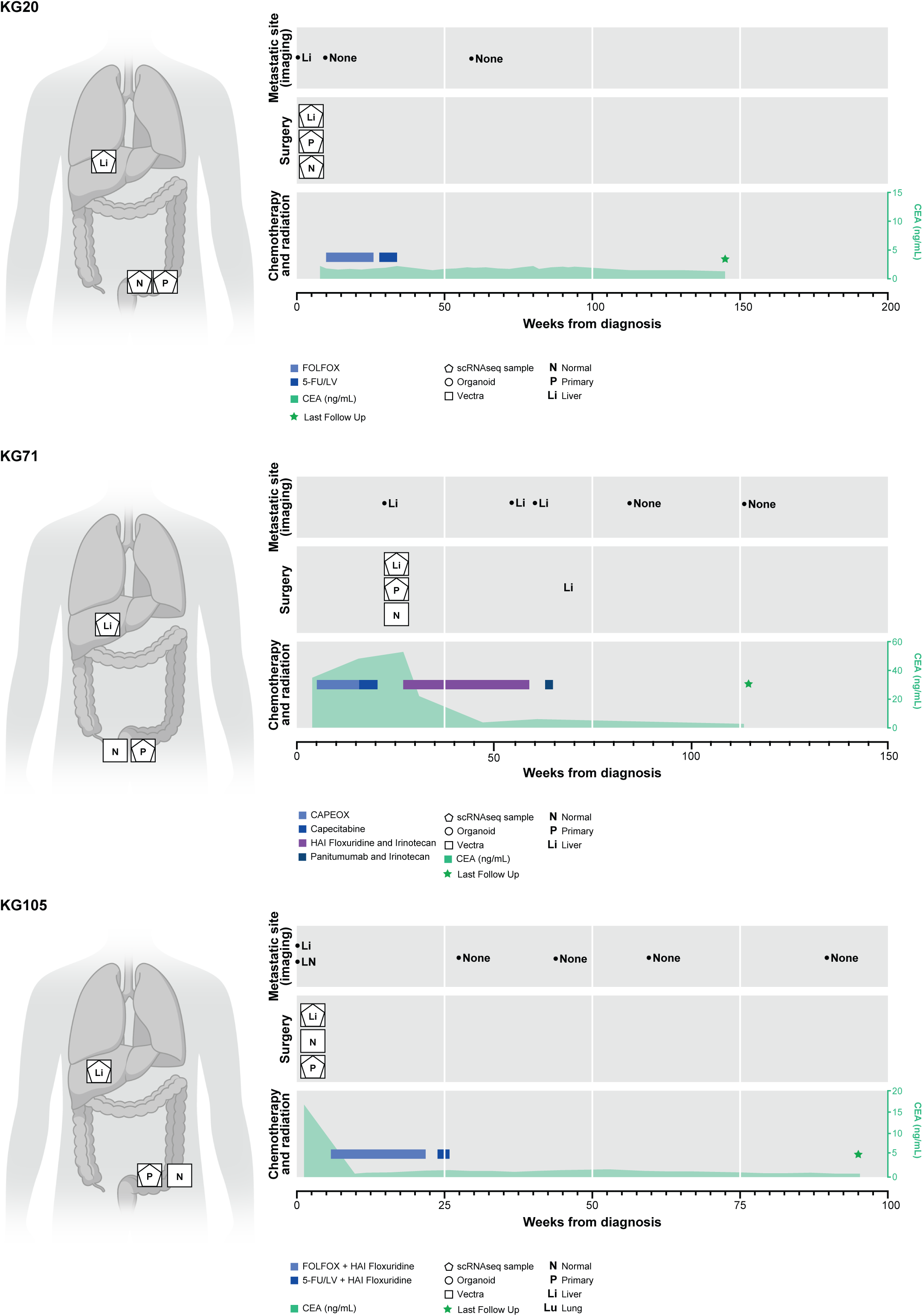
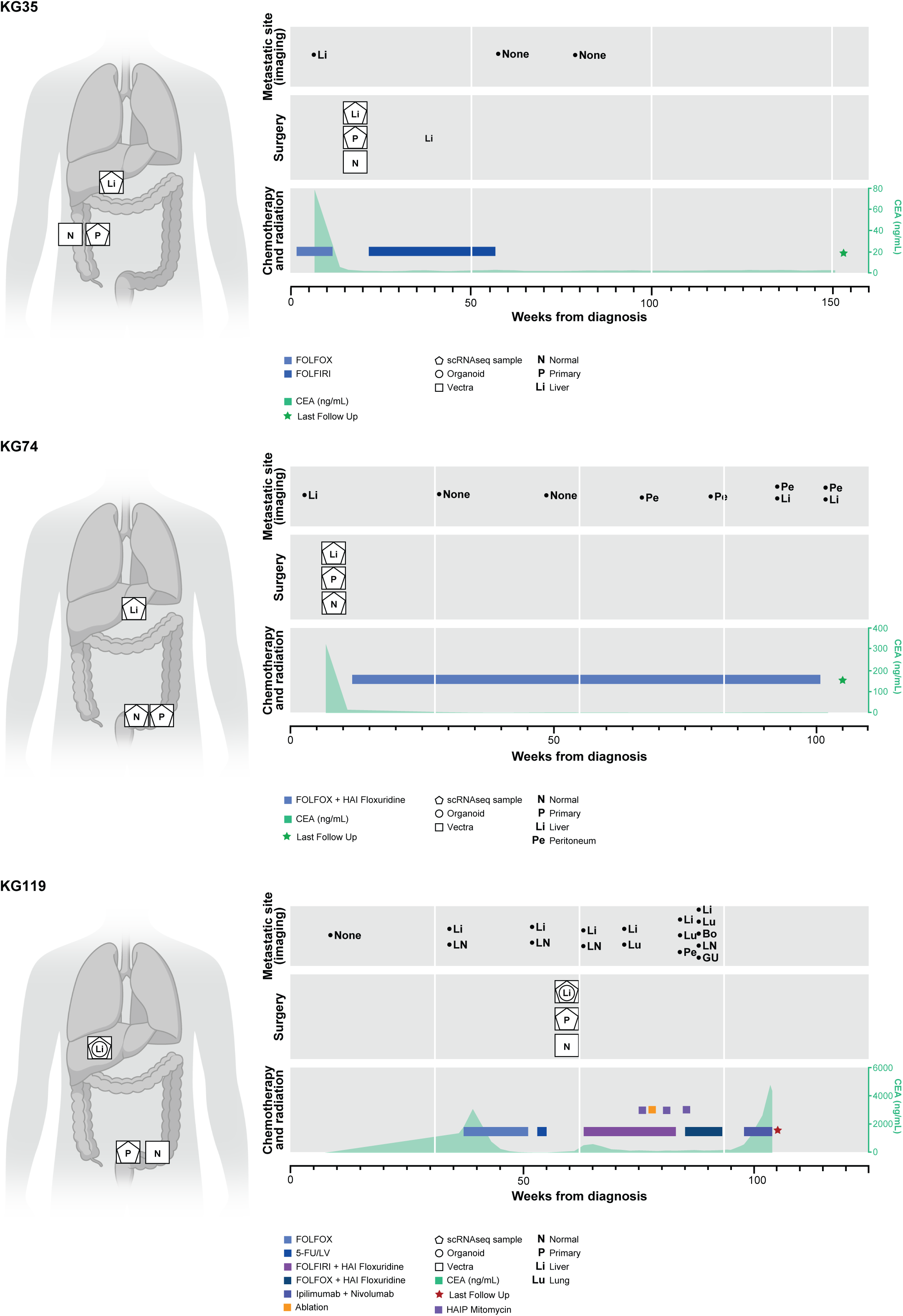
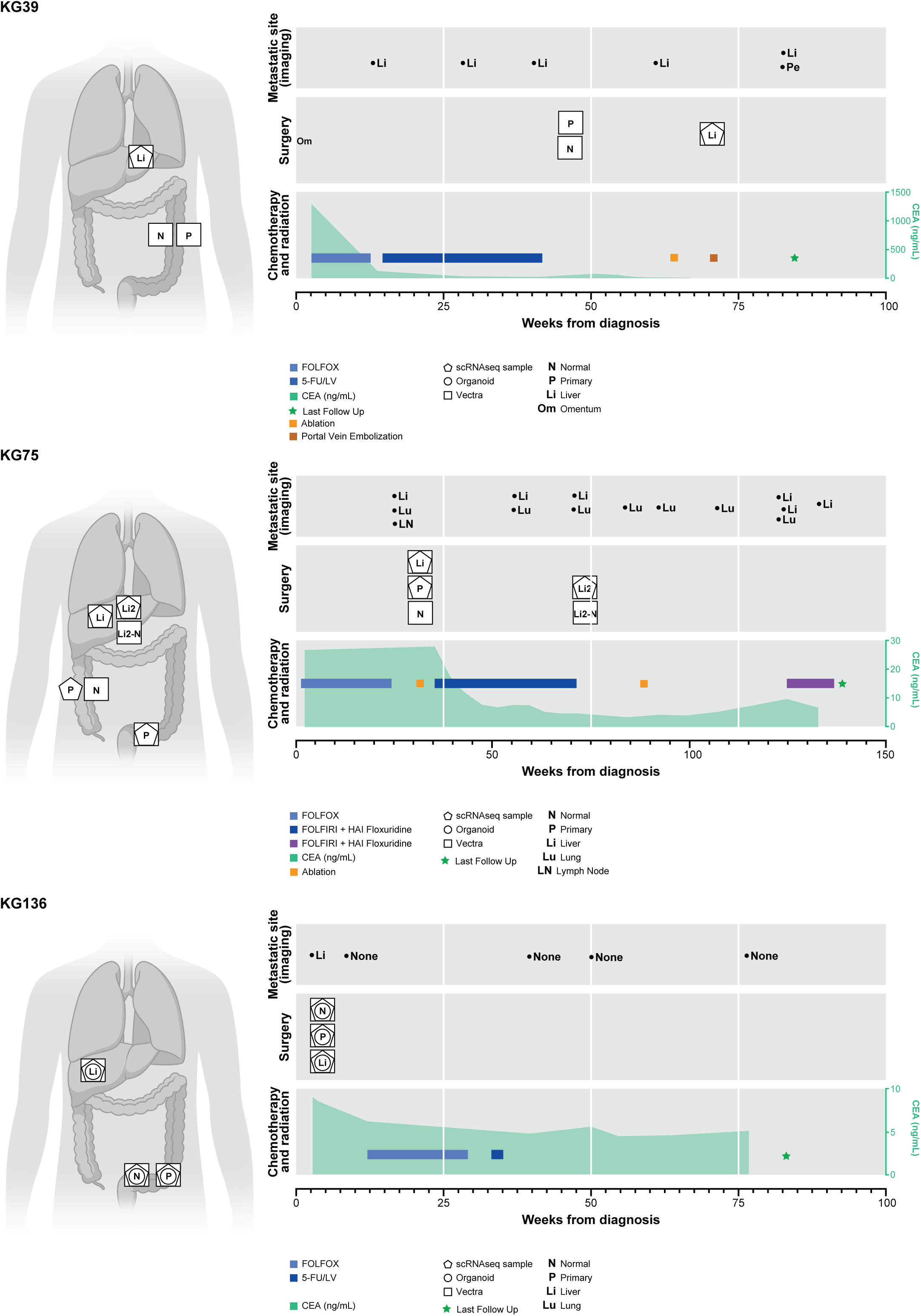
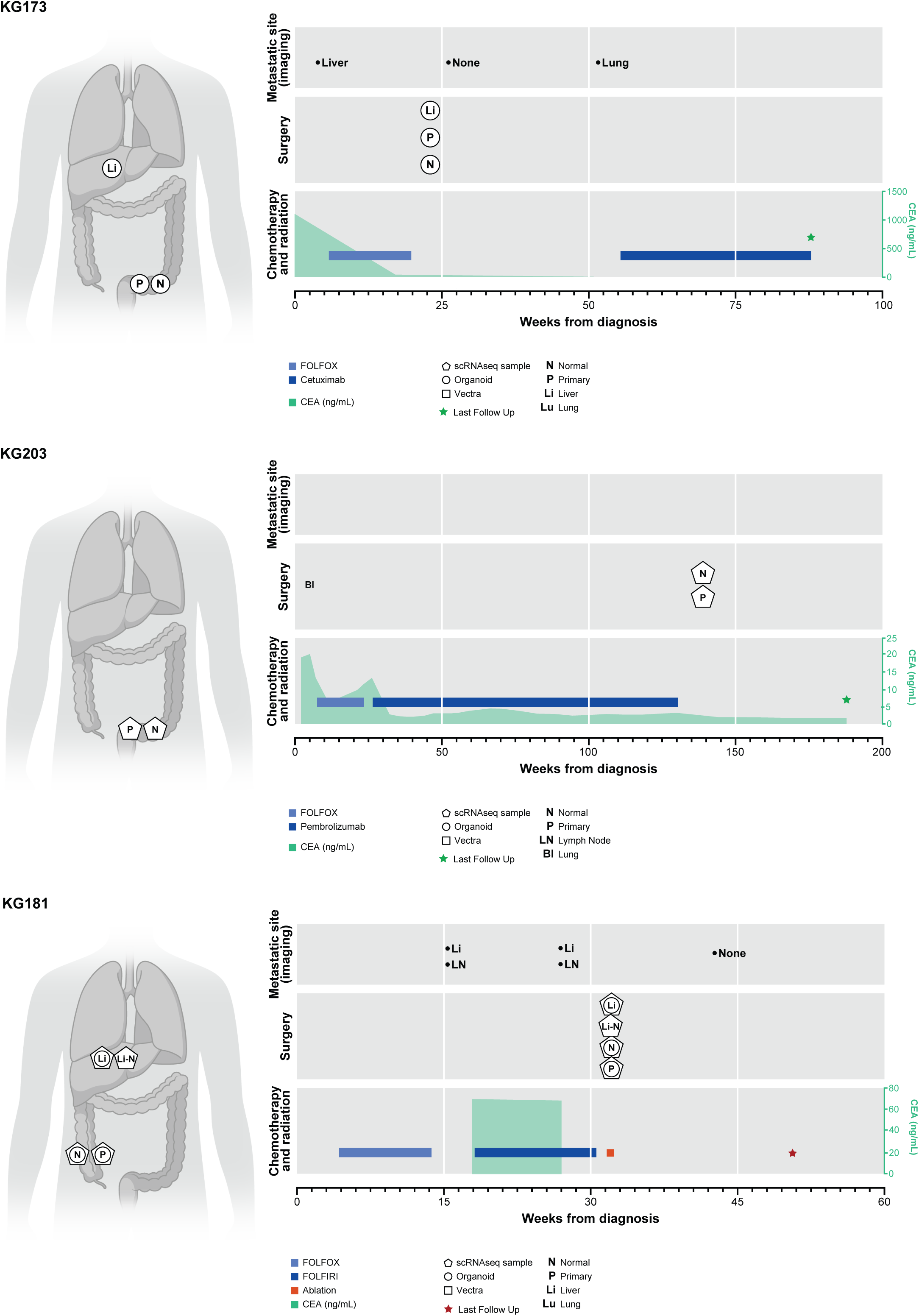
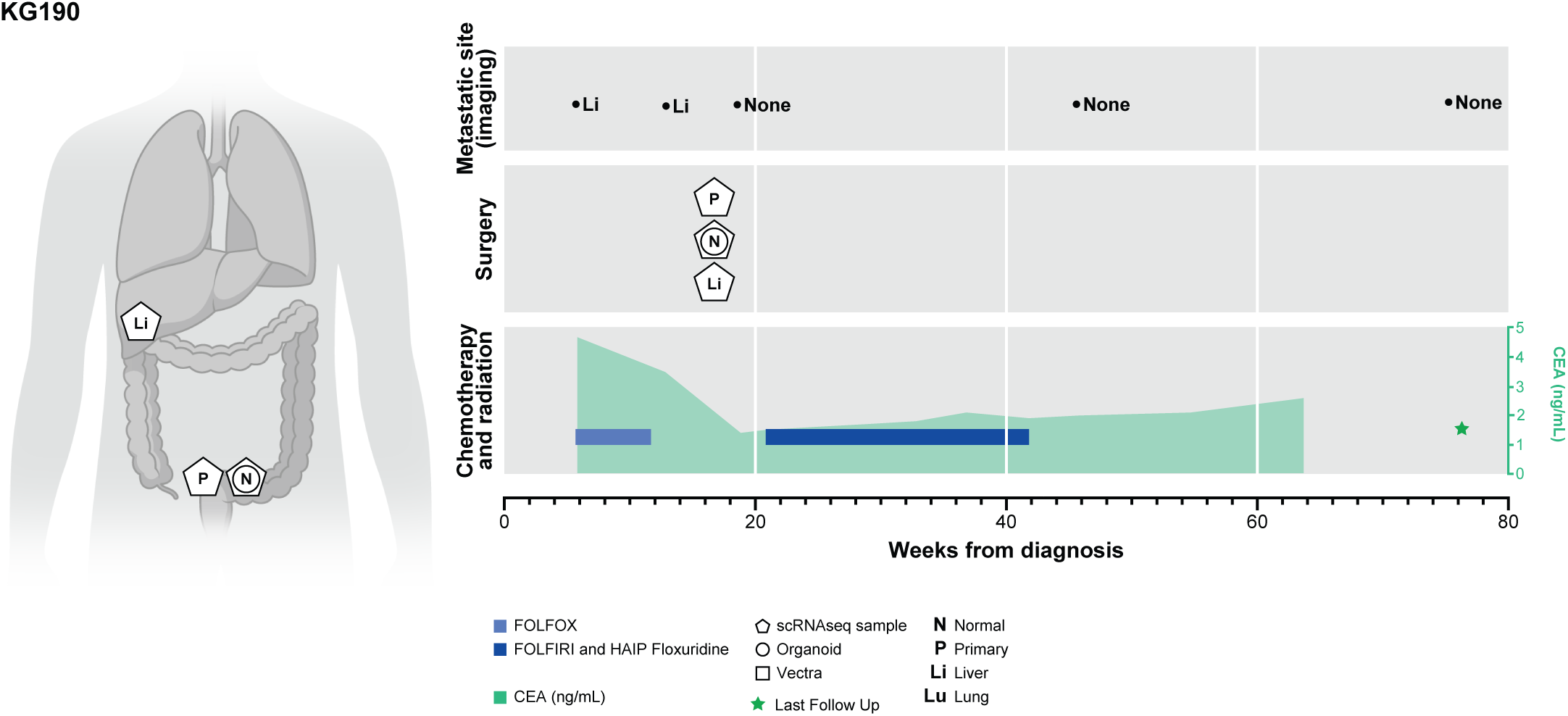

## Notes

### Competing Interest Statement

J.S. is a consultant for Paige AI. D.P. is on the scientific advisory board of Insitro. K.G. is a consultant for Seres Therapeutics and is an inventor on patents related to targeting metastasis.

